# Portable transcranial therapeutic ultrasound enhances targeted gene delivery for Parkinson’s disease: from rodent models to non-human primates

**DOI:** 10.1101/2025.03.02.639893

**Authors:** Alec J. Batts, Robin Ji, Sua Bae, Fotios N. Tsitsos, Sergio Jiménez-Gambín, Nancy Kwon, Samantha L. Gorman, Deny Tsakri, Rebecca L. Noel, Jonas Bendig, Daniella A. Jimenez, Melody DiBenedetto, Sofia A. Del Castillo, Filimon B. Keleta, James Caicedo, Alexander Romanov, Colleen T. Curley, Yulia Dzhashiashvili, Greglynn D. Walton-Gibbs, Bradley S. Hollidge, Serge Przedborski, Elizabeth Ramsburg, Olivier Danos, Vincent P. Ferrera, Esteban A. Engel, Jared B. Smith, Elisa E. Konofagou

## Abstract

Gene therapy for neurodegenerative diseases faces significant challenges due to the blood-brain barrier (BBB), which limits drug delivery to the central nervous system (CNS). While clinical trials for Parkinson’s disease (PD) have progressed, administration of vectors expressing enzymatic or neurotrophic factor transgenes have required extensive optimization of the delivery method to achieve potentially therapeutic levels of transgene expression. Focused ultrasound (FUS) combined with microbubbles has emerged as a promising non-invasive strategy to transiently open the BBB for targeted gene delivery via viral nanocarriers including recombinant adeno-associated viruses (AAVs). However, key factors influencing FUS-mediated AAV delivery, including dose distribution and therapeutic efficacy, remain underexplored in non-human primates (NHPs). Here, we evaluated the feasibility of AAV9-CAG-GFP delivery using two portable therapeutic ultrasound modalities: ultrasound-guided, spherically-focused FUS (USgFUS) and a novel low-frequency linear array configuration for imaging and therapy called theranostic ultrasound (ThUS). In mice, FUS-sonicated regions exhibited a 25-fold increase in AAV9 biodistribution compared to systemic injection alone. Extending this approach to NHPs, we observed up to a 200-fold increase in AAV9 DNA in treated brain regions, including PD-relevant structures. In assessing the translational therapeutic potential of this technique, ThUS-mediated AAV9-hSyn-hNTRN (human neurturin) delivery in a toxin mouse model of PD facilitated the rescue of up to 80% and 75% of degenerated dopaminergic neurons in the substantia nigra and striatum, respectively. These findings demonstrate that portable ultrasound technologies can non-invasively enhance AAV9 delivery to targeted brain regions in both mice and NHPs relative to what can be achieved with intravenous (IV) delivery of the same capsid alone. With further development, these approaches may offer a clinically viable, non-invasive alternative for gene therapy in neurodegenerative diseases.

**One sentence summary:** BBB opening with portable therapeutic ultrasound non-invasively increased viral gene delivery to the brain after systemic AAV vector administration in mice and rhesus macaques.

## INTRODUCTION

Neurodegenerative diseases are characterized by the progressive loss of neurons and their associated functions. Given that over 15% of the global population is estimated to be afflicted with a neurodegenerative disease, and their incidence is expected to increase rapidly over the next several decades (*1, 2*), there is an urgent need for more effective treatments. Gene therapy, a technique which modulates gene expression for therapeutic benefit, has advanced substantially since the first human trial in 1990. Strategies include gene overexpression or miRNA-mediated gene silencing to target disease mechanisms. These approaches are particularly relevant for neurodegenerative diseases, which can be genetic, as in Huntington’s disease (HD), or result from genetic and acquired factors, as in Alzheimer’s (AD) and Parkinson’s diseases (PD).

Adeno-associated viruses (AAV) have emerged as one of the preferred gene delivery vectors for neurodegenerative diseases due to their relatively low immunogenicity and genotoxicity (*3*). Various administration routes are being explored to optimize therapeutic efficacy while mitigating risks, each presenting distinct advantages and challenges. Intraparenchymal (IP) delivery, as exemplified by the recent approval of Upstaza—the first FDA-approved AAV-based gene therapy administered directly to the brain for treating aromatic L-amino acid decarboxylase (AADC) deficiency—allows targeted gene delivery with reduced systemic exposure and lower doses but requires an invasive neurosurgical procedure. In diseases with multiple affected brain regions or widespread pathology, this approach may necessitate prolonged surgical times to reach multiple targets, making it less practical for certain conditions. Alternative approaches, such as intracerebroventricular (ICV) or intrathecal (IT) administration into the cerebrospinal fluid (CSF), may facilitate broader CNS transduction in diseases with diffuse pathology, with ICV requiring an invasive burr hole procedure (like IP delivery) for direct ventricular access, and IT utilizing a less invasive lumbar puncture to deliver vectors to the CSF. However, variability in clinical outcomes with these methods highlights the necessity of precise preclinical assessments to optimize transgene biodistribution in patients (*4, 5*). Systemic delivery via intravenous (IV) injection circumvents the need for neurosurgical intervention but requires engineered blood-brain barrier (BBB)-penetrating capsids and high vector doses (*5*), increasing the risk of peripheral toxicity, including fatal liver damage (*6*). While intraparenchymal administration offers advantages such as localized delivery and reduced systemic exposure (*7–12*), limitations of the technique for treating PD specifically (*13–15*) motivates the need for novel AAV administration methods such as focused ultrasound (FUS) which may provide non-invasive alternatives to enhance gene transfer to the CNS.

FUS is a well-established technique to non-invasively and focally circumvent the BBB and therefore permits reduced IV injection doses. When used in conjunction with systemically-administered microbubbles, FUS has proven to elicit transient, safe, reversible and focal opening of the BBB, conferring therapeutic benefit in multiple mechanisms involved in neurodegenerative disorders both pre-clinically and clinically (*16–24*). Notably, FUS enables delivery to multiple brain regions in a shorter timeframe compared to the corresponding surgical procedures and exhibits the potential to target larger areas, such as the neocortex. While most clinical trials have utilized magnetic-resonance-guided FUS (MRgFUS) to achieve BBB opening, which was safe and effective in clinical trials for BBB opening in neurodegenerative diseases and cancer (*23, 25–30*), MRgFUS is inherently costly and not widely accessible to patients since a dedicated MRI scanner is required for the procedure. Additionally, MRgFUS procedures require patients to remain in the MRI scanner for prolonged periods of time, significantly reducing patient comfort. Our group has developed cost-effective, fast, and portable ultrasound-guided FUS (USgFUS) devices that enable BBB opening outside of an MRI scanner through the use of neuro-navigation technology. USgFUS modalities also leverage ultrasound guidance to monitor microbubble activity during BBB opening (*31–36*), most often with a separate receiving transducer in addition to the FUS therapy transducer. Advantageously, this FUS-induced intravascular microbubble activity correlates with both the volume of contrast-enhancement on T_1_-weighted MRI, the gold-standard in confirming BBB opening following FUS, and the amount of AAV transgene expression after BBB opening (*31*). Moreover, ultrasound guidance through B-mode and harmonic imaging of the skull enables precise and non-invasive targeting of specific brain structures without the need for intraprocedural MRI (*37*). A novel single-transducer modality for USgFUS, called theranostic ultrasound (ThUS), was also developed for an even more favorable tradeoff between flexibility and portability, and exhibits other advantages over traditional USgFUS configurations including integrated high-resolution microbubble cavitation imaging, accelerated BBB repair, and simultaneous multi-focal BBB opening (*31, 38–40*). Furthermore, this configuration is operated by existing diagnostic ultrasound scanner hardware available in the clinic. Therefore, BBB opening with USgFUS configurations including ThUS could provide a more accessible and cost-effective alternative to BBB opening procedures, currently in the clinic, that are performed with intraoperative MRI.

In terms of drug delivery, FUS enables reduction of the systemic dose because the increase in BBB permeability temporarily allows increased drug transport into the brain. Our group and others have demonstrated the tolerability, efficacy, delivery efficiency, and therapeutic benefit of FUS-mediated AAV delivery in both healthy and Parkinsonian animal models (*31, 41–45*). AAV9 has specifically been selected for many pre-clinical AAV delivery studies employing FUS and intravenous injection due to its natural CNS tropism and ability to cross the BBB, though in very limited concentrations (*46*). To date, only two published studies have reported increased focal AAV delivery following BBB opening with FUS in non-human primates (NHP), one conducted with MRgFUS in rhesus macaques (*43*), and the other with FUS in marmosets (*44*). While these studies demonstrated that FUS-mediated BBB opening for targeted AAV delivery is possible in NHP, there still remains an unconfirmed quantitative relationship between systemically-administered AAV dose and targeted gene expression within clinically relevant targets in the NHP brain, as well as an understudied relationship between FUS pulsing parameters and resulting transduction efficacy in NHP.

We designed our study presented herein to address the following: i) investigate the relationship between systemic AAV dose and resulting biodistribution after BBB opening in multiple brain regions, ii) evaluate the FUS parametric space for AAV delivery, iii) establish safety considerations, and iv) capitalize on the advantages posed by USgFUS. We conducted a dose escalation study in mice to determine a relationship between systemic dose and AAV9 transduction in brain regions opened with FUS. We then evaluated the feasibility for AAV9 delivery in NHPs with both a clinical USgFUS configuration and a low-frequency ThUS linear array configuration to model clinical translation of gene delivery for neurodegenerative disease facilitated by more accessible therapeutic ultrasound systems. Finally, we provide insight into the future opportunities for therapeutic ultrasound in treating neurodegenerative disorders through a study conducting neurotrophic factor (NTF) gene delivery with ThUS in a mouse model that recapitulates neurodegeneration in early PD (*47*). The results of these studies elucidate the advantages and limitations of potential USgFUS configurations for AAV-mediated gene delivery across the BBB, while revealing the potential for therapeutic efficacy of non-invasive transcranial gene delivery with accessible USgFUS configurations in neurodegenerative disorders.

## RESULTS

### Dose escalation of systemically-administered AAV9 reveals dose-dependent transduction following FUS-mediated BBB opening in C57BL6/J wild-type mice

To inform the AAV dosing for the following NHP study with USgFUS, we first conducted an AAV9 dose escalation study in mice. The following 3 doses of the AAV9-CAG-GFP construct were intravenously injected before BBB opening with FUS targeting the hippocampus (n=5 mice per group): 1.0e10 gc/mouse, 1.0e11 gc/mouse, and 5.0e11 gc/mouse. Two additional groups were included: one group of 5 mice received an intravenous injection of 5.0e11 gc/mouse without FUS BBB opening, while the last group received neither AAV nor FUS, but still underwent anesthesia.

The average BBB opening volume induced with FUS in groups receiving FUS and IV injection of AAV was 88.06 ± 23.17 mm^3^, quantified from contrast-enhanced T_1_-weighted MRI acquired 30 minutes post-FUS (Figure 2A). No statistically significant differences in BBB opening volume were observed between the AAV dose groups receiving FUS, indicating that groupwise differences in CNS transduction were driven by systemic dose, not variability in FUS sonication parameters or induced BBB opening volume (Figure 2B). Mice were then survived for 21 days to allow for gene expression to occur before euthanasia. After euthanasia and tissue processing, ddPCR revealed statistically significant increases in AAV DNA within the hippocampus between groups; a 25-fold increase in DNA was detected in the hippocampi of mice which received the highest AAV dose and FUS BBB opening relative to mice receiving only a systemic injection of the highest AAV dose with no FUS intervention (Figure 2C). Quantification of GFP luminance in brain sections allocated for IHC revealed a similar trend with statistically significant increases in GFP transgene expression facilitated by FUS BBB opening with increasing AAV dose (Figure 2D). This is depicted qualitatively on microscopy images of representative sections from the FUS + low AAV dose group (Figure 2E), the FUS + medium AAV dose group (Figure 2F), and the FUS + high AAV dose group (Figure 2G). Finally, cell-type specificity of AAV transduction was evaluated across all dose groups with representative IHC presented of the high dose group in Figure 2H-L, where approximately 80% of transduced cells were astrocytes, while the remaining 20% were neurons (Figure 2M).

**Figure 1:**
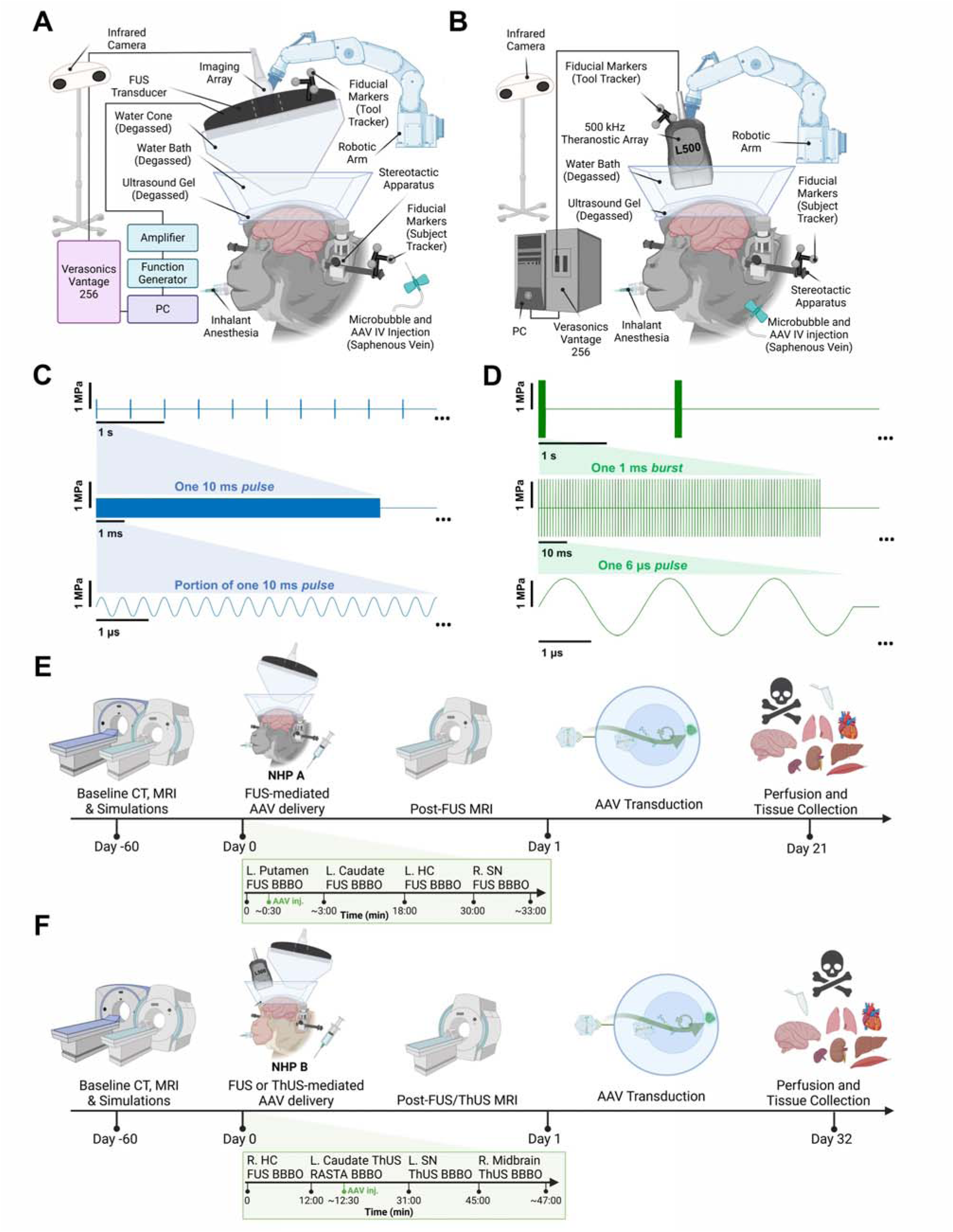
Experimental apparatus and NHP study overview. **A)** FUS BBB opening experimental configuration with neuronavigation guidance. **B)** ThUS BBB opening experimental configuration with neuronavigation guidance. **C)** Long-pulse FUS sequence diagram. **D)** Short-pulse ThUS sequence diagram. **E)** Timeline for FUS-AAV delivery with NHP A. **F)** Timeline for FUS and ThUS-AAV delivery with NHP B.

**Figure 2:**
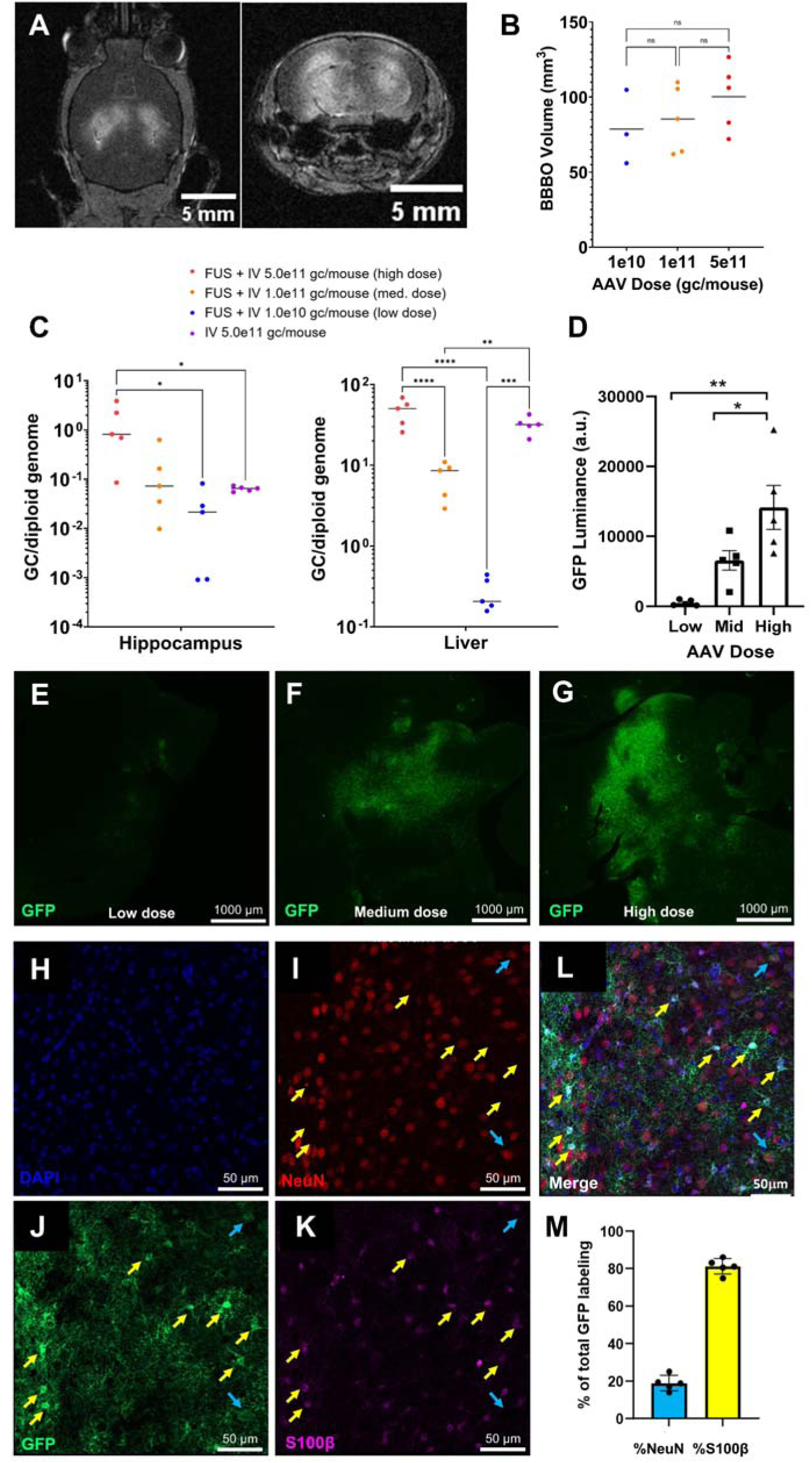
AAV dose escalation increases FUS-facilitated gene delivery to the mouse CNS. **A)** Representative axial (left) and coronal (right) T_1_-weighted MRI depicting bilateral hippocampal BBB opening. **B)** Groupwise comparisons of BBB opening volume across AAV dose groups. **C)** Liver and hippocampal genome copies per cell derived from ddPCR. **D)** Luminance of GFP expression in IHC microscopy images. Representative GFP expression from IHC in **E)** 1.0e10 gc (low dose), **F)** 1.0e11 gc (med. dose) and **G)** 5.0e11 gc (high dose) groups. Confocal imaging depicting representative proportion of astrocytic and neuronal GFP expression from the high dose group with individual **H)** DAPI, **I)** NeuN, **J)** GFP, **K)** S100β channels and **L)** merged channels. Yellow and blue arrows depict transduced astrocytes and neurons respectively. **M)** Percent of GFP expression in astrocytes versus neurons. Statistical significance in B-D determined by one-way ANOVA with Tukey’s multiple comparisons correction. *\*p*<0.05, *\*\*p*<0.01, *\*\*\*p*<0.001, *\*\*\*\*p*<0.0001.

While the intravenous route of administration yielded substantial liver transduction, which is particularly characteristic of AAV9, BBB opening with FUS reduced the ratio of liver to brain transduction from a 375-fold increase in gene copies within the liver over the brain at the highest systemic dose delivered without FUS intervention, to a 55-fold increase in gene copies within the liver over the brain at the same dose after FUS-mediated BBB opening (Figure 2C). Although the application of FUS to the brain did not significantly alter gene transduction in the liver, the significantly increased AAV DNA within the brain after FUS relative to that in peripheral organs highlights a significant opportunity for non-invasively reducing the required therapeutic dose of transgenic DNA with USgFUS.

### Systemic AAV9 injection and FUS-mediated BBB opening in NHP yields substantial improvement in targeted AAV9 transduction in the brain

The prior dose escalation study in mice informed the systemic dose used for the subsequent NHP studies. Specifically, the 5.0 × 10¹¹ gc/mouse high dose, administered to mice weighing an average of 25 g was scaled proportionally based on body weight to match the ∼13 kg rhesus macaques described herein, resulting in a corresponding total dose of approximately 2.6 × 10¹ gc per macaque. Given the increased microstructural complexity and thickness of the primate skull relative to the mouse skull, subject-specific estimates of the FUS focal volume and anticipated attenuation due to ultrasound propagation through the NHP skull were derived from acoustic wave propagation simulations and were used to apply the specific voltage to the FUS transducer needed to achieve the required in situ focal pressure for sufficient AAV delivery across the BBB, corresponding to an ultrasound mechanical index (MI) of 0.8 (Supplementary Figure 1, Supplementary Figure 2).

During the four FUS sonications, real-time cavitation doses calculated from the passive acoustic mapping (PAM) images of microbubble cavitation (overlaid atop regions of BBB opening denoted on MRI in Figure 3A-D) rose substantially approximately 30 seconds after MB injection into the saphenous vein indicating microbubble activity within the FUS focal volume. Consequently, contrast-enhanced T_1_-weighted MRIs acquired approximately 1 hour after the last sonication for each NHP confirmed successful FUS BBB opening along the FUS-targeted trajectories, as well as contrast-enhancement within each targeted region apart from the hippocampus, which did not exhibit radiological indications of BBB opening (Figure 3E-H). Quantification of contrast enhancement after subtraction of pre-sonication MRI revealed BBB opening volumes of 267.9 mm^3^, 358.6 mm^3^, and 227.4 mm^3^, for the combined putamen and caudate trajectories, the hippocampus trajectory, and the substantia nigra trajectory, respectively (Figure 3I), yielding a combined BBB opening volume of 853.9 mm^3^, or 0.854 cc over a ∼30-minute FUS procedure in NHP A (Supplementary Video 1). The single FUS sonication in NHP B yielded a contrast-enhanced volume of 886.52 mm^3^ along the FUS trajectory and sulci proximal to it (Figure 4C).

**Figure 3:**
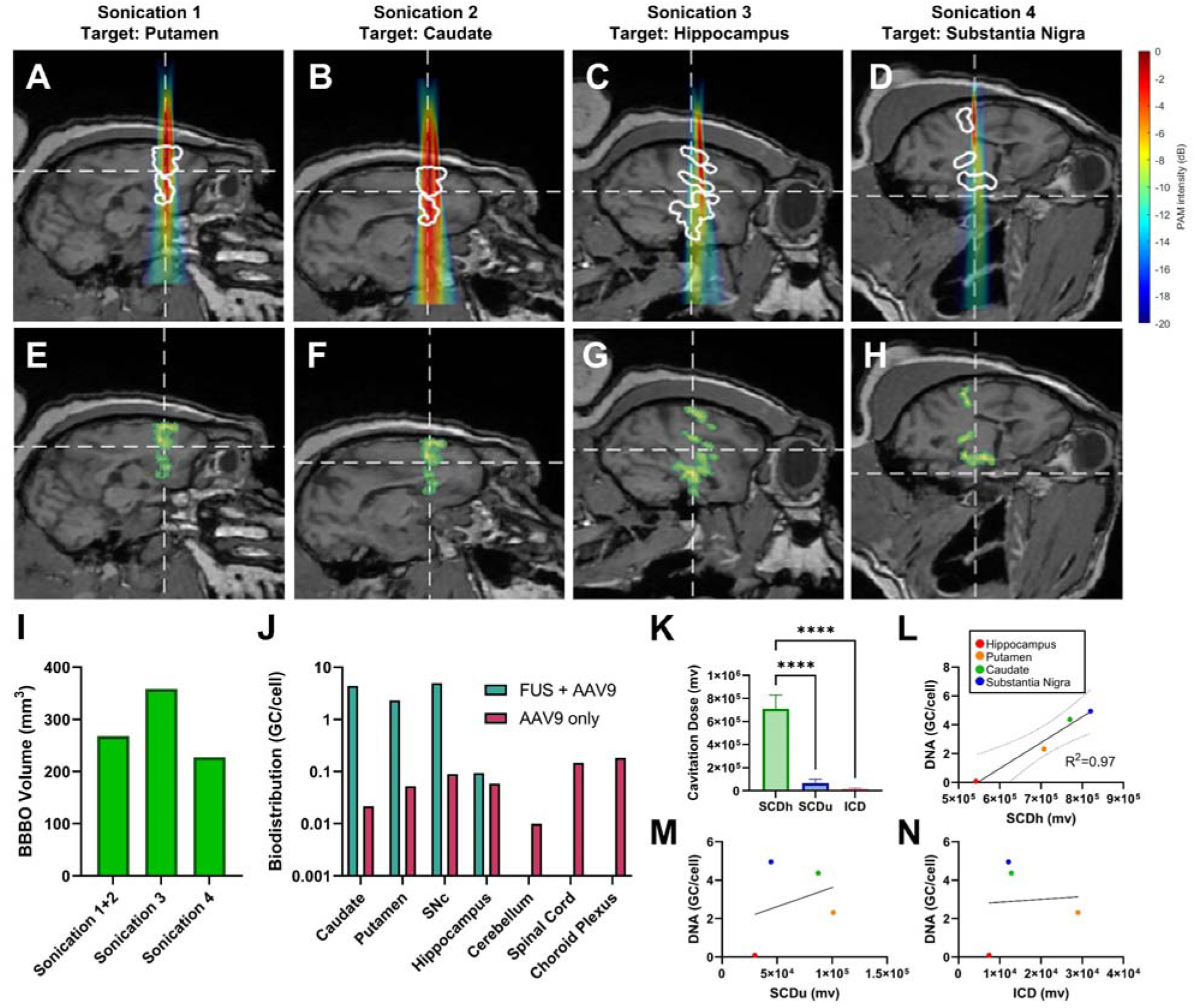
FUS-facilitated gene delivery correlated with PAM cavitation dose in NHP. **A.** PAM images overlaid onto the sagittal slice of MRI corresponding to the center of the PAM imaging plane for the **A)** putamen trajectory, **B)** caudate trajectory, **C)** hippocampus trajectory, and **D)** substantia nigra trajectory. White outlines correspond to green contours outlining the BBB opening volume in E-H. Maximum projection of contrast-enhanced BBB opening volume on MRI for **E)** putamen trajectory, **F)** caudate trajectory, **G)** hippocampus trajectory, and **H)** substantia nigra trajectories. Intersection of dashed lines indicate the center of the FUS focus. **I)** Quantification of BBB opening volume within targeted brain regions in NHP A. **J)** Biodistribution of AAV DNA within the CNS. **K)** Quantification of cavitation dose derived from real-time PAM. Statistical significance determined by one-way ANOVA with post-hoc Tukey’s multiple comparisons test. \*\*\*\**p*<0.0001. Linear relationship between vector genome copies per cell and **L)** stable harmonic cavitation dose (SCDh), **M)** stable ultraharmonic cavitation dose (SCDu) and **N)** inertial broadband cavitation dose (ICD) determined by standard linear regression.

**Figure 4:**
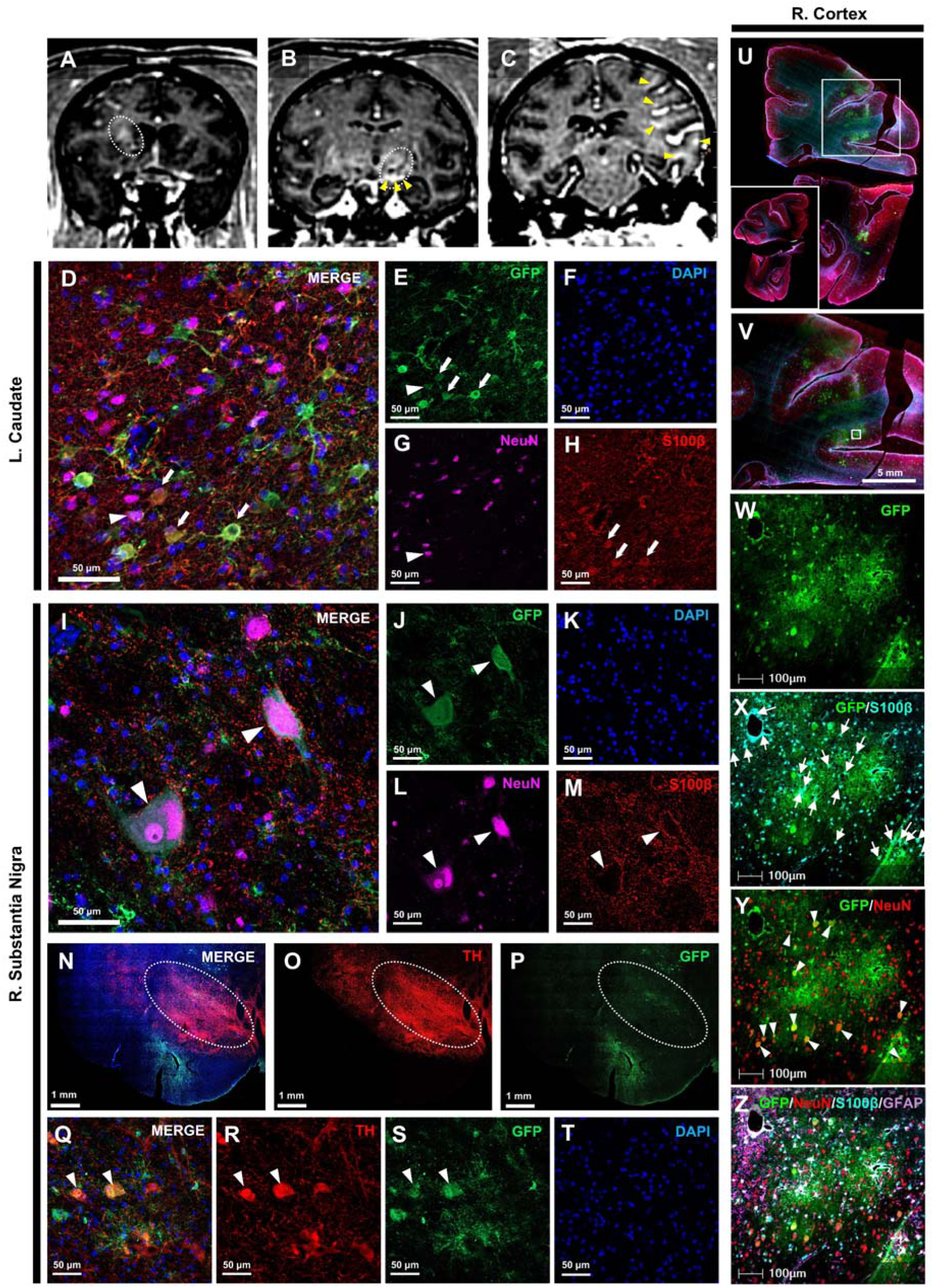
FUS-mediated AAV9-CAG-GFP delivery elicits astrocytic and neuronal transgene expression in several NHP brain regions. Raw coronal contrast-enhanced T_1_-weighted MRI for the sonications targeted to the **A)** left caudate (NHP A, sonications 1-2), **B)** right substantia nigra (NHP A, sonication 4), and **C)** right cortex (NHP B, single FUS sonication). White ellipses in A-B denote approximate ROIs of BBB opening, and yellow arrowheads in B-C denote less obvious ROIs within BBB openings. **D)** Merged image of GFP expression in the sonicated left caudate. Arrowheads denote transduced neurons, and arrows denote transduced astrocytes. Individual **E)** GFP, **F)** DAPI, **G)** NeuN, and **H)** S100β channels from merged image in (D). **I)** Merged image of GFP expression in the sonicated right substantia nigra. Arrowheads denote transduced neurons. Individual **J)** GFP, **K)** DAPI, **L)** NeuN and **M)** S100β channels from merged image in (I). **N)** Merged image of the right substantia nigra (horizontally flipped) with individual **O)** TH and **P)** GFP channels depicting GFP and TH signal overlap. White dashed ellipses denote ROI of GFP and TH overlap. **Q)** High magnification merged image of colocalized **R)** TH signal and **S)** GFP signal, with **T)** DAPI nuclei staining in the right substantia nigra from ROIs denoted in N-P. Arrowheads denote transduced dopaminergic neurons. **U)** GFP expression within the sonicated cortex in NHP B. Inset depicts contralateral cortex. **V)** Enlarged image of ROI shown in U. Enlarged images of **W)** GFP, **X)** GFP/S100β **Y)** GFP/NeuN, and **Z)** GFP/NeuN/S100β/GFAP channels from ROI in V.

Three weeks after FUS-mediated BBB opening, NHP A was euthanized for downstream tissue analysis of AAV9 delivery with ddPCR and IHC. Biodistribution analysis of tissue punches taken from the targeted and respective contralateral brain regions with ddPCR revealed a 200-fold improvement in AAV9 transduction in the caudate, a 44-fold improvement in the sonicated putamen, and a 50-fold improvement in transduction in the substantia nigra in FUS-treated regions relative to contralateral regions (Figure 3J). An average AAV9 DNA yield of 0.07 ± 0.06 gc/cell was observed in unsonicated regions of the CNS, representing the average transduction efficiency of AAV9 without transient FUS-mediated opening of the BBB, whereas an average of 2.94 ± 2.28 gc/cell was observed in FUS treated regions (Figure 3J), comprising an average 42-fold increase in AAV9 DNA achieved with FUS in NHP A. In NHP B, which was euthanized 4 weeks after FUS-mediated AAV9 delivery along the planned trajectory shown in Supplementary Figure 2A, qPCR of brain punches along the FUS-sonicated cortex revealed 20-fold, 13-fold, and 28-fold increases in vector DNA within the parietal, superior temporal, and ventral inferotemporal cortex, relative to the unsonicated, left cortical regions (Supplementary Figure 3).

Given that FUS sonications were monitored with real-time PAM using a separate confocally-aligned ultrasound imaging array, we sought to evaluate the association of cavitation dose with the amount of vector DNA quantified by ddPCR to determine whether PAM-based treatment monitoring could predict resulting gene expression. Quantification of cavitation dose across the 4 separate sonications in NHP A revealed significantly higher contributions of stable harmonic cavitation (SCDh) than stable ultraharmonic cavitation (SCDu) or inertial cavitation doses (ICD), indicating that the induced BBB opening volumes were primarily driven by stable cavitation during AAV delivery with FUS (Figure 3K). Remarkably, SCDh was highly correlated with the amount of vector DNA quantified within the FUS-targeted brain regions (R^2^ = 0.97, *p* = 0.0165, Figure 3L), while neither SCDu or ICD exhibited a correlation with resulting vector genome copies (Figure 3M-N). Given that SCDh was the primary contribution to the overall cavitation dose influencing BBB opening volume, and indeed exhibited the highest correlation with resulting vector DNA, cavitation dose could be a potential monitoring strategy for predicting gene delivery after FUS-mediated BBB opening.

Consistent with ddPCR-derived AAV biodistribution results, IHC revealed a marked increase in observable gene expression along all FUS-sonicated trajectories in NHP A (Figure 4A-B) and NHP B (Figure 4C) relative to unsonicated brain regions which altogether displayed extremely minimal GFP expression on fluorescence microscopy images. No GFP expression was detected in the hippocampus of NHP A, corresponding to sonication 3 (Figure 3G). Fluorescence microscopy images of the sonicated caudate target revealed substantial GFP expression predominantly in astrocytes as indicated by GFP colocalization with astrocyte marker S100β (Figure 4D-F,H), along with several neurons, depicted by GFP colocalization with neuronal marker NeuN (Figure 4D-G). IHC conducted in the substantia nigra depicted both neuronal and astrocytic transduction (Figure 4I-L), where TH staining revealed GFP expression within the substantia nigra (Figure 4N-P), specifically within dopaminergic neurons as demonstrated by colocalization between GFP and TH signal in Figure 4Q-T. GFP expression was observed along the FUS trajectory within various cortical structures in NHP B, whereas the contralateral hemisphere was devoid of detectable GFP expression (Figure 4U). The regions with the most GFP expression (Figure 4V) also overlapped with contrast-enhanced regions on the T_1_-weighted MRI shown in Figure 4C. GFP (Figure 4W) colocalization with astrocytic and neuronal (Figure 4X-Z) markers exhibited similar relative levels of GFP expression across FUS treated brain regions.

While astrocytes and neurons comprised the majority of cells expressing GFP after FUS-facilitated AAV delivery, other cell types including oligodendrocytes (Supplementary Figure 4A-H) were also transduced but comprised a significant minority of cells transduced within all regions evaluated. No microglia transduction was observed as depicted by lack of colocalization of GFP with microglia marker IBA1 (Supplementary Figure 4I-L).

### Systemic AAV9 delivery and BBB opening with a novel 500 kHz theranostic ultrasound linear array yields targeted AAV9 transduction with a low-cost portable therapeutic ultrasound system

ThUS is a novel low-frequency ultrasound imaging array configuration designed to enhance the potential for point-of-care therapeutic ultrasound of the brain through use of existing clinical diagnostic ultrasound scanner hardware. In addition to evaluating the feasibility of AAV9 delivery with a conventional clinic-ready USgFUS prototype system (*16*), we also sought to determine whether ThUS could elicit BBB opening and AAV9 delivery in NHP.

Prior to AAV delivery experiments, we evaluated the feasibility of using ThUS to target brain regions implicated in neurodegeneration in PD in two NHPs: NHP B (targeting the caudate, putamen, and periaqueductal gray) and NHP C (targeting the caudate, putamen, and hippocampus) (Supplementary Figure 8). A single ThUS session targeting the substantia nigra in NHP B elicited successful BBB opening as confirmed by contrast-enhanced T_1_-weighted MRI acquired ∼1 hour post-ThUS (Figure 5A-C). PCI acquired and displayed during the sonication (Figure 5D) agreed well with the volume of BBB opening as shown in the superimposed maximum intensity projection contours denoting the BBB opening volume in Figure 5E-F. This level of agreement underscores the utility of high-resolution real-time PCI to monitor BBB opening with ThUS, particularly when compared qualitatively to the less resolved PAM images in Figure 3. In addition to targeting the substantia nigra, we also targeted the caudate and putamen simultaneously using the ThUS rapid alternating steering angles (RASTA) pulse sequence (*31*) with two steering angles, one targeted 2.5 mm anterior and the second targeted 2.5 mm posterior to the line normal to the array face (Supplementary Figure 6A-B). This yielded two BBB opening trajectories during a single sonication as depicted on contrast-enhanced T_1_-weighted MRI (Figure 5G-J), using only a single bolus microbubble injection. Overall, PCI signal intensity was lower in the striatum when compared to the substantia nigra, potentially due to a number of factors including the beam incidence angle (Supplementary Figure 6C-L) and reduced effective PRF when using the RASTA sequence, as discussed in the Supplementary Material. However, BBB openings in Figure 5M denoted by the white contours in Figure 5L still agreed spatially with PCI acquired during ThUS RASTA (Figure 5K). Most notably, over all sonications targeted to the substantia nigra and striatum, a high correlation between the cumulative PCI signal intensity (representing the total cavitation dose over the sonication) and induced BBB opening volume was observed (Figure 5N, R^2^ = 0.88, *p* = 0.0017), further emphasizing the value of quantitative PCI employed during sonication.

**Figure 5:**
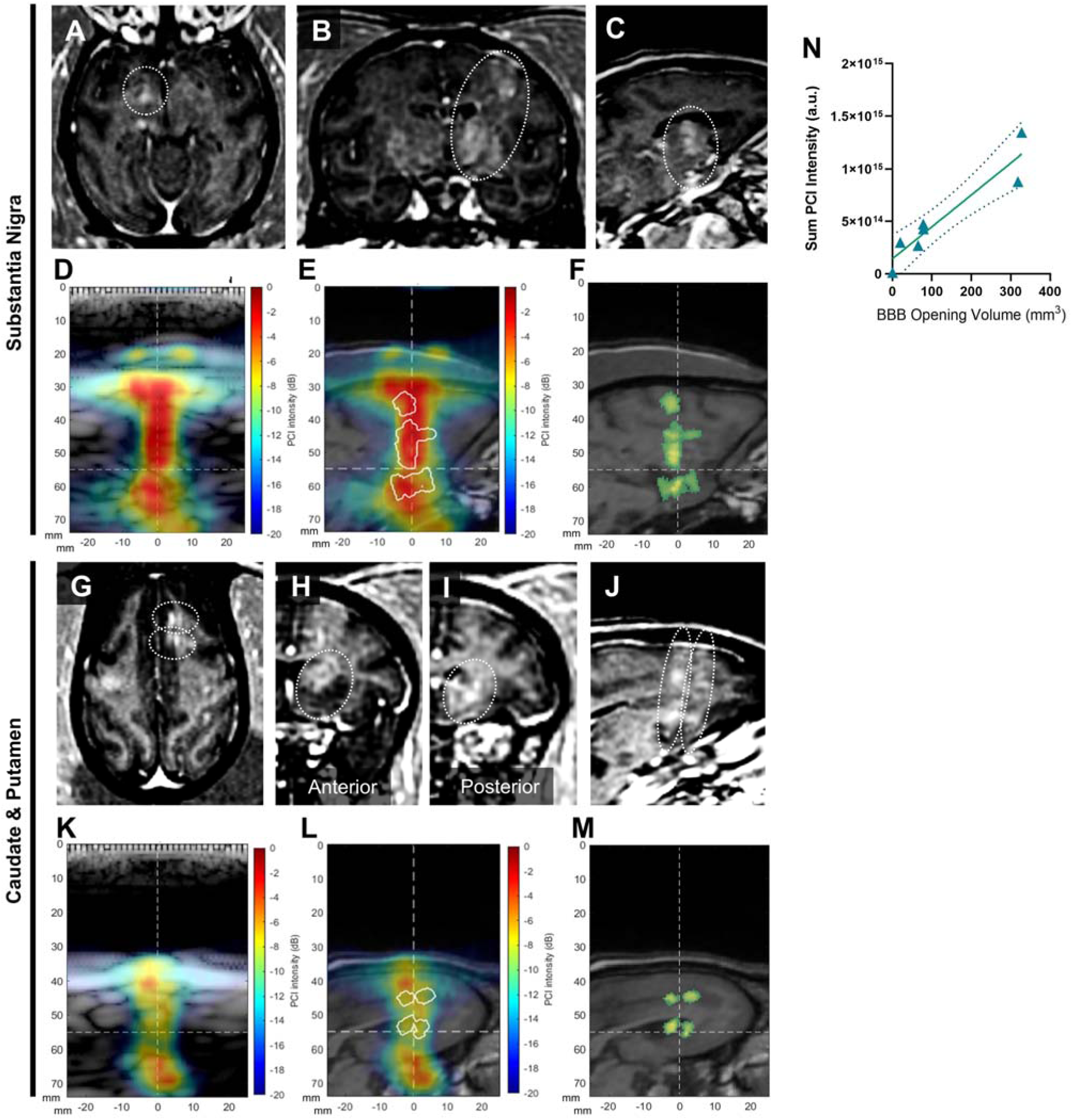
Feasibility of BBB opening in PD-afflicted brain regions with ThUS in NHP. **A)** Axial **B)** coronal and **C)** sagittal contrast-enhanced T_1_-weighted MRI of a ThUS-mediated BBB opening with a trajectory targeted through the substantia nigra. Corresponding summed PCI acquired during the sonication overlaid onto **D)** pre-sonication B-mode and **E)** slice of T_1_-weighted MRI corresponding to the center of the BBB opening volume denoted by the white contours. **F)** Same slice of T_1_-weighted MRI as (E) with green overlay depicting region of contrast enhancement after BBB opening. **G)** Axial, **H)** anterior coronal, **I)** posterior coronal, and **J)** sagittal contrast-enhanced T_1_-weighted MRI of a BBB opening induced with ThUS RASTA with trajectories targeted to the caudate and putamen. Corresponding summed PCI acquired during the sonication overlaid onto **K)** pre-sonication B-mode and **L)** slice of T_1_-weighted MRI corresponding to the center of the BBB opening volume denoted by the white contours. **M)** Same slice of T1-weighted MRI as (L) with green overlay depicting region of contrast enhancement after BBB opening. **N)** Correlation between cumulative PCI pixel intensity and induced BBB opening volume. n=6 sonications, R^2^ = 0.88 determined by standard linear regression.

Given that we consistently induced successful BBB opening with ThUS, we aimed to determine whether this low-frequency multi-element linear array configuration could deliver AAV to the NHP brain through the study presented in Figure 1F. Immediately after beginning the sonication with ThUS RASTA targeted to both the anterior and posterior portions of the striatum in NHP B, a bolus of microbubbles followed by 2.0e13 gc/kg AAV9-CAG-GFP was intravenously injected. After sonication for 4 minutes, the ThUS array was repositioned to target the substantia nigra where a second bolus injection of microbubbles was administered before sonication for two minutes with a single ThUS focus. Finally, the ThUS array was repositioned to target a trajectory through the midbrain before another bolus injection of microbubbles and sonication for two minutes. BBB opening was confirmed along each of the three ThUS-targeted trajectories as shown in Figure 6A-C, with good spatial agreement between BBB opening volumes (Supplementary Video 2) and PCI (Supplementary Figure 7).

**Figure 6:**
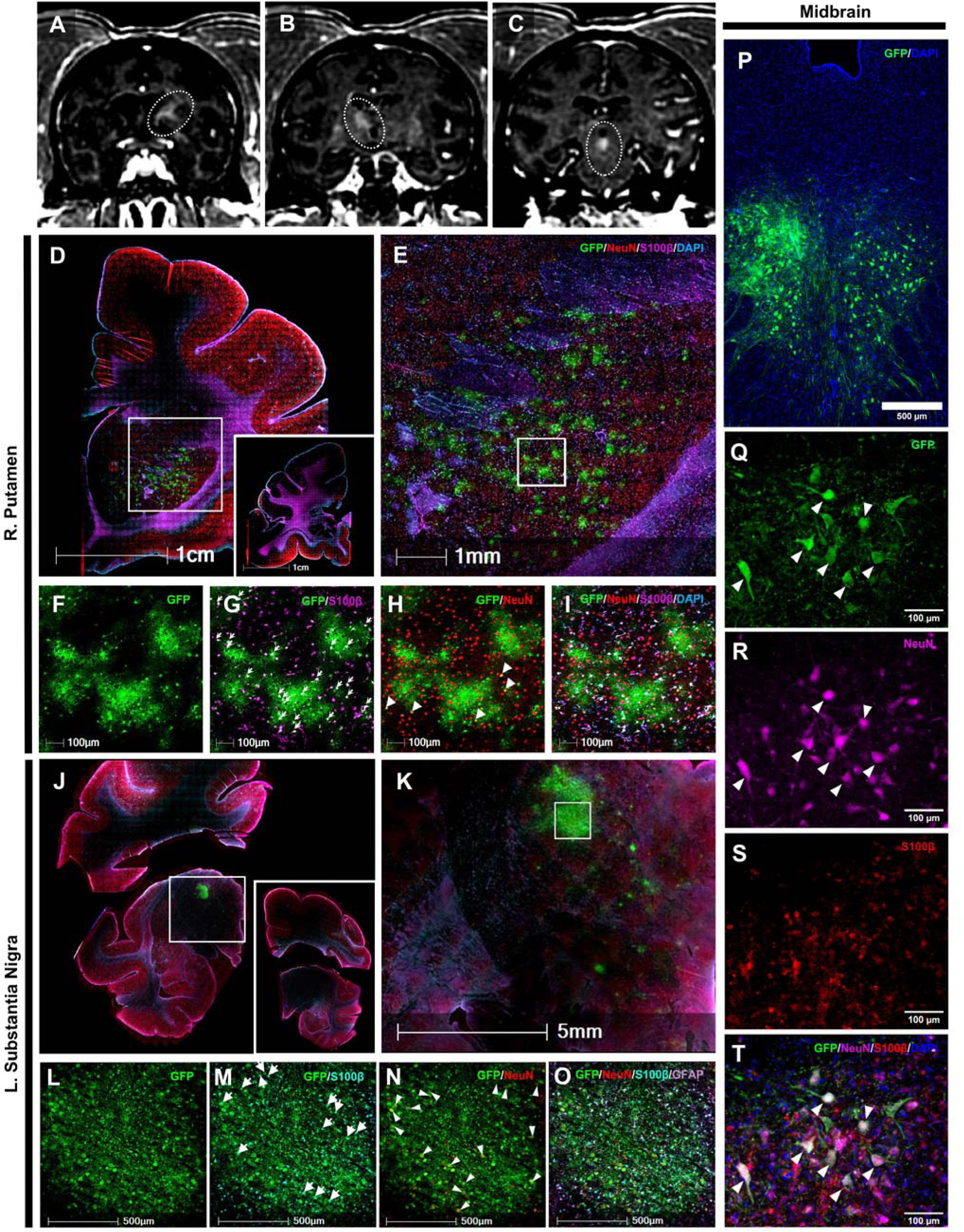
AAV9-CAG-GFP delivery with ThUS elicits targeted astrocytic and neuronal gene expression within several NHP brain regions. Raw coronal contrast-enhanced T1-weighted MRI for the sonications targeted to the **A)** right putamen, **B)** left substantia nigra, and **C)** midbrain in NHP B. **D)** Scanned fluorescence microscopy image depicting GFP expression within the sonicated putamen. Inset on right corresponds to unsonicated contralateral hemisphere. **E)** Enlarged image of rectangular ROI in D depicting GFP expression within the putamen. **F)** GFP expression within the rectangular ROI in E. **G)** GFP and S100β colocalization denoted by white arrows. **H)** GFP and NeuN colocalization denoted by white arrowheads. **I)** Merged image of GFP, S100β, NeuN, and DAPI staining from F-H. **J)** Image of GFP expression within the thalamus along the SN-targeted trajectory. Inset denotes contralateral hemisphere. **K)** Enlarged image of ROI in J. Enlarged images of ROI in K) depicting **L)** GFP, **M)** GFP/ S100β, **N)** GFP/NeuN, and **O)** GFP/NeuN/S100ß/GFAP channels. **P)** GFP expression within the periaqueductal gray region of the midbrain. **Q)** GFP expression and **R)** NeuN staining depicting neuronal GFP expression. **S)** S100β staining revealed no astrocytic gene expression in oculomotor nucleus. **T)** Merged image of GFP, NeuN, S100β and DAPI staining depicting neuronal GFP expression denoted by white arrows.

NHP B was closely monitored for one month after ThUS to allow for gene expression to occur before euthanasia by transcardial perfusion. The brain and peripheral organs were then prepared for biodistribution and histology assays as detailed in the methods section. After immunostaining and microscopy, GFP expression was observed along all three trajectories. Within the putamen (Figure 6D), GFP expression was predominantly observed within astrocytes (Figure 6E), depicted by colocalization of GFP with astrocytic marker S100β (Figure 6F-G,I). GFP+ neurons were also detected within the sonicated putamen depicted by colocalization of GFP with neuronal marker NeuN in Figure 6F,H,I. Along the substantia nigra trajectory, GFP expression was localized within a small region in the thalamus, slightly dorsal to the substantia nigra pars compacta (Figure 6J-K), within astrocytes (Figure 6L-M,O) and neurons (Figure 6L,N-O) at a distribution which agreed qualitatively with other transduced brain regions in the study. However, within the sonicated midbrain, specifically the periaqueductal gray region (Figure 6P), neurons were primarily transduced as demonstrated by colocalization of GFP with NeuN in Figure 6Q,R,T. No astrocytic GFP expression was observed in this region (Figure 6S), revealing a relationship between brain structure and cellular tropism exhibited by AAV9-CAG-GFP within the NHP brain. Taken together, these results demonstrate that AAV9-mediated gene expression is expected to occur within multiple cerebral cell types, but the relative expression level between cell types depends on the brain region targeted with therapeutic ultrasound.

Overall, the resulting level of gene expression appeared to exhibit a higher level of dependence on brain region rather than the volume of BBB opening induced. While the 78.65 mm^3^ BBB opening volume along the substantia nigra trajectory (Supplementary Figure 7E) was 4-fold greater than the 19.61 mm^3^ BBB opening induced in the striatum (Supplementary Figure 7D), gene expression within the putamen (Figure 6D) appeared to exceed that of the level of expression observed along the substantia nigra trajectory within the thalamus and substantia nigra pars compacta (Figure 6J-K). Additionally, while the BBB opening volume along the midbrain trajectory encompassed discontinuous volumes surrounding the periaqueductal gray region where gene expression is visible in Figure 6P, the more dorsal BBB opening regions shown in Supplementary Figure 7F were devoid of observable GFP expression. This apparent discrepancy may indeed be caused by a number of factors including the tissue allocation method used for downstream assays, but also may reflect the specific regional transduction patterns of AAV9 within the rhesus macaque brain, as was also observed in mice (*48*).

### Safety of FUS- and ThUS-facilitated BBB opening in NHP after intravenous administration of AAV9

To evaluate the *in vivo* safety of both FUS and ThUS sonication procedures in the NHPs, T_2_-weighted fluid-attenuated inversion recovery (T_2_-FLAIR) imaging was conducted immediately prior to gadodiamide injection for contrast-enhanced T_1_-weighted imaging (Figure 7A-D, P-R) in both NHPs on the day of AAV delivery. For FUS sonications, T_2_-FLAIR images were devoid of hyperintensities along all trajectories (Figure 7E-G) apart from the BBB opening region in the right cortex of NHP B, where small hyperintensities depicting minor acute edema were detected (Figure 7H). In conducting histological analysis, we also discovered evidence of a potential lasting astrocytic immune response within part of the aforementioned cortical BBB opening in NHP B (Figure 7I-J). Regions with persistent astrocytic activation exhibited a high proportion of GFAP expression (Figure 7M) within S100β+ astrocytes (Figure 7O), represented visually by colocalization between GFP, S100β, and GFAP (Figure 7J). Given that no other FUS or ThUS-sonicated regions expressing GFP exhibited both T_2_-FLAIR hyperintensities and evidence of persistent astrocyte activation, this instance was likely due to a deviation in the treated trajectory from the planned and simulated trajectory. For ThUS sonications, no obvious hyperintensities were detected within the BBB opening regions (Figure 7P-R) in any of the targets sonicated with ThUS on T_2_-FLAIR images (Figure 7S-U).

**Figure 7:**
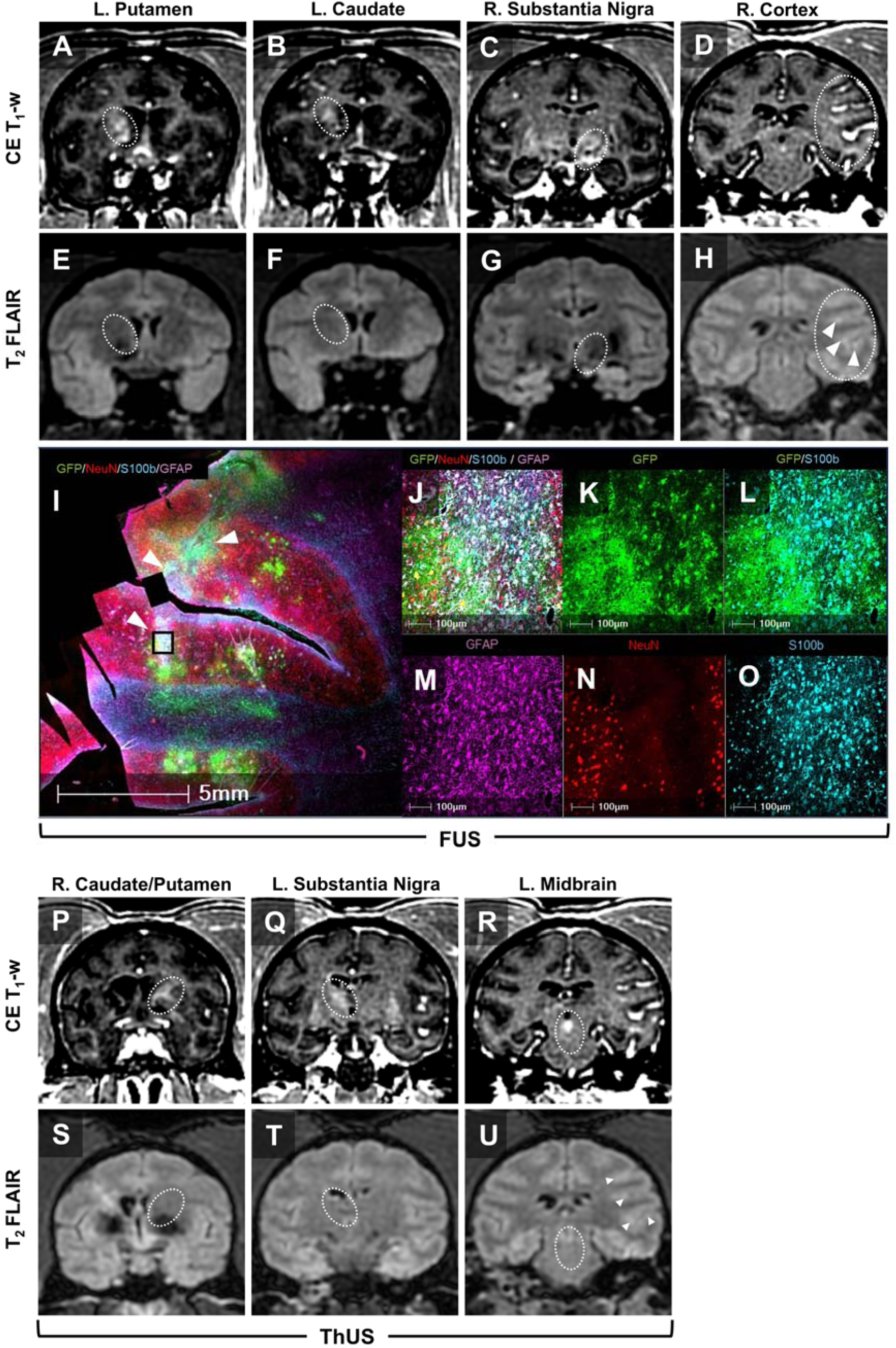
Safety of ThUS- and FUS-mediated AAV9-CAG-GFP delivery in NHP. Contrast-enhanced T_1_-weighted MRI depicting ThUS-mediated BBB opening in the **A)** left putamen, **B)** left caudate, **C)** right substantia nigra, and **D)** right cortex (NHP B) with corresponding T_2_-FLAIR MRI shown in **E-H).** Dashed ellipse ROI denotes region of BBB opening. White arrowheads in (H) correspond to hyperintense regions on T_2_-FLAIR. **I)** Fluorescence microscopy image depicting astrocytic immune response in FUS-sonicated cortex. White arrowheads denote regions of high GFAP and S100β colocalization. **J)** Multichannel fluorescence microscopy image from region in (I) denoted by black rectangular ROI. **K)** GFP, **L)** GFP/ S100β, **M)** GFAP, **N)** NeuN, and **O)** S100β channels. Contrast-enhanced T_1_-weighted MRI depicting ThUS-mediated BBB opening in **P)** right caudate/putamen, **Q)** left substantia nigra and **R)** left midbrain with corresponding T_2_-FLAIR MRI shown in **S-U).** Dashed ellipse ROI denotes region of BBB opening. The pre-existing hyperintensity on the contralateral hemisphere in (S) was not study related. Arrowheads in (U) denote minor hyperintensities from a separate FUS sonication not induced by ThUS.

Preliminary ddPCR results of peripheral tissue samples from NHP A provided insight into the relationship between targeted transduction within the brain as a result of BBB opening with FUS and peripheral tissue transduction after a systemic injection of AAV9. Tissue samples from NHP A revealed systemic biodistribution of 181 gc/cell in the liver, and 3 gc/cell in the heart—a magnitude which is consistent with the level expected at the injected dose of 2.0e13 gc/kg. Given the high level of off-target gene expression in peripheral tissue, and majority of gene transduction occurring in astrocytes, novel capsid engineering strategies will be leveraged for increased neuronal targeting and liver de-targeting in future experiments.

### Systemic AAV9-hNTRN delivery and bilateral BBB opening with ThUS facilitates neurorestoration in the 1-methyl-4-phenyl-1,2,3,6-tetrahydropyridine (MPTP) mouse model

Given that ThUS enabled non-invasive gene delivery to the NHP brain in structures implicated in PD, we next evaluated the therapeutic potential for ThUS-mediated AAV delivery by assessing the restoration of a partially-degenerated nigrostriatal pathway in the subacute MPTP mouse model. This mouse model enables selective degeneration of nigrostriatal neurons at a severity determined by the dosing regimen (*47*) and has been routinely used in pre-clinical PD research in both mice and NHP. After a 21-day period of neurodegeneration post-MPTP dosing corresponding to the timeline shown in Figure 8A, mice underwent bilateral sonication using two anatomically symmetric focal volumes encompassing the striatum and substantia nigra, achieved by ThUS RASTA, and/or received AAV as described in the methods section. Contrast-enhanced T_1_-weighted MRI acquired on sonicated animals displayed bilateral BBB opening extending throughout both the substantia nigra and posterior CPu (Figure 8B). Representative summed PCI over the 2-minute sonication duration are shown in Supplementary Figure 9D, where no significant differences in PCI cumulative pixel intensity were observed between the MPTP + ThUS and MPTP + ThUS + AAV groups (Supplementary Figure 9E), indicating that any post-treatment effects are primarily due to AAV administration, not differences in ThUS-mediated BBB opening.

**Figure 8:**
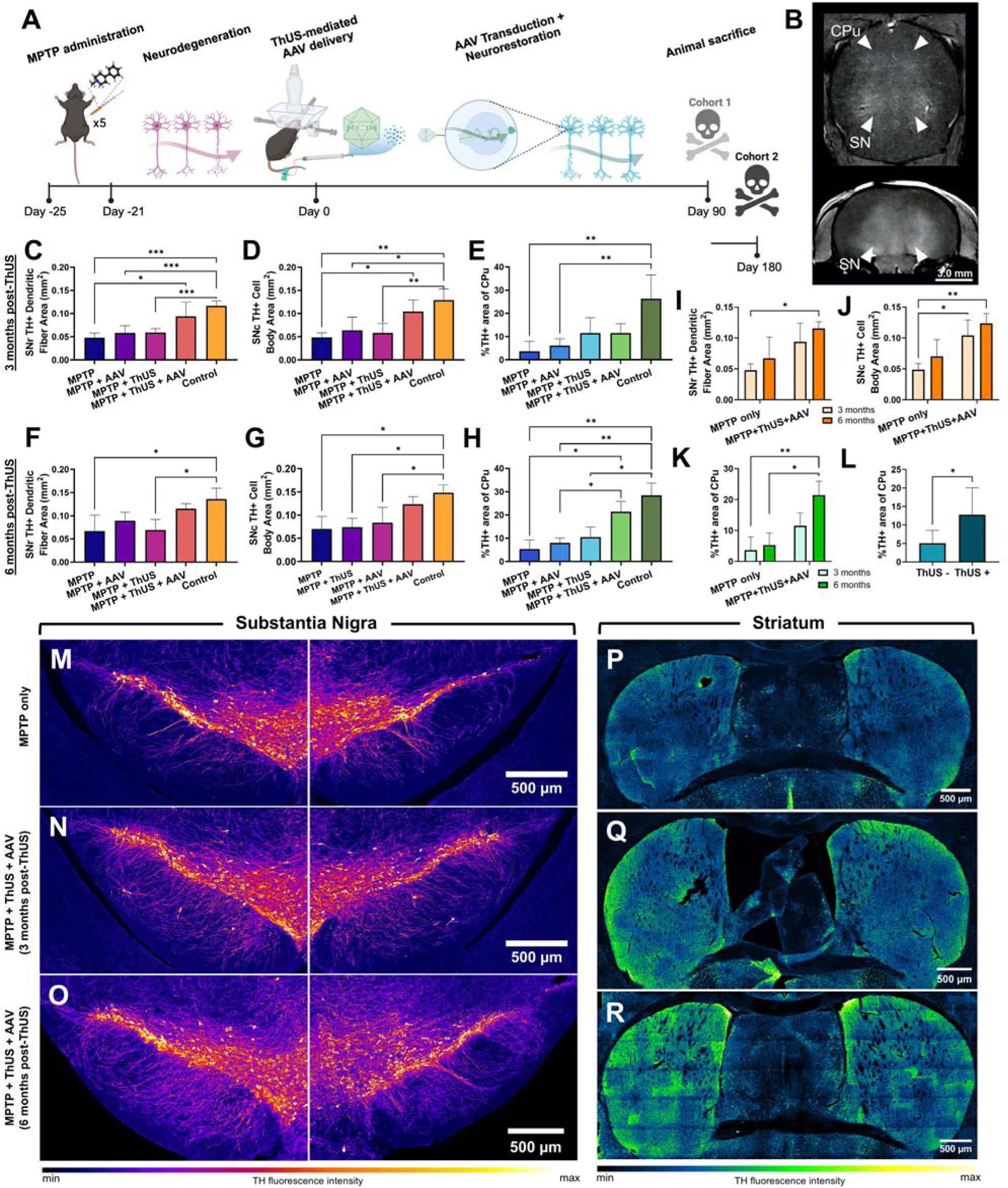
ThUS-mediated AAV9-hSyn-hNTRN delivery facilitates neurorestoration in MPTP mice. **A)** Timeline of experiment from MPTP dosing to euthanasia. **B)** Representative contrast-enhanced T1-weighted MRI acquired ∼ 30 min post-ThUS and AAV dosing. Bilateral ThUS targeting elicited BBB opening in both the striatum (CPu) and substantia nigra (SN) with a single sonication. **C)** Groupwise quantification and comparison of fluorescence area of dendritic network within SNr, and **D)** cell bodies within SNc. **E)** Groupwise comparison of %TH positivity over striatal area. **F)** Groupwise comparison of fluorescence area of dendritic network within SNr, and **G)** cell bodies within SNc. **H)** Groupwise comparison of %TH positivity over striatal area. Comparisons between TH immunoreactivity in **I)** SNr and **J)** SNc in MPTP mice and MPTP+ThUS+AAV mice after 3 months and 6 months post-treatment. **K)** Groupwise comparisons between CPu TH immunoreactivity in MPTP mice and MPTP+ThUS+AAV mice after 3 months and 6 months post treatment. **L)** Groupwise comparisons between all mice not receiving ThUS (MPTP only and MPTP+AAV groups) and mice receiving ThUS BBB opening (MPTP+ThUS and MPTP+ThUS+AAV groups). Representative pseudo-colored fluorescence microscopy images of left and right SN in **M)** a mouse receiving only MPTP injections and no therapeutic intervention, and mice receiving MPTP injections and ThUS+AAV treatment **N)** 3 months post-dosing, and **O)** 6 months post dosing. Representative pseudo-colored fluorescence microscopy images of striata of mice receiving **P)** only MPTP injections and no therapeutic intervention, and mice receiving MPTP injections and ThUS+AAV treatment **Q)** 3 months post-dosing, and **R)** 6 months post-dosing. Statistical significance in C-H determined by one-way ANOVA with post-hoc Tukey’s multiple comparisons correction. *\*p<0.05*, *\*\*p<0.01*, *\*\*\*p<0.001*, N = 4 per group. Statistical significance in I-K determined by two-way ANOVA with post-hoc Tukey’s multiple comparisons correction, *\*p<*0.05, *\*\*p<*0.01, n=3-4 per group.

After a survival period of either 3 months or 6 months post-AAV dosing, mice were euthanized and neurorestoration was evaluated with TH immunohistochemistry on coronal cryosections from the substantia nigra and caudate putamen. Image analysis of TH-stained sections containing the substantia nigra (Figure 8M-O) revealed a ∼96 % significant increase in SNr dendritic fiber network density 3 months post-dosing in MPTP mice treated with ThUS+AAV relative to untreated MPTP mice (Figure 8C), with no additional significant improvement observed after 6 months (Figure 8F). A ∼107% significant increase in cell body density in the SNc was observed 3 months post-dosing in the MPTP + ThUS + AAV group relative to untreated MPTP mice (Figure 8D), with a small but statistically insignificant improvement in the average area of SNc TH immunoreactivity after 6 months (Figure 8G). Additionally, nearly 3-fold reductions in dendritic fiber density and cell body area were calculated in the MPTP group relative to healthy wild-type controls at the 3-month timepoint, attesting to the effectiveness of the MPTP dosing protocol (Figure 8C-D). No significant differences in dendrite fiber network density or cell body area were observed in MPTP mice receiving either AAV or ThUS compared to untreated MPTP mice at either timepoint. However, such quantities in the MPTP + ThUS + AAV group were statistically indistinguishable from wild-type control mice at both timepoints (Figure 8C-D, F-G, Supplementary Figure 9A-B), although trends indicated incomplete neuronal rescue. These results demonstrate that ThUS-facilitated hNTRN gene delivery may induce lasting neurorestorative effects in the substantia nigra.

In addition to analyzing TH density in sections containing the substantia nigra, TH immunoreactivity in the caudate putamen, corresponding to axon terminals of neurons whose cell bodies reside in the substantia nigra, as shown in the light sheet microscopy image in Supplementary Figure 9C, was also quantified and compared across groups at both timepoints (Figure 8P-R). Quantification of TH immunoreactivity in the caudate putamen demonstrated an ∼3-fold trending increase in the MPTP + ThUS + AAV group relative to untreated MPTP mice after 3 months post-dosing (Figure 8E), albeit statistically insignificant. Interestingly, the ThUS treatment alone conferred a greater average increase in terminal area than all other treatment interventions (Figure 8E), which was not significantly different than healthy control mice after 3 months post-dosing. Additionally, comparisons between the mean TH+ density of groups receiving ThUS (MPTP + ThUS and MPTP + ThUS + AAV) and groups not receiving ThUS (MPTP only and MPTP + AAV) revealed a significant increase in TH immunolabeling in ThUS-treated groups, whether receiving AAV or not (Figure 8L). Given the statistically significant differences in dendrite growth and density of cell bodies within the substantia nigra yet trending but not statistically significant differences between axon terminal staining between MPTP mice and MPTP + ThUS + AAV mice, we sought to determine whether increased neurorestoration would occur as a result of longer cumulative treatment effects. In addition to observing generally increased TH signal within the striata of MPTP mice treated with ThUS+AAV (Figure 8R), we observed a significant ∼4-fold average increase in TH+ terminal density relative to the MPTP only group (Figure 8H), corresponding to nearly a ∼2-fold increase in TH+ terminal density in the striatum from 3 months to 6 months post-ThUS+AAV (Figure 8K). This indicates that while neurorestoration is statistically increased 3 months post ThUS+AAV relative to untreated mice in the SN (Figure 8I-J), a potentially longer duration of hNTRN overexpression is required to induce significant changes in neurorestoration at the axon terminal level. Together, these results indicate that ThUS-mediated hNTRN delivery induces both short-term and long-term neurorestoration, as confirmed by histological analysis in two main regions of the nigrostriatal pathway implicated in dopaminergic neuron loss in early PD.

## DISCUSSION

In this study, the most novel results demonstrate a marked increase in AAV9-CAG-GFP delivery across the BBB in multiple brain regions, achieved with two different USgFUS configurations in geriatric rhesus macaques. This NHP study was informed by an AAV9 dose escalation study in wild-type mice, where we observed significant increases in hippocampal FUS-delivered AAV9 DNA after increasing the systemic AAV9 dose, supported by both biochemical and immunohistochemical outcomes. We elected to target brain regions implicated in PD including the substantia nigra, caudate, and putamen for our NHP study to elucidate the effectiveness of USgFUS for facilitating non-invasive gene therapy in neurodegenerative disorders. With FUS+PAM, we achieved BBB opening volumes ranging from 267 mm^3^ in the striatum to over 800 mm^3^ in the cortex. The increase in targeted BBB permeability yielded an average increase in vector DNA over contralateral structures of nearly 50-fold, comprising the first quantitative biochemical readout of AAV transduction within FUS-targeted regions in NHP. No substantial increase in vector DNA was observed in targeted regions where no BBB opening was detected, such as the hippocampus which proved to be challenging to target due to poor FUS beam incidence angle with respect to the skull curvature. Note that for complex trajectories, acoustic holograms coupled to a single-element transducer have been shown to achieve deeper and more selective targeting (*49, 50*). However, the hippocampal targeting difficulties in the present study were also resolved when using the ThUS linear array in NHP C, marking an advantage of the linear array ThUS configuration over the spherically focused FUS configuration. Interestingly, SCDh which has been shown to correlate with BBB opening volume (*51*), was directly correlated with the quantity of vector DNA within the targeted regions, where BBB opening volume quantified by contrast-enhancement on post-FUS T_1_-weighted MRI failed to exhibit such a correlation. This attests to the significant advantage that USgFUS systems may pose in cavitation-based viral delivery across the BBB. IHC also confirmed targeted GFP transgene expression within the targeted regions in astrocytes, neurons including dopaminergic neurons within the substantia nigra, and a small number of oligodendrocytes.

Given the previously demonstrated portability, cost, and flexibility advantages of ThUS for AAV delivery in mice (*31*), we also evaluated the feasibility for ThUS-mediated targeted AAV delivery in NHP to elucidate the advantages and shortcomings of this novel linear array theranostic system as USgFUS development progresses towards clinical translation of non-invasive gene therapy. Opening volumes induced by ThUS were smaller overall when compared to FUS (∼20-80 mm^3^) primarily due to the 2-fold higher ThUS transmit frequency relative to the FUS configuration used in this study. We also revealed important relationships between integrated high-resolution PCI and resulting BBB opening volume, as PCI pixel intensity was directly related to BBB opening volume and spatially agreed with contrast-enhanced regions on T_1_-weighted MRI. In evaluating the ability of ThUS RASTA to induce BBB opening in multiple structures within the same sonication, specifically the caudate and putamen, we found that success of multi-focal BBB opening with RASTA highly depends on the incidence angle of the ThUS-steered beam to the skull, noting a decrease in BBB opening volume and PCI intensity of nearly 4-fold and 1.5-fold, respectively, with a sub-optimal incidence angle as confirmed by pre-sonication B-mode ultrasound imaging with the ThUS array. This also motivates the future use of B-mode for targeting refinement and PCI-based monitoring to correct for sub-optimal BBB opening during treatment. Unfortunately, the BBB opening most affected by suboptimal incidence angle within the striatum occurred on the day of AAV delivery, but despite this, targeted GFP expression was still confirmed on IHC in the sonicated putamen in addition to all other trajectories targeted with ThUS. No apparent differences in the proportion of astrocytic transduction versus neuronal transduction between FUS and ThUS were observed.

We also evaluated safety of both FUS and ThUS parameters used for AAV delivery through examination of T_2_-FLAIR MRI and neuroimmune activation signatures within fluorescence microscopy images. One occurrence of edema on T_2_-FLAIR MRI corresponding to high astrocytic GFAP expression in histology was observed along the cortical FUS trajectory in NHP B which exhibited a nearly 4-fold increase in BBB opening volume relative to the other FUS-induced BBB opening volumes. However, for all other BBB openings induced by either FUS or ThUS, no hyperintensities were observed in T_2_-FLAIR imaging, attesting to the safety of the USgFUS transmit parameters used in this study. Due to the intravenous route of AAV administration, substantial peripheral transduction occurred most prominently in the liver. This is a substantial limitation of targeted AAV delivery with transcranial FUS after a systemic AAV injection, which motivates the application of viral vector capsid engineering strategies to increase targeting efficacy (*52, 53*), or exploration of other, more direct routes of AAV administration such as intranasal delivery (*20, 54*).

Overall, the results presented herein offer a crucial demonstrated relationship between systemic dose and expected vector DNA in FUS-targeted regions, while confirming the feasibility of both long-pulse FUS and short-pulse ThUS modalities to elicit focal AAV delivery across the BBB in rhesus macaques. In addition, it is worth mentioning that the total treatment duration, from the beginning of the first sonication to end of the last sonication of 4 targets ranged from 30-45 min for each NHP, which is a major advantage of portable USgFUS for BBB opening and AAV delivery outside of an MRI scanner. We also waited ∼10 minutes between sonications to allow for microbubble clearance from the brain vasculature in an effort to associate BBB opening volume with a given concentration of injected microbubbles for experimental design purposes rather than allowing for microbubbles to accumulate in the vasculature from successive microbubble injections. With microbubble infusion methods, the treatment durations could be reduced further, potentially enabling realization of a fast, outpatient, non-invasive gene therapy procedure in patients in the future.

In the present study, we also evaluated bulk neurorestoration effects due to ThUS-mediated AAV9-hNTRN delivery in Parkinsonian MPTP mice via TH immunohistochemical quantification of neuron dendritic processes in the SNr, cell soma density in the SNc, and axon terminal density in the striatum up to 6 months post-treatment. While the prior NHP study demonstrated the feasibility for AAV9 delivery with portable USgFUS modalities, this mouse study was conducted to evaluate the therapeutic efficacy of non-invasive gene therapy with our most flexible therapeutic ultrasound platform, ThUS. Three months after AAV9-hNTRN delivery with ThUS BBB opening, we observed up to a 96% increase in dendritic network density in the SNr, and nearly double the density of cell soma within the SNc. Consistent with these observations in the substantia nigra, we also observed a 3-fold increase in axon terminal density in the striatum in MPTP mice receiving ThUS+AAV treatment. Interestingly, axon terminal density nearly doubled 6 months after AAV9-hNTRN delivery with ThUS, while no significant differences in SNr or SNc TH immunopositivity between 3 months and 6 months occurred. Given our use of the sub-acute MPTP dosing scheme to induce relatively minor nigrostriatal neuronal loss relative to other modes of MPTP insult (*47*), we also considered the possibility for natural reversal of neurodegeneration over time. This effect was made evident by comparison of TH immunopositivity in the substantia nigra between 3 and 6 months. We observed an average ∼50% increase in SNr and SNc density in the MPTP mice euthanized at the 6-month timepoint (corresponding to nearly 7 months after the first MPTP injection), relative to MPTP mice analyzed at the 3-month timepoint. To our knowledge, no prior studies have confirmed the length and stability of MPTP-induced neurodegeneration up through 6 months after sub-acute dosing (*47, 55, 56*).

While an extensive body of literature supports the observation of neuroprotection or neurorestoration with NTF overexpression either through AAV delivery or recombinant protein infusion (*57–63*), these results were not sufficiently recapitulated in human clinical trials. In this study, the non-invasive route of delivery facilitated by a next-generation therapeutic ultrasound device like ThUS could enable potentially increased efficacy with reduced complications with further optimization of AAV dosing and ThUS parameters. Karakatsani et al., demonstrated significant increases in substantia nigra and striatal TH immunoreactivity with multiple sessions of unilateral FUS-mediated GDNF delivery, or a single session of unilateral FUS-mediated AAV9-GDNF delivery in MPTP mice in both neuroprotective and neurorestorative experimental designs (*41*). Additionally, Mead et al., observed reduction of behavioral deficits and neurodegeneration in 6-OHDA mice up to 10 weeks after unilateral MR-guided FUS delivery of brain penetrating nanoparticles (BPN) containing GDNF, which enabled uniform striatal coverage (*64*). Our study is the first to leverage simultaneous image-guided bilateral delivery of AAV9-hNTRN to both the substantia nigra and striatum, and evaluate subsequent effects of a single AAV administration up to 6 months post-treatment, a timeline which more closely aligns with the time point at which modest clinical benefits of AAV-GDNF administration were observed in patients (*65*).

To the best of our knowledge, only two other studies have reported on AAV delivery in NHP with FUS. Blesa et al confirmed GFP expression in targeted regions in both healthy and Parkinsonian MPTP rhesus macaques after BBB opening with MRgFUS followed by systemic injection of AAV9-hSyn-GFP (*43*). Apart from the use of portable USgFUS for BBB opening, our study exhibits several key differences from the aforementioned study related to the timing and dose of AAV injection, and AAV construct. First, our study leveraged co-injection of MBs and AAV to capitalize on potential cavitation-mediated mechanisms for vector transport across the BBB during sonication, whereas the prior study injected AAV after confirmation of BBB opening on MRI, yielding a time difference between sonication and AAV administration of up to 2 hours. In addition to this, the prior study utilized a systemic dose of 5.0e13 gc/animal of AAV9-hSyn-GFP, compared to injection of ∼2.6e14 gc/animal of AAV9-CAG-GFP in the study presented herein. The combination of the increased dose of AAV9 administered *during* BBB opening and the use of the ubiquitous CAG promoter rather than the neuron-specific hSyn promoter yielded an apparent increase in overall transgene expression in the targeted regions relative to histological observations made in the prior study. Our study also offers additional insight into the transduction efficiency of AAV through quantification of vector DNA through ddPCR, confirming the effectiveness of USgFUS in targeted AAV transduction through both histological and biochemical analyses. Indeed, the prior study evaluates MRgFUS-mediated AAV delivery to the striatum in MPTP monkeys, attesting to the tolerability of systemic gene delivery and focal BBB opening in PD animal models, and provides critical translational insight for the field.

In addition to the aforementioned study, a second study by Parks et al, evaluated efficacy of focal transgene delivery with systemically-administered AAV2 and AAV9, with gene expression driven by hSyn, in marmosets (*44*). Consistent with our findings and the observations of Blesa and colleagues, Parks et al noted an increase in transgene expression in BBB-opened regions, with transgene expression in neurons due to the use of the hSyn promoter in AAV constructs. Parks et al also noted increased transduction efficiency with AAV9 relative to AAV2 after FUS-mediated BBB opening in marmosets, consistent with our previous findings in mice (*42*). The major difference between the study conducted by Parks et al and the study presented herein is the difference in NHP model. Marmosets are significantly smaller than rhesus macaques exhibiting much thinner skulls, which arguably do not recapitulate the thickness or microstructure of the human skull (*66*). Similarly to the study by Blesa et al., AAV was systemically administered after confirmation of BBB opening on MRI, which may have contributed to differences in observed transduction efficiency relative to our present study. Crucially, our study was designed to model multiple aspects of future potential FUS-mediated gene therapy techniques in humans, from transducer hardware design considerations for acoustic wave propagation through the human skull, to elucidating potential limitations related to the substantially higher titer of AAV needed for a systemic route of injection in patients.

Another primary difference between the study presented herein and previous studies investigating FUS-mediated AAV delivery in NHP lies in our use of ultrasound guidance during FUS or ThUS. Our study elucidated important advantages of USgFUS for BBB opening, from the direct association observed between SCDh derived from PAM with resulting vector DNA in the sonicated regions, to targeting advantages related to B-mode targeting capabilities inherently enabled by ultrasound imaging arrays. Our group has previously demonstrated an association between stable cavitation and BBB opening in NHP (*67–69*), but this study is the first to relate cavitation dose to AAV transduction efficiency in primates. This observed association is highly advantageous in that the amount of vector DNA to be expected within the FUS-targeted region could potentially be inferred non-invasively, during the treatment, without MRI confirmation. Indeed, future studies are needed to confirm this relationship, however this preliminary evidence underscores the advantages of acoustic monitoring for FUS-mediated AAV delivery. In addition to this, B-mode imaging prior to FUS and ThUS sonications reiterated the importance of normal beam-to-skull incidence angle to achieve optimal BBB opening efficiency (*70*). While we did not correct for sub-optimal incidence angle prior to sonication in this study, the observed relationship between symmetric skull geometry on B-mode imaging and BBB opening volume motivates additional investigation into the utility of anatomical ultrasound imaging integrated into USgFUS for ensuring maximal AAV delivery into sonicated brain regions. Our group has recently developed ultrasound-based strategies for skull imaging using harmonic ultrasound imaging which may be implemented in AAV delivery experiments in NHP to facilitate targeting in the future (*37*).

Some limitations of our present study merit additional discussion. First, only 2 NHP were used for this study, necessitating additional biological replicates to further validate our results, specifically a relationship between the efficacy of USgFUS-mediated AAV delivery and the particular brain region targeted. Other limitations relate to the viral method of gene therapy proposed in this study. NHP and humans alike commonly possess neutralizing antibodies (nAb) against particular AAV serotypes (*71–74*), limiting the applicability of serotype-specific AAV delivery for all patients for a particular gene therapy indication. The presence of nAb against AAV9 for example precluded observation of transduction in one NHP in the aforementioned study conducted by Blesa et al (*43*) emphasizing a limitation of both a systemic route of administration, and the use of AAV vectors in general. Despite this, tradeoffs between the low immunogenicity of AAV, size and packaging capabilities relative to other viral vectors such as lentivirus (LV) or adenovirus (ADV) still make it a primary choice for gene therapy in the clinic (*75*).

Future studies are primarily aimed at improving the specificity of gene delivery induced by FUS and AAV, both increasing expression in neurons of interest and reducing peripheral gene expression which could lead to systemic toxicity. In addition, future studies are aimed at assessing the feasibility of USgFUS-mediated gene delivery with other naturally-occurring, as well as engineered AAV capsids given the specific benefits related to tissue targeting and limiting immune response exhibited by particular AAV serotypes other than AAV9. Finally, we aim to implement non-invasive USgFUS-mediated AAV delivery in additional experimental models of neurodegenerative diseases to evaluate expression of a functional transgene given the feasibility for such an approach demonstrated with ThUS-mediated AAV9-hNTRN delivery in MPTP mice.

## CONCLUSION

FUS and gene therapy have each demonstrated significant potential for enabling non-invasive and efficacious neurodegenerative disease therapy in the clinic. However, there is still a need for pre-clinical research to answer questions related to their optimal synergistic utility. In this study, we offer insights to these unanswered questions relating to quantification of brain transduction resulting from USgFUS-mediated BBB opening and the relationship between FUS and ThUS parameters on gene delivery efficacy. By understanding how much AAV delivery occurs within the brain after systemic injection and targeted delivery with the parameters reported herein, along with revealing the vital association between image-derived cavitation dose and gene delivery within the NHP brain for the first time, FUS continues to emerge as a promising strategy to significantly aid in gene therapy pre-clinical and clinical studies. Specifically, alternative options posed by portable USgFUS systems in gene therapy administration in clinical trials may offer greater reach to potential candidates averse to surgical administration. This approach not only broadens the pool of eligible participants but also addresses a significant barrier to enrollment in trials that involve surgical interventions, particularly in regard to the elderly patient population most afflicted by neurodegenerative disease. As a result, USgFUS could play a crucial role in advancing the development and adoption of gene therapies for conditions like Parkinson’s disease by making these trials more accessible and appealing to a wider population. With future studies aimed at improving the targeting capabilities and safety profile of this combinatorial technique in both the therapeutic ultrasound and gene therapy spaces, USgFUS could enable accessible, non-invasive, and therapeutically efficacious gene therapy to help minimize the immense burden of neurodegenerative disease treatment on patients and their families.

## MATERIALS AND METHODS

### Experimental Design

The overall objectives of this study were to evaluate the feasibility and provide a multifaceted and quantitative readout of AAV9 delivery across the BBB in mice and rhesus macaques with two modalities of portable low-intensity therapeutic ultrasound: conventional single-element, spherically-focused FUS, or a multielement, point-of-care ThUS array. The ThUS configuration constitutes a simplified yet flexible design which could enable increased accessibility of therapeutic ultrasound for non-invasively treating neurological disorders, which motivated the final objective of the study herein, which was to determine whether ThUS-facilitated AAV9 delivery could elicit disease-modifying therapeutic effects in a mouse model of early PD. To investigate these objectives, 3 studies were designed. Study 1 constituted an AAV9-CAG-GFP dose escalation study in mice with FUS to inform the systemic AAV9 dose for a subsequent feasibility study using both FUS and ThUS with the same AAV9-CAG-GFP construct in NHP, referred to herein as Study 2. Finally, Study 3 was conducted to evaluate ThUS+AAV9 delivery for therapeutic neurorestoration in MPTP mice and offer future research direction for therapeutic ultrasound-facilitated gene delivery in treating neurodegenerative diseases.

All procedures involving animals were performed in accordance with approved protocols under the guidelines of the Columbia University Institutional Animal Care and Use Committee (IACUC). The terminal NHP study (Study 2) was initiated after n=2 geriatric NHPs (29-30 y.o., 12.8-13.0 kg) housed within the Columbia University Institute of Comparative Medicine (ICM) were recommended for euthanasia due to arthritis and other complications minimizing their tolerance for anesthetic procedures: NHP A and NHP B. Advantageously, the progressed age of the NHPs allowed us to recapitulate transcranial therapeutic ultrasound-facilitated AAV9 delivery through the aged skull and brain in an elderly human population most likely to be afflicted with neurodegenerative disorders, while projected euthanasia permitted institutional approval of this terminal study. Before study initiation, both NHPs were also screened for AAV9 neutralizing antibodies (nAb). Each NHP received one intravenous injection of AAV9 but were euthanized at different time points post-AAV dosing (3 weeks vs. 4 weeks). A summary of the individual experimental details for the study timeline of both NHPs is shown in Figure 1. An additional adult male rhesus macaque NHP C, was used for ThUS parameter optimization and targeting confirmation as discussed later on, did not receive AAV and was not euthanized at the terminal study endpoint with NHP A and NHP B.

Both NHP A and NHP B were placed on an immunosuppressive steroid regimen consisting of once-daily, orally administered prednisolone (1 mg/kg) beginning 14 days prior to FUS- or ThUS-mediated AAV delivery and ending on the day of euthanasia, 21-32 days post-sonication. Prior to AAV delivery, baseline CT and MRI scans were acquired for numerical acoustic wave simulations (*49*) and for evaluation of pre-sonication MR signatures before BBB opening procedures for target selection before dosing. On the day of AAV delivery, BBB opening using either FUS or ThUS was performed according to the protocol in the following section. After sonications were completed and the NHPs were transferred to the MRI suite, a series of anatomical MRI were acquired within two hours after BBB opening to evaluate both safety and efficacy of BBB opening with ThUS and FUS. Following MRI, the NHPs were closely monitored during recovery from anesthesia by veterinary staff. After a period of 3-4 weeks (21 days for NHP A, and 32 days for NHP B) to allow transduction and gene expression to occur, NHPs were euthanized, and tissues were dissected and collected for biodistribution and immunohistochemistry assays as described in later sections.

Prior to conducting this study, a dose escalation study with the same AAV9-CAG-GFP construct was conducted in wild-type mice to inform AAV9 dosing for the NHPs. A total of 25 8-10-week-old, female C57BL/6J mice (The Jackson Laboratory, Bar Harbor, ME, USA) were allocated for this study and were divided into the following 5 groups: 1.0e10 gc/mouse IV AAV9+FUS, 1.0e11 gc/mouse IV AAV9+FUS, 5.0e11 gc/mouse IV AAV9+FUS, 5.0e11 gc/mouse IV AAV9 only, and a sham group receiving anesthesia only. Mice received FUS and/or AAV9 on day 0, followed by a 21-day survival period before euthanasia by transcardial perfusion with ice-cold 1X PBS. Excised brains were split along the midline such that one hemisphere was allocated for paraffin embedding and IHC to observe cell-type specificity of GFP expression, while the hippocampus was dissected from the opposite hemisphere for quantification of AAV9 DNA and RNA with ddPCR. Liver tissue from each mouse was also split for IHC and ddPCR analyses.

An additional study (Study 3) employing ThUS-mediated AAV9-hSyn-hNTRN-WPRE delivery in the subacute MPTP mouse model was conducted to elucidate the utility of a next-generation therapeutic ultrasound system to elicit neurorestoration in MPTP mice. The timeline for this study is shown in Figure 8A. A total of 42 male C57BL/6J mice (Charles River Laboratories, Kingston, NC, USA) were obtained at 12 weeks of age and were acclimated to isolated housing facilities for 4 weeks before MPTP dosing. The acute model of MPTP-induced neurodegeneration is extremely sensitive to environmental variations between animals and was thus generated according to the specific procedures set forth by Jackson-Lewis and Przedborski et al. (*47*). 34 mice underwent daily IP injections of 1-methyl-4-phenyl-1,2,3,6-tetrahydropyridine (MPTP) for 5 days, while 8 mice underwent control saline injections on the same dosing schedule. After a 21-day period of neurodegeneration. Mice were randomly split into five experimental groups: MPTP only, MPTP+ThUS, MPTP+AAV, MPTP+ThUS+AAV, and saline-injected healthy controls. Within each experimental group, approximately half of the animals were allocated to be euthanized for analysis 3 months post-treatment, while the other half was allocated for euthanasia 6 months post-treatment. After euthanasia by transcardial perfusion with ice-cold 1X PBS, mice brains were excised and prepared for TH immunohistochemistry and image analysis performed by investigators blinded to experimental conditions.

### BBB opening procedure in NHP

A schematic of the experimental apparatus used for FUS-mediated BBB opening in NHP is shown in Figure 1A, with an overall timeline of the study presented in Figure 1E. NHPs were anesthetized by injection of ketamine and dexmedetomidine followed by maintenance of the anesthetic plane with a mixture of isoflurane and oxygen. The head was fixed in a stereotactic instrument before shaving and depilating hair on the scalp to facilitate proper acoustic coupling between the transducer and scalp. A layer of degassed ultrasound gel was placed atop the head below a degassed water bath designed to easily facilitate transducer positioning from target to target. After initialization and registration of the neuronavigation system used to provide real-time guidance during FUS or ThUS (Brainsight, Rogue Research Inc., Montréal, Québec, Canada), targeting was achieved using a robotic arm (UR5e, Universal Robots, Odense, Denmark). B-mode imaging was conducted to ensure normal incidence of the FUS beam to the skull before acquiring baseline cavitation activity immediately prior to injection of microbubbles for each target. During the BBB opening procedure, NHPs received a bolus intravenous (IV) injection of polydisperse microbubbles followed by immediate sonication and injection of 2.0e13 gc/kg of AAV9-CAG-GFP after a rise in cavitation level was observed with PAM or PCI as described in later sections. NHP A underwent BBB opening along 4 separate trajectories summarized in Figure 1E, while NHP B underwent a single FUS sonication, followed by AAV injection during the first of three ThUS sonications with the experimental configuration shown in Figure 1B at distinct targets sonicated in the order displayed in Figure 1F.

NHP A underwent FUS-mediated BBB opening targeted to the following brain regions in chronological order of sonication: left putamen, left caudate, left hippocampus, and right substantia nigra. A single bolus injection of house-made polydisperse microbubbles was injected immediately prior to the bolus injection of AAV9-CAG-GFP and commencement of sonication at the left putamen for 2 minutes. The transducer was then translated 5 mm to the right before immediate sonication of the left caudate for another 2 minutes with the previously injected microbubbles. The transducer was then positioned to target the left hippocampus before a second bolus injection of microbubbles and sonication for 2 minutes. Lastly, the transducer was positioned at the right substantia nigra before a final bolus injection of microbubbles and sonication for 2 minutes.

NHP B underwent FUS-mediated BBB opening targeted to the right hippocampus for 2 minutes, followed by the bolus injection of AAV9-CAG-GFP and opening of the left caudate and putamen using the rapid alternating steering angles (RASTA) pulse sequence for ThUS (Supplementary Figure 6A) to target both the caudate and putamen simultaneously for 4 minutes. The ThUS transducer was then targeted to the left substantia nigra before sonication using a single ThUS focus for 2 minutes, followed by a final ThUS sonication targeted to the left brainstem for 2 minutes, again using a single ThUS focus.

### Microbubbles

In-house-manufactured microbubbles were synthesized and activated according to previously published protocols (*76, 77*). In brief, 1,2-distearoyl-sn-glycero-3-phosphocholine (DSPC, Avanti Polar Lipids Inc., Alabaster, AL, USA) and 1,2-distearoyl-sn-glycero-3-phosphoethanolamine-N-[methoxy(polyethylene glycol)-2000] (DSPE-mPEG2000, Avanti Polar Lipids Inc., Alabaster AL, USA) were combined at a 9:1 molar ratio. The combined mixture of lipids was dissolved in 50 mL of solution containing filtered phosphate-buffered saline, glycerol, and propylene glycol at an 8:1:1 ratio and immersed in a sonication bath for 1 to 2 hours until complete dissolution of lipids. After aliquoting 1.5 mL of the solution into individual 3 mL vials for storage, microbubbles used on the day of sonication underwent activation via 5 alternating sequences of air aspiration and perfluorobutane (PFB) gas infusion (Decafluorobutane, FluoroMed L.P., Round Rock, TX, USA). Following PFB gas infusion, the vial was activated via a modified amalgamator (VialMix^TM^, Lantheus Medical Imaging, N. Billerica, MA, USA) for 45 s. Concentration and size distribution were determined with a particle counter (MultiSizer 4e, Beckman Coulter, Indianapolis, IN) before normalization according to NHP weight for Study 2. 1.02e10 MBs (NHP A) to 1.04e10 MBs (NHP B) were administered for each bolus injection in NHP, in a concentration ranging from 0.033-0.083 mL/kg. For the mice studies (Study 1 and Study 3), 5 µL of MBs were administered for each injection at a concentration of 8e8 MBs/mL.

For target confirmation experiments in NHP C. 2 vials of FDA-approved LUMASON microbubbles (Bracco Diagnostics Inc., Princeton NJ, USA) were used as described in the previous section. Manufacturer estimated microbubble concentrations ranged from 1.50e8 – 5.6e8 MBs/mL, with a mean diameter range of 1.5-2.5 µm.

### Adeno-associated viruses (AAV)

The AAV9-CAG-GFP vectors employed in these studies were provided by REGENXBIO Inc. (study 1 and study 2 NHP A) and Spark Therapeutics Inc. (study 2 NHP B). A systemic dose of 2.0e13 gc/kg, corresponding to a total injected dose of 2.56e14 – 2.60e14 gc for each NHP was carefully diluted in an IV formulation buffer before injection via an IV port in the right saphenous vein. For Study 3, AAV9-hSyn-hNTRN-WPRE was manufactured and underwent quality control protocols (Vector Biolabs, Malvern, PA, USA) at a titer of 4.0e13 gc/mL. AAV was diluted in sterile saline in preparation for an intravenously injected dose of 1.1e11 gc/animal.

### Statistical analysis

All statistical analyses were performed in Prism (Ver. 10.3.1, GraphPad), where statistical tests and significance criteria are specified within relevant figure captions. Unless otherwise mentioned, *\*p*<0.05, *\*\*p*<0.01, *\*\*\*p*<0.001, *\*\*\*\*p*<0.0001, and error bars denote ± one standard deviation of the mean.

## List of Supplementary Materials

Supplementary Materials and Methods

Supplementary Results

Supplementary Discussion

Fig. S1 to S9

Table S1 to S2

Videos 1-2

## Acknowledgements

The authors wish to thank the following veterinary personnel from the Columbia University Institute of Comparative Medicine (ICM) for support during NHP studies: Melissa Tamimi, Dominik Hajosi, Felicia Teller, Emily Zhang, Brenton Mays, Marise Rodriguez, Aram Safarov, and Olena Shevchenko. The authors would also like to thank the following from Spark Therapeutics Inc. for study and experimental support: Ashley Carter, Mohamad Nayal, Renee Gentzel, Drew Peterson, Graciela Rivera-Pena, Charlie Li, Jodi McBride, and Melinda Peters. The authors wish to also thank Vernice Jackson-Lewis for her deep expertise in working with the MPTP mouse model. Some figures were created with Biorender.com

## Funding

This study was funded in part by the National Institutes of Health grants R01EB009041, R01AG038961, R56AG038961 [EEK], and R01MH133020 [EEK & VPF], REGENXBIO Inc., Spark Therapeutics Inc., the Naval Information Warfare Center (NIWC), the Defense Advanced Research Projects Agency (DARPA) under Contract No. N66001-19-C-4020 [EEK, VPF], and the Focused Ultrasound Foundation. LSM was performed with support from the Zuckerman Institute’s Cellular Imaging Platform and the National Institutes of Health (NIH 1S10OD023587-01).

## Author Contributions

Conceptualization: AJB, RJ, EEK, JBS, EAE, CTC, BSH

Methodology: AJB, RJ, SB, FNT, SJG, NK, CTC,

Investigation: AJB, RJ, SB, FNT, SJG, NK, SLG, DT, RLN, JB, DJ, MD, SAD, FBK, JC, CTC, YD, VPF, JBS

Visualization: AJB, RJ, SB, SJG, CTC, YD, GDG

Funding acquisition: EEK, VPF, JBS, EAE, OD, ER

Project administration: AJB, RJ, EEK, JBS, EAE, AR

Supervision: EEK, JBS, EAE, AR, ER, OD, VPF, SP

Writing – original draft: AJB, RJ

Writing – review & editing: AJB, RJ, FNT, AR, EEK, CTC, YD, GDG, EAE, ER, BSH

## Competing Interests

Some of the methodology presented herein is supported by patents optioned to Delsona Therapeutics, Inc. where EEK serves as co-founder and scientific adviser. The authors also disclose that REGENXBIO Inc. and Spark Therapeutics Inc. have contractual agreements exclusively with Columbia University and no other third parties.

## Data and materials availability

Data produced for this study can be made available upon reasonable request to the corresponding authors.

