## Supplementary Material for "Portable transcranial therapeutic ultrasound enhances targeted gene delivery for Parkinson’s disease: from rodent models to non-human primates"

This document contains:

Supplementary Materials and Methods

Supplementary Results

Supplementary Discussion

Supplementary Figures S1-S9

Supplementary Tables S1-S2

Supplementary Video Captions 1-2

**Supplementary Materials and Methods:**

Numerical acoustic simulations

Acoustic wave simulations were performed to estimate the focal volume and pressure attenuation at each target for each NHP. Simulations were performed using the acoustic package of the k-wave toolbox (GPU-optimized) in MATLAB *(78, 79)*, where the heterogeneous 3D maps of density and sound speed, and homogeneous acoustic absorption of the cranial muscle, skull, and brain were derived from pre-sonication CT scans *(49)* for input onto the computational grid with 7 and 12 ppw (points per wavelength) resolution for the working frequencies of 500 kHz (ThUS) and 250 kHz (FUS), respectively, ensuring simulation convergence and accuracy by contemplating at least 8 grid points per average NHP skull thickness *(37)*. Sound speed (*c)*, density (ρ) and absorption (α) of water (background), skull, brain and muscle tissues were: *c_water_* = 1473 m/s, ρ_water_ = 1000 kg/m^3^, α_water_500kHz_ = 150e-6 dB/cm, α_water_250kHz_ = 9e-6 dB/cm, c_skull_max_=4112 m/s, c_skull_mean_=2463 m/s, ρ_skull_max_=3323 kg/m^3^, ρ_skull_mean_ = 1851 kg/m3, α_skull_500kHz_ = 1.35 dB/cm, α_skull_250kHz_ = 0.68 dB/cm, c_brain_ = 1560 m/s, ρ_brain_ = 1000 kg/m^3^, α_brain_500kHz_ = 0.41 dB/cm, α_brain_250kHz_ = 0.19 dB/cm, c_muscle_ = 1600 m/s, ρ_muscle_=1000 kg/m^3^, α_muscle_ = 0.75 dB/cm.

BBB opening sequences

Given the difference in skull thickness and geometry between mice and NHP, and the objective to evaluate translational applicability of therapeutic ultrasound systems for clinical use, two different configurations for each therapeutic ultrasound modality, i.e. FUS or ThUS, were employed. For each configuration, *in situ* acoustic pressure was estimated with a combination of water tank experiments using ex-vivo skulls and a hydrophone, and 3D transcranial acoustic wave propagation simulations (Supplementary Figures 1-2). A major difference between the FUS and ThUS configurations employed in these studies was the pulsing paradigm, where BBB opening was induced with either long pulses on the order of milliseconds, or short pulses on the order of microseconds, with particular advantages and disadvantages outlined in the discussion section. For the purposes of the studies reported herein, FUS was operated with a long-pulse sequence, while ThUS was operated with a short-pulse sequence with the parameters reported in Supplementary Tables 1-2.

*1.5 MHz single-element long-pulse USgFUS (mice)*

The FUS system for BBB opening in mice consisted of a 1.5 MHz single-element, geometrically focused transducer (Imasonic, Voray-sur-l’Ognon, France), with a 6.7 ms pulse length, 10 Hz PRF and an in-situ PNP of 650 kPa (0.53 MI). 4 total 60-second sonications were performed per BBB opening session to target the entire hippocampus as described in previously published work by our group *(17)*. 5 uL bolus injections of house-manufactured polydisperse microbubbles were injected prior to the 1^st^ and 3^rd^ sonication, where AAV9-CAG-GFP was co-injected with microbubbles for animals receiving AAV9+FUS.

*1.5 MHz multi-element short-pulse ThUS (mice)*

The ThUS system for BBB opening and treatment monitoring with PCI in mice consisted of a single repurposed diagnostic imaging phased array (P4-1, ATL, Philips, Amsterdam, Netherlands) operated at a 1.5 MHz center frequency by a research ultrasound system (Vantage 256, Verasonics Inc. Kirkland, WA) with the following transmit parameters: 1.0 MPa PNP (0.82 MI), 5-cycle pulse length, 1000 Hz PRF, 0.5 Hz, BRF. The transducer was positioned such that central focal coordinates were 2.0 mm anterior and 1.5 mm mediolateral relative to lambda and refined using B-mode imaging as described in previously published studies *(31)*. Immediately after initiating the ThUS RASTA pulse sequence, 1.1e11 gc/mouse of AAV9-hSyn-hNTRN-WPRE was intravenously co-injected with a 5 uL bolus of microbubbles at the concentration specified in a later section, while the ThUS sonication continued for 2 minutes.

*0.25 MHz single-element long-pulse USgFUS (NHP)*

BBB opening with FUS was achieved using a long-pulse sequence (Figure 1C) transmitted from a single-element clinical 0.25 MHz FUS transducer (H-231, 110 mm outer diameter, 44 mm inner diameter, 6 x 6 x 49 mm focal volume, Sonic Concepts, Bothell Washington) with a central hole containing a confocally-aligned imaging array (P4-1, 2.5 MHz, 96 elements, ATL, Philips, Amsterdam, Netherlands) for B-mode imaging and passive acoustic mapping (PAM) *(16)*. Each FUS sonication consisted of 10 ms pulses, repeated at a PRF of 2 Hz for a sonication duration of 2 minutes at an estimated in situ PNP of 0.4 MPa (MI of 0.8). The same formulation and concentration of house-made polydisperse microbubbles as mentioned previously was injected according to the weight of the NHP on the day of sonication.

*0.50 MHz multi-element short-pulse ThUS (NHP)*

Synchronous BBB opening and PCI with ThUS in NHP was achieved using a custom short-pulse sequence (Figure 1D) operated by a Verasonics Vantage research ultrasound system, transmitted by a custom 0.5 MHz multielement linear array (32 elements, 4 x 16 x 40 mm focal volume, Vermon, Tours, France). Both a single ThUS focus and two steered foci (ThUS RASTA) were employed in this study, where bursts of 100 3-cycle pulses repeated at a PRF of 1000 Hz were deployed at a BRF of 0.5 Hz for a sonication duration of 2 minutes, or 4 minutes for ThUS RASTA with an estimated in situ PNP of 1.0 MPa (MI of 1.4). The same formulation and concentration of house-made polydisperse microbubbles as mentioned previously was injected according to the weight of the NHP on the day of sonication.

During experiments used to confirm targeting with ThUS in NHP C where no AAV was delivered, slight variations to the parameters and microbubble formulations were chosen to evaluate feasibility of BBB opening with commercially-available LUMASON (Bracco Diagnostics Inc., Princeton NJ, USA) microbubbles. The pulse length was increased to 5 cycles rather than 3 cycles, and two sonications were performed: the first targeted to the caudate (Supplementary Figure 8A-C) using the manufacturer’s recommended dilution factor for preparation of 5 mL sterile saline in 1 vial of microbubbles, and the second targeted to a volume containing the lateral geniculate nucleus (LGN), hippocampus, and posterior putamen (Supplementary Figure 8D-F), using a greater concentration of microbubbles using a dilution factor for preparation of 1 mL of sterile saline in 1 vial of microbubbles. Each sonication lasted for approximately 4 minutes until the cavitation dose derived from intraprocedural PCI (Supplementary Figure 8G,I) approached pre-sonication levels (Supplementary Figure 8H,J).

Magnetic resonance imaging (MRI)

For the mice studies (Study 1 and Study 3), contrast-enhanced T_1_-weighted MRI for confirmation and quantification of BBB opening were acquired according to the exact protocol described in previously published work from our group *(17, 19, 31, 39)*. Approximately 30 min after FUS or ThUS, mice received an 0.2 mL IP injection of gadodiamide (Omniscan®, GE Healthcare, Chicago, IL, USA) before imaging in a 9.4 T vertical bore MRI system (Ascend, Bruker Medical, Billerica, MA, USA) for imaging with a T_1_-weighted 2D FLASH sequence (TR: 230 ms, TE: 3.3 ms, Flip angle: 70°, 6 averages, FOV: 25.6 mm × 25.6 mm, Matrix size: 256 × 256, Slice thickness: 0.4 mm, Resolution 0.1 mm × 0.1 mm).

For the NHP studies (Study 2), safety and BBB opening evaluation were performed on a wide-bore 3T MRI scanner (SIGNA^TM^ Premier, GE Healthcare, Chicago, IL, USA) approximately 1 hour post-FUS/ThUS. BBB opening was evaluated using contrast-enhanced T_1_-weighted MRI with the following sequence parameters after an intravenous injection of gadodiamide contrast agent (0.2 mL/kg, Omniscan®, GE Healthcare, Chicago, IL, USA): TR: 7.44 ms, TE: 3.11 ms, averages: 0.7, flip angle: 11°, resolution: 0.5 x 0.5 mm, slice thickness: 0.8 mm. T_2_ FLAIR MRI with the following sequence parameters were acquired prior to contrast administration to evaluate safety after the BBB opening procedure: TR: 7500 ms, TE: 93.93 ms, averages: 1, flip angle: 90°, resolution: 0.41 x 0.41 mm, slice thickness: 0.99 mm. BBB opening was quantified by subtraction of the 1^st^ T_1_-weighted MRI acquisition post contrast injection from the 5^th^, and automatically delineated by regions where the mean intensity within a manually-selected BBB opening ROI was greater than that of the surrounding tissue within a 98% confidence interval according to a previously reported method *(16)*.

Tissue preparation

At the endpoint of study 1, mice were deeply anesthetized using a mixture of isoflurane and oxygen before euthanasia by transcardial perfusion with ice-cold 1X PBS. Brains were extracted and split along the midline to allocate one hemisphere for fixation in 4% PFA, cryoprotection in 30% sucrose, and IHC as described in the following section, and the other for hippocampal dissection and snap-freezing for AAV biodistribution analysis with ddPCR. Livers were also extracted for both IHC and biodistribution analyses.

For study 2, both NHPs underwent similar tissue collection and processing protocols. Each NHP was deeply anesthetized with a mixture of isoflurane and oxygen before thoracotomy and transcardial perfusion with ice-cold 1X PBS until the effluent ran clear and tissues appeared visibly blanched, indicating complete perfusion. Gross tissues were collected from the periphery and were sampled using 3 mm tissue punches. The excised brains were sliced coronally into 4-6 mm-thick slabs according to post-sonication contrast-enhanced T_1_-weighted MRI. Tissue punches were taken within the sonicated regions and contralateral control regions including the cortex, putamen, caudate, substantia nigra, hippocampus and lateral geniculate nucleus, along with additional brain regions of interest to evaluate background AAV9 transduction including the choroid plexus, spinal cord, cerebellum, medulla, and pre-frontal cortex. Tissue punches (3 mm diameter, 4 mm thickness) were immediately snap frozen on dry ice and stored at -80 ℃ before quantification of AAV genome copies per cell using ddPCR. Slabs allocated for histology were either stored in 4% PFA for 48 hours and transferred to 30% sucrose for cryosectioning or fixed in 10% NBF for 24 hours followed by processing for FFPE and microtome sectioning of paraffin-embedded sections.

Tissue punches (3 mm diameter, 4 mm thickness) of the following peripheral organs were acquired to evaluate systemic gene transduction as a result of systemic AAV injection: liver, heart, skeletal muscle, pancreas, lung, spleen, and kidney. Tissue punches were also snap frozen and stored at -80 ℃ for quantification of AAV genome copies per cell using ddPCR. A slab of liver tissue was also fixed with 4% PFA for 48 hours and transferred to 30% sucrose for cryosectioning and evaluation of GFP expression with fluorescence microscopy.

At either the 3-month or 6-month endpoint for study 3, mice were deeply anesthetized using a mixture of isoflurane and oxygen before euthanasia by transcardial perfusion with ice cold 1X PBS. Brains were extracted and fixed in 4% PFA for 48 h before storage in 30% sucrose with 0.01% sodium azide at 4 ℃ until sectioning.

Immunohistochemistry

For study 1, the mouse brain hemispheres allocated for ICH were prepared according to previously published protocols *(80)*. First, hemispheres were embedded in OCT (Tissue Tek, Torance, CA, USA), sectioned into 50-µm-thick slices using a freezing microtome, and stored in a solution containing 0.045 M phosphate buffer, 30% ethylene glycol, and 25% glycerol before staining. Sections were first washed 3X with Tris-buffered saline (TBS) before blocking for 1 h in a solution containing TBS and 3% normal horse serum (Vector Laboratories, Newark, CA, USA) and 0.25% Triton-X (TBS+). Sections were incubated with the following primary antibodies in TBS+ at 4 ℃ for 48 hours: chicken anti-GFP (#NB100-1614, Novus Biologicals, Centennial, CO, USA) at 1:1000, rabbit anti-Iba1 (#E404W, Cell Signaling Technologies, Danvers, MA, USA) at 1:500, mouse anti-NeuN (#ab104224, Abcam, Cambridge, MA, USA) at 1:1000, and rabbit anti-S100β (#ab52642, Abcam, Cambridge, MA, USA) at 1:1000. After incubation in primary antibodies, sections were washed twice with TBS for 15 min each followed by TBS+ for 30 min before incubation in the following secondary antibodies for 2 hours at room temperature: donkey anti-chicken IgY Alexa Fluor 488 (Invitrogen, Waltham, MA, USA), donkey anti-mouse IgG Alexa Fluor 555 (Invitrogen, Waltham, MA, USA), and donkey anti-rabbit IgG Alexa Fluor 647 (Invitrogen, Waltham, MA, USA), each at a dilution of 1:250. Finally, sections were incubated with DAPI (Invitrogen, Waltham, MA, USA) at 5 µg/mL for 5 minutes, washed 3 more times with TBS for 15 min each, and mounted onto slides with coverslips and mounting media (ProLong Gold anti-fade).

For study 2, slabs allocated for cryosectioning were frozen and sliced into 35-µm-thick floating coronal sections within the BBB opening and contralateral brain regions. Slabs allocated for microtome sectioning were paraffin-embedded and sectioned into 5-um coronal sections mounted directly to glass microscope slides. For NHP A, IHC was conducted in an analogous manner to the above description for study 1. For NHP B, paraffin embedded brain slabs were coronally sectioned into 5-micron thick slices and were immunostained for GFP, S100β, GFAP, and NeuN. The IHC protocol was performed as follows. Sections were baked in the ACD HybEZ^TM^ II Oven (Biotechne 321710) for 1 hour at 60°C to melt the paraffin, deparaffinized in Histo-Clear (#50-899-90147, Fisher Scientific, Waltham, MA, USA), rehydrated through a 100%, 95%, 70%, and 50% ethanol series, and rinsed in distilled water. Heat-induced epitope retrieval was performed using pH 6 citrate buffer (BioSB BSB 0020) in a TintoRetriever Digital Pressure Cooker (BioSB BSB 7008) at 100°C with low pressure setting for 20 minutes. Subsequently, the slides were cooled for 30 min at room temperature and washed three times in 1X PBS, 5 minutes each. Tissue sections were then incubated in blocking solution (5% BSA, 1% donkey serum, and 0.2% Triton X-100 in 1X PBS) for 1 hour at room temperature. Primary antibodies against GFP (1:1000, #ab6673, Abcam, Waltham, MA, USA), GFAP (1:2000, # AB5541, MilliporeSigma, Burlington, MA, USA), NeuN (1:500, #ab104224, Abcam, Waltham, MA, USA), and S100β (1:2000, #ab52642, Abcam, Waltham, MA, USA) were applied in the blocking solution overnight at 4℃. The next day, the sections were washed once in 1X PBS containing 0.2% Triton X-100 and then twice in 1X PBS, for a total of three washes, 10 minutes each. Secondary antibodies were applied in the blocking solution for 2 hours at room temperature. The following secondary antibodies were used: donkey anti-goat 555 (1:500, #A32816, ThermoFisher, Waltham, MA, USA), donkey anti-rabbit 488 (1:300, #A32790, ThermoFisher, Waltham, MA, USA), donkey anti-mouse 647 (#A32787, ThermoFisher, Waltham, MA, USA), and donkey anti-chicken 790 (1:500, #703-655-155, Jackson ImmunoResearch, West Grove, PA, USA). After removing the secondary antibodies, the sections were washed in 1X PBS containing 0.2% Triton X-100 followed by washes in 1X PBS as described above, and autofluorescence blocking was performed for 3 minutes using Vector® TrueVIEW® Autofluorescence Quenching Kit (Vector Labs SP-8400-15) as per manufacturer’s instructions. Immediately after, the sections were rinsed in distilled water and mounted with ProLong Gold with DAPI (#P36931, ThermoFisher, Waltham, MA, USA).

For study 3, excised brains stored in 30% sucrose were coronally cryosectioned into 35 µm-thick sections in the substantia nigra and striatal regions. Immunostaining for tyrosine hydroxylase (TH) for evaluation of neurorestoration was performed according to the following protocol: briefly, sections were washed 3x with 1X PBS followed by blocking for 1 h with 5% donkey serum and 0.3% Triton X-100. Sections were incubated overnight with TH anti-rabbit primary antibody (#Ab 657012, Sigma-Aldrich, St. Louis, MO, USA). Sections were washed again before incubation in AlexaFluor-594 donkey anti-rabbit secondary antibody solution in the dark for 1 h. Stained sections were washed a final time, mounted onto slides and coverslipped with DAPI mounting solution before imaging.

Microscopy and image analysis

For study 1, mounted coronal brain sections were imaged for luminance quantification with a Leica M205 FCA at 1x magnification with a 500 ms exposure time. Visualization of colocalization of GFP with cell type-specific markers including Iba1, NeuN, and S100β was performed with a confocal microscope (LSM 900 Airyscan 2, Zeiss, Oberkochen, Germany) at either 20X or 63X magnification. For study 2, imaging was performed for NHP A using the above microscopy apparatuses. For NHP B, images were acquired using a slide scanner (Axio Scan Z7, Zeiss, Oberkochen, Germany) with a 20X air objective at a resolution of 0.69 microns per pixel.

For study 3, fluorescence microscopy images for TH-stained coronal sections containing the substantia nigra and caudate putamen regions were acquired using a tile scan at 10x magnification on an Olympus microscope (BX61, Olympus Corporation, Tokyo, Japan) configured with a 647 nm filter cube. Imaging was conducted for all analyzed sections using the same exposure parameters for consistent fluorescence quantification. Images of TH-stained coronal brain sections were analyzed with a custom image processing pipeline in MATLAB and ImageJ. Images of 3 sections per mouse per brain region (centered at approximately -3.40 mm and +0.14 from bregma for the SN and CPu, separated by ± 1 increment of 35 µm) were background-normalized using histogram normalization after converting to grayscale. In a semi-automated script, the SNc and SNr were individually segmented before thresholding to isolate TH+ neuron cell bodies and dendritic processes. The areas of TH+ cell bodies and dendritic processes within the segmented SNc and SNr, defined as the maximum 75% of the luminance range in the normalized, thresholded images were quantified and normalized to the area of segmentation. An analogous procedure was conducted with fluorescence microscopy images of the striatum, except the criteria for determining TH-positivity was reduced to the top 50% of the luminance range due to the decrease in contrast relative to the SN images. Presented images were pseudo-colored in ImageJ for enhanced contrast and visualization of TH-stained neurons.

Biodistribution assays

Droplet digital polymerase chain reaction (ddPCR) was performed for tissue samples in study 1 and NHP A in study 2, whereas qPCR was performed for tissue samples from NHP B in study 2. For ddPCR, samples were homogenized using a tissue homogenizer (Precellys Evolution, Bertin Technologies, Rockville, MD, USA) in a lysis buffer containing RA1 (Macherey-Nagel, Allentown, PA, USA) and 1% β-mercaptoethanol. DNA extraction from tissue homogenates was performed according to the manufacturer’s instructions (DNeasy Blood & Tissue Kit, Qiagen, Germantown, MD, USA). Extracted DNA was combined with Naica multiplex PCR mix, probes, and primers for GFP, mouse or rhesus macaque glucagon, and mouse or rhesus macaque TATA-binding protein (TBP), and was loaded into Sapphire Chips (Stilla Technologies, Beverly, MA, USA) for PCR. Sample partitioning and the PCR thermal cycling program was performed by the Naica Geode instrument (Stilla Technologies, Beverly, MA, USA) before image acquisition on the Naica Prism3 reader (Stilla Technologies, Beverly, MA, USA) with the following exposure times: 50 ms for the blue channel, and 150 ms for the green channel. Total droplet enumeration and quality control was performed with the Crystal Reader software (Stilla Technologies, Beverly MA, USA) through detection of the FITC reference dye in the blue channel. Droplet-specific fluorescence values were analyzed using the Crystal Miner software (Stilla Technologies, Beverly, MA, USA), where GFP copy number was measured in the blue channel (FAM), and glucagon or TBP copy numbers were measured in the green channel (VIC).

For NHP B, tissue punch samples (3 mm) were collected from peripheral organs and brain slabs (4 mm thick). Tissue samples were homogenized on the Tissuelyser II (Qiagen, Germantown, MD, USA) in lysis buffer with 5 mm stainless steel beads. Following a 30-minute incubation at 56 C with shaking, tissue lysates were transferred to the QIAsymphony SP instrument for DNA extraction with the QIAsymphony DSP DNA Mini Kit (Cat # 937236, Qiagen, Germantown, MD, USA). Sample concentrations were measured with a nanodrop. qPCR of DNA samples was performed on the QuantStudio 7 Pro System (ThermoFisher) using primers and a TaqMan probe targeting EGFP. Quantification of vector genomes was performed using a standard curve generated with linearized plasmid DNA.

Passive acoustic mapping (PAM) and power cavitation imaging (PCI) processing

PAM images acquired during USgFUS sonication were reconstructed as described previously *(16, 81)*. Briefly, real-time PAM images were displayed during sonication using coherence-factor-based PAM with GPU acceleration (RTX A6000, NVIDIA, Santa Clara, CA, USA). Final reconstructed PAM images were generated by averaging individual PAM frames after the microbubble injection for each sonication. Stable harmonic cavitation doses (SCDh), and stable ultraharmonic cavitation doses (SCDu) were calculated as described in the aforementioned studies, and included the 3^rd^ through 6^th^ harmonic and ultraharmonic frequencies. Inertial cavitation dose (ICD) was computed as the sum of the averaged amplitude of frequencies between each harmonic to ultraharmonic interval for each burst, and below the 3^rd^ harmonic extending to 75 kHz.

PCI acquired during ThUS were reconstructed using a GPU (Quadro P5000, NVIDIA, Santa Clara, CA, USA) to generate the final PCI maps displayed in Figure 5 and Supplementary Figures 6-9, by summing the frames for each burst after the microbubble injection as described previously *(31)*. Cavitation dose, i.e. PCI signal intensity was defined as the sum of pixel intensities within the final PCI map for each sonication.

iDISCO tissue clearing, whole-brain immunolabeling, and light sheet microscopy

A brain from an MPTP mouse in study 3 treated with ThUS+AAV which was euthanized 6 months post treatment underwent whole-brain tissue clearing and TH immunolabeling with iDISCO. Briefly, the previously fixed brain was dehydrated over a 20%-100% methanol gradient and bleached overnight using a solution of dichloromethane (DCM) and methanol. The brain was then washed before incubating in permeabilization solution containing glycine DMSO for 24 hours at 37 ℃ and blocked with a gelatin solution containing porcine skin and Triton X for another 24 hours. The brain was then incubated in a heparinized solution containing TH primary antibody (Ab 657012, Sigma-Aldrich) at a concentration of 1:1000 for 11 days before washing, dehydrating and incubation in secondary antibody (AlexaFluor-647) for 8 days. After immunolabeling, the brain was washed, rehydrated and incubated in DCM before clearing in dibenzyl ether (DBE). The entire tissue clearing process was conducted over a period of approximately 4 weeks.

Light sheet microscopy (LSM) was conducted on a single brain to visualize the entire intact nigrostriatal pathway after iDISCO tissue clearing and immunostaining using an Ultramicroscope II Light Sheet Microscope (Miltenyi Biotec, Bergisch Gladbach, Germany) with a **2X** 0.5 NA Olympus VMPLAPO Plan Apochromat Objective. The brain was silicon-mounted before being placed in an imaging cuvette containing DBE and imaged horizontally with a 3.4 um-thick light sheet at a 640 nm excitation wavelength.

**Supplementary Results:**

Given the success of ThUS RASTA to elicit multi-target BBB opening in mice in prior studies, we extended the ThUS RASTA sequence for BBB opening volume expansion in NHP B with analogous pulsing parameters (Supplementary Figure 6A). The ThUS array was positioned with the lateral dimension of the array parallel to the sagittal plane of the NHP brain rather than targeting two hemispheres simultaneously. Steering angles were chosen to minimize skull incidence angle effects, and the center of each steered focal volume were separated by 5 mm as shown in Supplementary Figure 6B. Pre-sonication B-mode imaging was conducted after targeting the ThUS array along a trajectory passing through the anterior caudate and posterior putamen to ensure a 90° beam to skull incidence angle. PCI acquired during the sonication depicted a substantial rise in cavitation activity with nearly the same average magnitude along both the anterior trajectory and posterior trajectory, denoted by the red and blue traces in Supplementary Figure 6F, respectively. Summed PCI over the 4-minute sonication duration agreed spatially with BBB opening volumes denoted by the white contours registered and superimposed onto the final summed PCI in Supplementary Figure 6G, yielding a final BBB opening volume of 78.13 mm^3^. In a separate sonication targeting the same coordinates within the striatum, BBB opening volume was reduced by nearly 4-fold due to poor skull incidence relative to the aforementioned sonication. Supplementary Figure 6H demonstrates the asymmetrical skull reflection in pre-sonication B-mode imaging which yielded substantial reflections in therapeutic transmits denoted by the high PCI signal intensity at the skull in Supplementary Figure 6I. Relative to the PCI maps from the aforementioned sonication shown in Supplementary Figure 6E, this sonication exhibited reduced PCI signal intensity within the brain, corroborated by reduced PCI signal intensity in the volume sonicated by the anterior steering angle (Supplementary Figure 6K). The mismatch between cavitation induced by the two steering angles yielded an overall 1.6-fold decreased cavitation dose over the 4-minute sonication, and reduced BBB opening volume of 19.65 mm^3^ as depicted by the white contour in Supplementary Figure 6L, relative to the sonication presented in Supplementary Figure 6C-G. These results emphasize the importance of proper targeting before sonication with ThUS, and the potential role of PCI to enable adjustments to the sonication protocol during BBB opening in future experiments.

During targeting confirmation experiments in NHP C which did not receive AAV, BBB opening was confirmed after a single ThUS sonication within the caudate and anterior putamen comprising a volume of 134.16 mm^3^ (Supplementary Figure 8A-C), and after a second sonication within a 303.52 mm^3^ volume containing the LGN, hippocampus, and posterior putamen (Supplementary Figure 8D-F). Most notably, contrast enhancement was detected within the dentate gyrus in the hippocampus, which could not be successfully targeted with the 0.25 MHz FUS configuration due to poor beam incidence angle. This indicates that the footprint and dimensions of the linear ThUS array may yield increased flexibility in targeting deep structures within the NHP brain. Additionally, the feasibility of inducing BBB opening with commercially-available, FDA-approved LUMASON microbubbles is advantageous for regulatory approval of ThUS-mediated gene delivery in the clinic.

**Supplementary Discussion**

Our neurorestoration observations in study 3 are consistent with results of other previously published studies investigating the effects of the GDNF family of neurotrophic factors on amelioration of lesions caused by MPTP or 6-OHDA. While we specifically investigated neurorestorative effects in this study, meaning that treatment interventions were conducted after lesioning by MPTP, other studies have also observed neuroprotective effects by inducing overexpression of NTF prior to lesioning. Kearns et al. revealed that GDNF administration at most 6 hours before 6-OHDA injection conferred complete neuroprotective effects *(57)*. Tomac et al., conducted one of the first studies investigating the impact of GDNF administration on reversal of MPTP-induced neurodegeneration in mice, where both neuroprotection with GDNF administration prior to MPTP dosing, and neurorestoration when injecting GDNF after MPTP dosing was observed *(58)*. While these aforementioned studies were conducted with GDNF, a protein within the same family of NTF as NTRN, Oiwa et al. specifically demonstrated both neuroprotection of TH+ neurons in the substantia nigra from 6-OHDA lesioning, and neurorestoration of TH+ neurons in the striatum 4 weeks after NTRN administration *(59)*. Given the disparity between the relatively short timepoints for restoration evaluation in prior studies and the results presented herein, we conclude that at least 3-6 months of NTRN overexpression may be necessary to observe significant changes in neuronal rescue in both the substantia nigra and striatum in mice. Beyond rodent models, Kells et al., demonstrated significant neurorestorative effects in Parkinsonian NHP dosed with MPTP after CED of AAV2-GDNF to the striatum, noting retrograde transport from the striatum to the SN as both a mechanism for initiating neurorestoration, and as a limitation of the striatal gene delivery target *(60)*.

It is important to note that in our study, and in many of the aforementioned studies, TH staining was agnostic to subtypes of neurons implicated in PD. While TH+ dopaminergic neurons make up the excitatory pathway stemming from the SNc to the striatum, TH+ GABAergic neurons within the SNr contribute to the balance of inhibitory signals which maintain proper firing of dopaminergic neurons in the SNc *(61)*. Alterations in the complex interplay between these types of neurons contributes to the motor dysfunction observed in PD patients and must be further investigated in gene delivery studies to assess the impact of neurorestorative therapy on motor function. To further understand the differential impact of AAV9-hSyn-hNTRN delivery to different neuronal subtypes susceptible to transduction by the hSyn promoter, AAV enhancers may be used to increase specificity for targeting particular neuronal subtypes *(62, 63)*.

**Supplementary Figures (Figure S1-S9)**

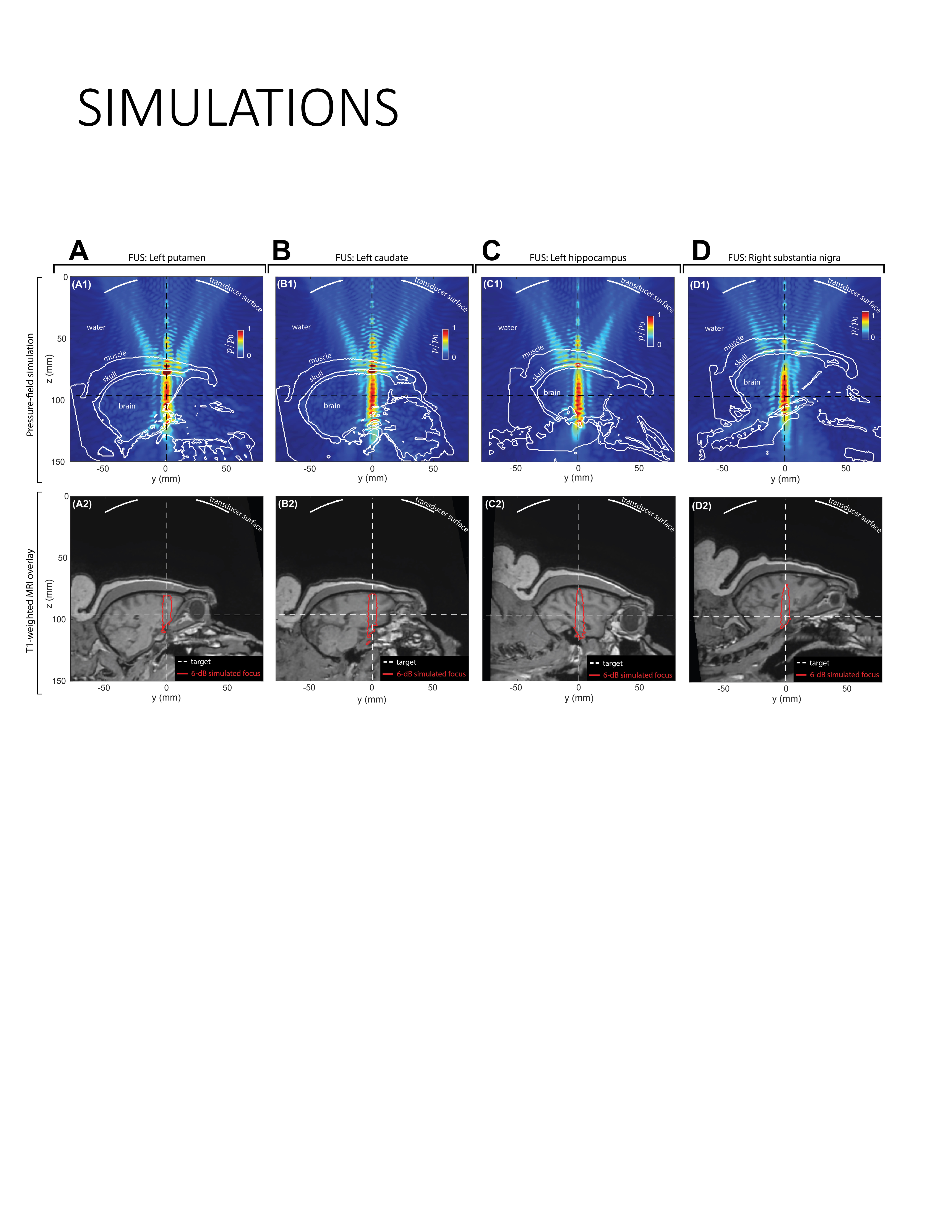

**Supplementary Figure 1: Acoustic wave simulations for FUS treatment planning in NHP A. A)** Computational grid depicting FUS focal pressure map through the intact primate skull at the left putamen trajectory (top). The intersection of the black dashed lines denote the center of the FUS focus. The estimated -6 dB FUS focal dimensions overlaid onto the sagittal slice of the pre-FUS MRI is shown in red, while the white crosshairs denote the center of the focus (bottom). **B)** Simulated FUS pressure map along the left caudate trajectory (top) with -6dB focal contour overlaid onto the pre-FUS MRI (bottom). **C)** Simulated FUS pressure map along the left hippocampus trajectory (top) with -6dB focal contour overlaid onto the pre-FUS MRI (bottom). **D)** Simulated FUS pressure map along the right substantia nigra trajectory (top) with -6dB focal contour overlaid onto the pre-FUS MRI (bottom).

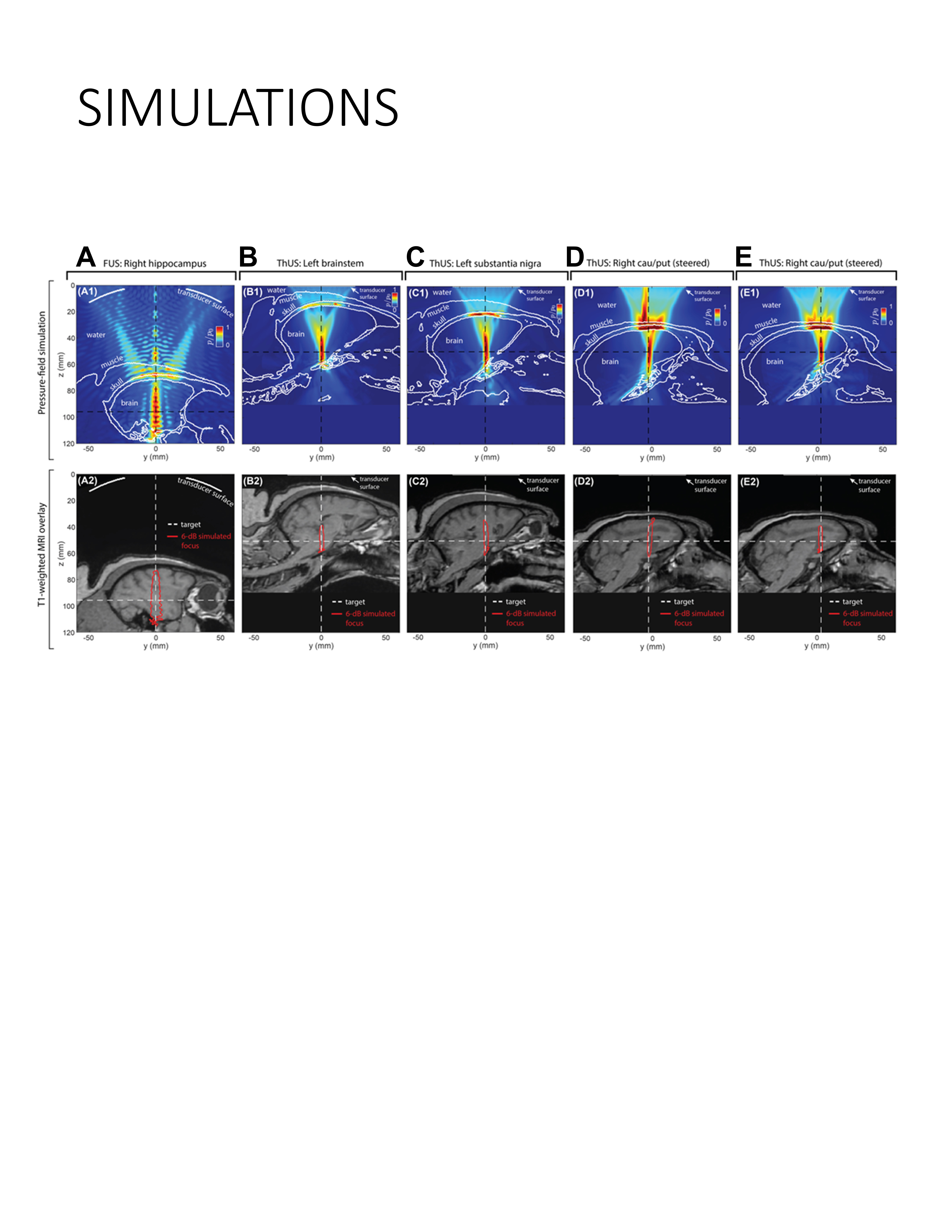

**Supplementary Figure 2: Acoustic wave simulations for FUS and ThUS treatment planning in NHP B. A)** Computational grid depicting FUS focal pressure map through the intact primate skull at the right hippocampus trajectory (top). The intersection of the black dashed lines denote the center of the FUS focus. The estimated -6 dB FUS focal dimensions overlaid onto the sagittal slice of the pre-FUS MRI is shown in red, while the white crosshairs denote the center of the focus (bottom). **B)** Simulated ThUS pressure map along the left brainstem trajectory (top) with -6dB focal contour overlaid onto the pre-FUS MRI (bottom). **C)** Simulated ThUS pressure map along the left substantia nigra trajectory (top) with -6dB focal contour overlaid onto the pre-FUS MRI (bottom). **D)** Simulated ThUS pressure map along the posterior angle of the right caudate/putamen trajectory (top) with -6dB focal contour overlaid onto the pre-FUS MRI (bottom). **E)** Simulated ThUS pressure map along the anterior angle of the right caudate/putamen trajectory (top) with -6dB focal contour overlaid onto the pre-FUS MRI (bottom).

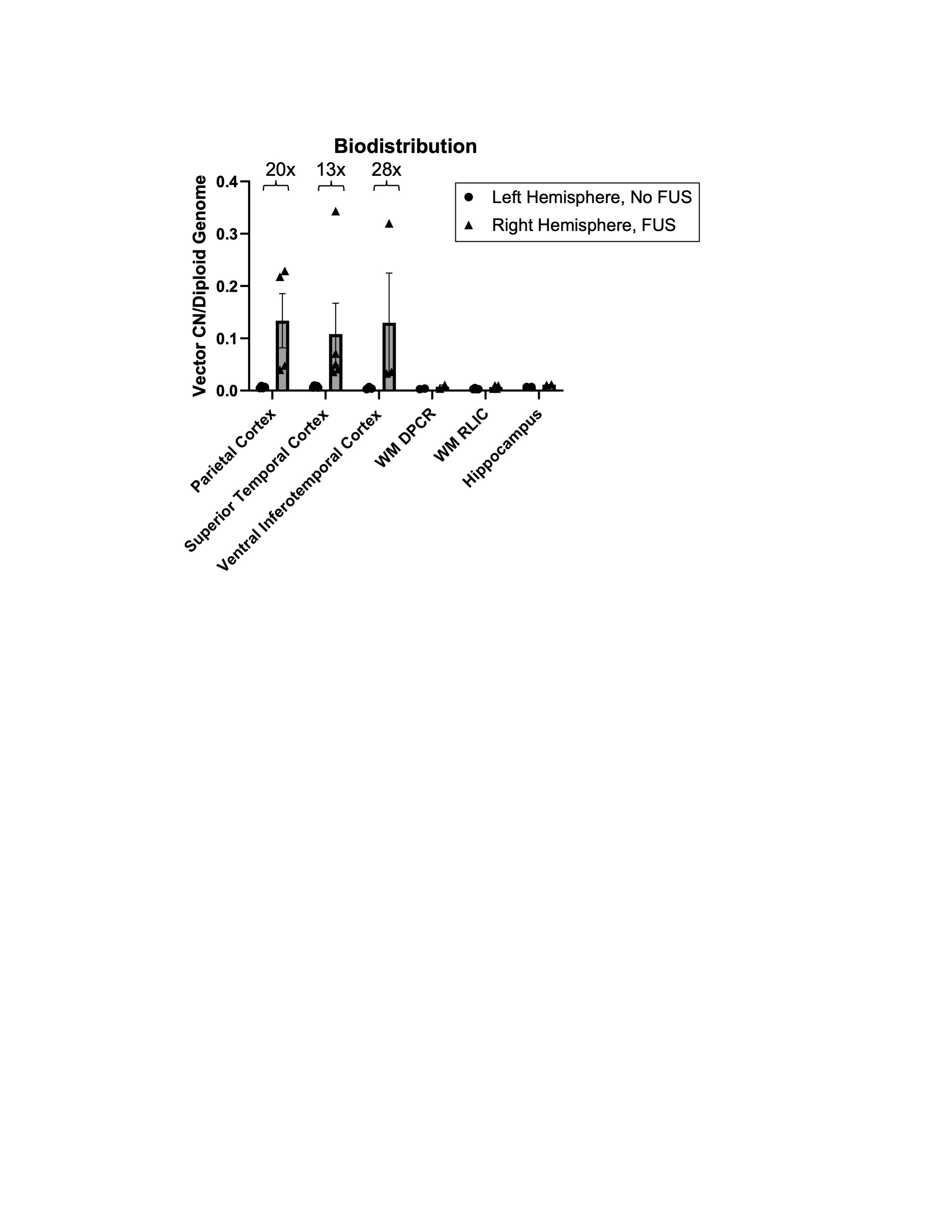

Supplementary Figure 3: Cortical AAV biodistribution in NHP B. Increased vector genome copy number (CN) per cell in cortical areas sonicated with FUS. DPCR = dorsal posterior corona radiata; RLIC = retrolenticular limb of the internal capsule. Error bars represent standard deviation of the mean of n=2-5 qPCR samples per region.

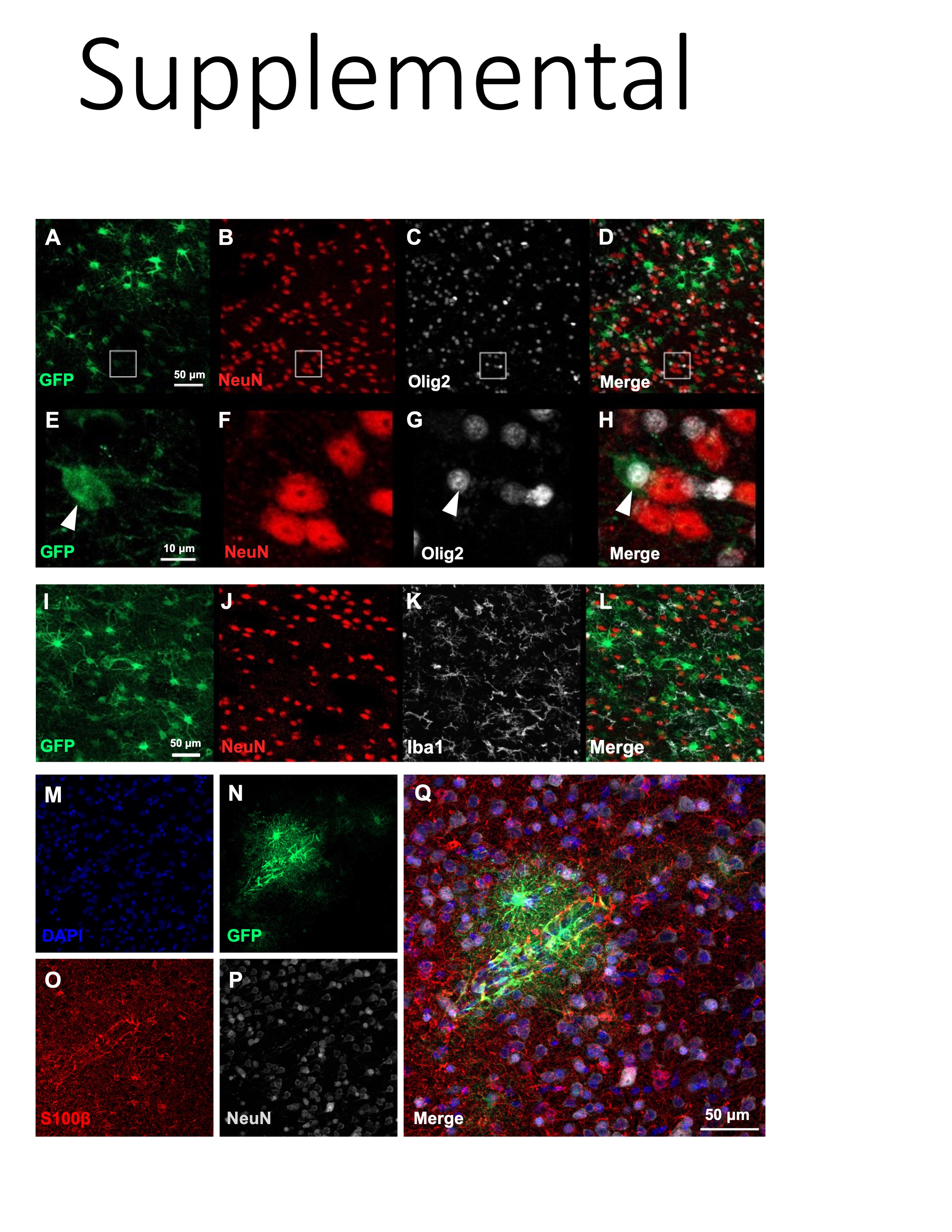

Supplementary Figure 4: AAV9-CAG-GFP transduction of other cerebral cell types in NHP A with FUS. A) GFP, B) NeuN, C) Olig2, and D) merged channel image with corresponding enlarged images from rectangular ROIs shown in E-H. I) GFP, J) NeuN, K) Iba1, and L) merged channel image devoid of GFP expression within microglia. M) DAPI, N) GFP, O) S100ß, P) NeuN, and Q) merged channel image depicting unique transduction pattern of astrocyte-vascular coupling.

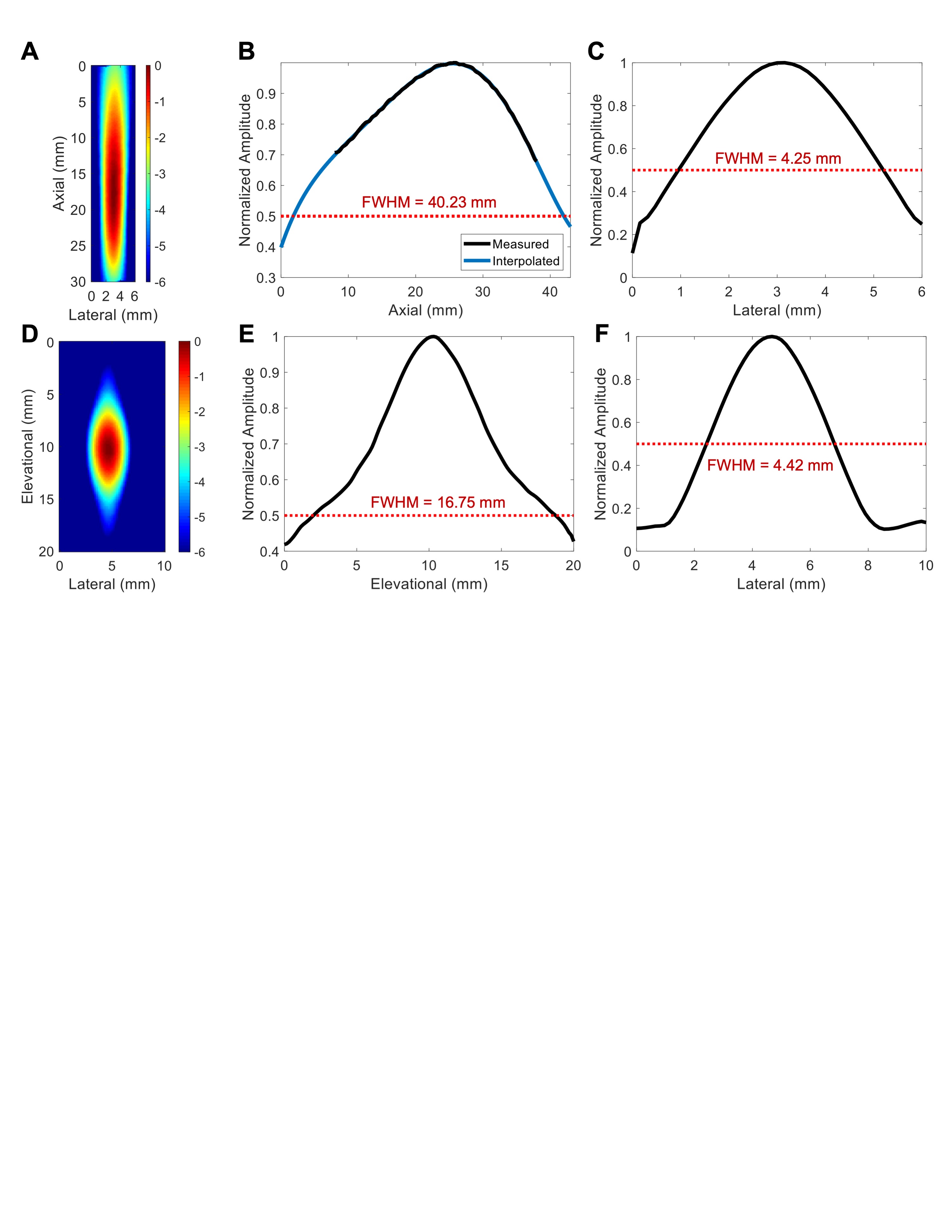

Supplementary Figure 5: Focal profiles of 32-element 500 kHz ThUS array at 55 mm depth. A) 2D axial beam plot thresholded at -6dB. B) Axial and C) lateral 1D focal profiles obtained from (A). D) 2D horizontal focal profile in the elevational/lateral dimension thresholded at -6 dB. E) Elevational and F) lateral 1D focal profiles obtained from (D).

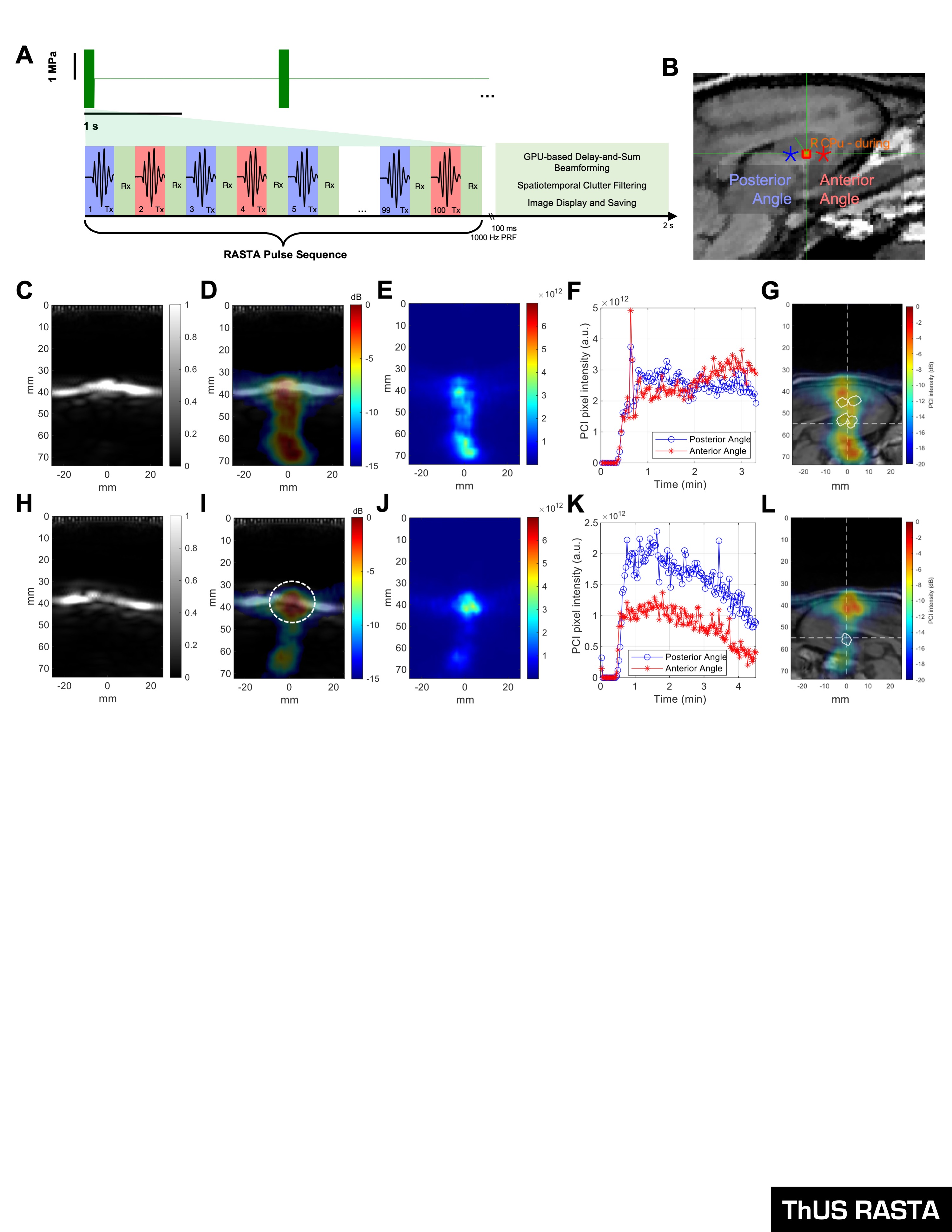

Supplementary Figure 6: Bi-focal BBB opening with ThUS RASTA in NHP. A) Schematic of the ThUS RASTA pulse sequence used in NHP. B) Diagram of focus locations denoted as colored asterisks superimposed onto sagittal brain MRI during target selection. Targets separated by 5 mm in sagittal dimension. Representative ideal ThUS RASTA sonication with C) pre-sonication B-mode, D) self-normalized PCI superimposed onto B-mode, E) absolute PCI, F) trace of PCI intensity over the sonication duration, and G) BBB opening contours overlaid onto PCI registered with sagittal MRI slice at the center of the BBB opening volume. Another sonication targeting the same region with poor beam incidence angle effects with H) pre-sonication B-mode, I) self-normalized PCI superimposed onto B-mode with ellipse dashed ROI denoting poor beam to skull incidence angle, J) absolute PCI, K) trace of PCI intensity over the sonication duration, and L) BBB opening contours overlaid onto PCI registered with sagittal MRI slice at the center of the BBB opening volume.

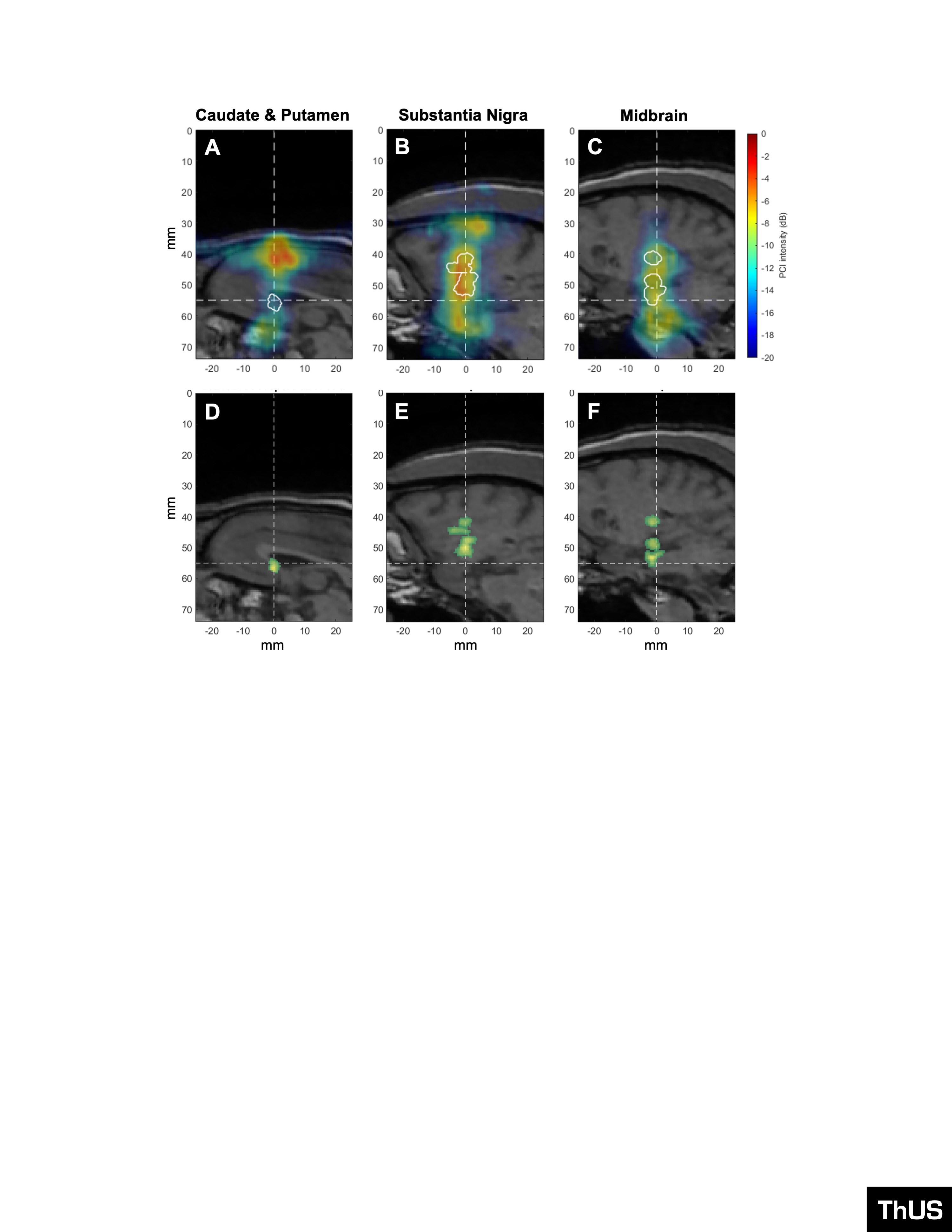

Supplementary Figure 7: PCI and BBB opening volumes corresponding to targeted AAV9-CAG-GFP delivery with ThUS. PCI (colored overlays) and BBB opening contour (white outlines) superimposed onto central T1-weighted MRI slice after sonication with A) ThUS RASTA targeted to the caudate & putamen, B) a single ThUS focus targeted to the substantia nigra, and C) a single ThUS focus targeted to the midbrain. BBB opening volumes denoted by the green overlays for D) the caudate & putamen target (V_BBB_ = 19.61 mm3), E) substantia nigra target (V_BBB_ = 78.65 mm3), and F) the midbrain (V_BBB_ = 65.02 mm3). Dashed lines indicate the center of the planned ThUS focus after targeting. Note that the PCI and BBB opening shown in (A) and (D) correspond to the sonication depicted in Figure 2H-L.

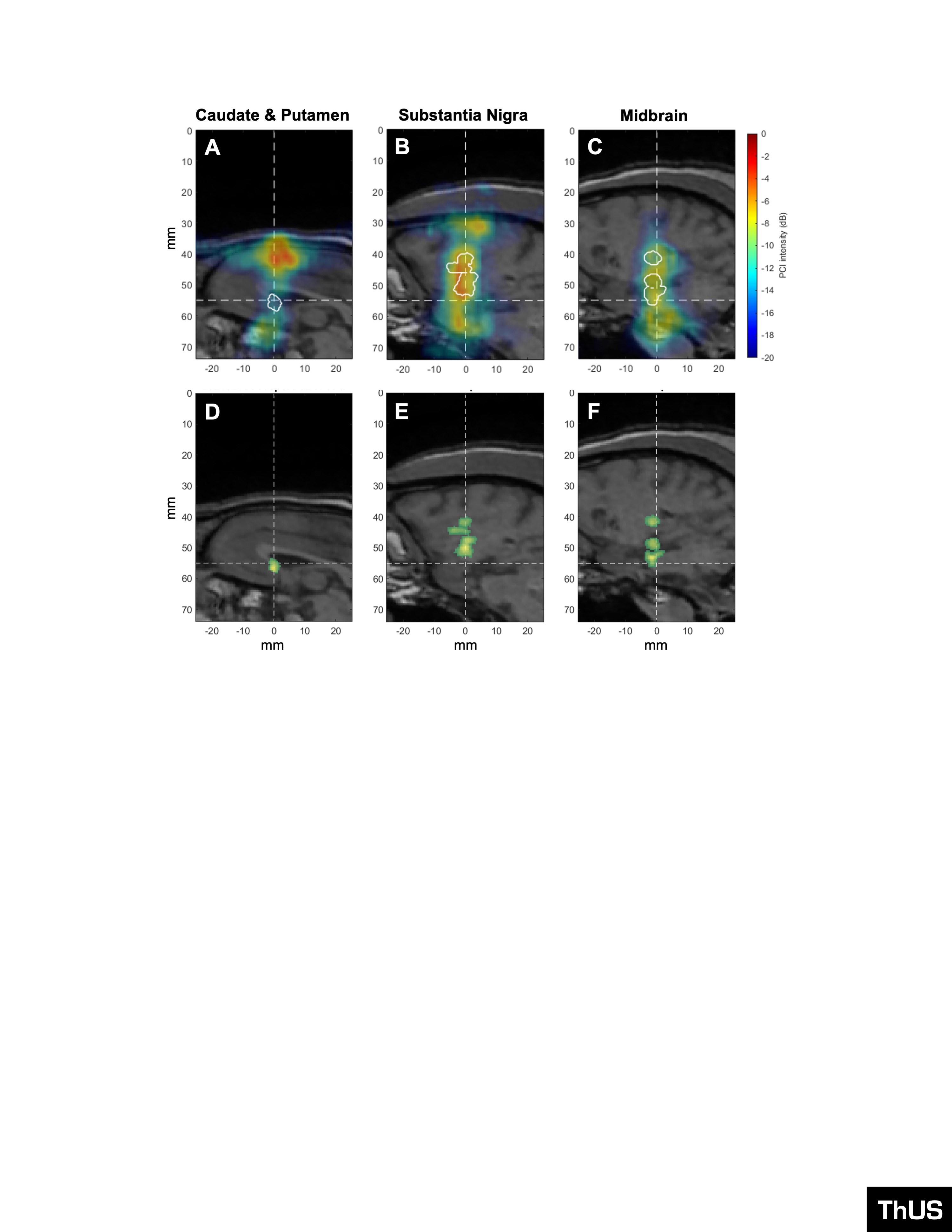

Figure 586: PCI and BBB opening volumes corresponding to targeted AAV9-CAG-GFP delivery with ThUS. PCI (colored overlays) and BBB opening contour (white outlines) superimposed onto central T1-weighted MRI slice after sonication with A) ThUS RASTA targeted to the caudate & putamen, B) a single ThUS focus targeted to the substantia nigra, and C) a single ThUS focus targeted to the midbrain. BBB opening volumes denoted by the green overlays for D) the caudate & putamen target (V_BBB_ = 19.61 mm3), E) substantia nigra target (V_BBB_ = 78.65 mm3), and F) the midbrain (V_BBB_ = 65.02 mm3). Dashed lines indicate the center of the planned ThUS focus after targeting. Note that the PCI and BBB opening shown in (A) and (D) correspond to the sonication depicted in Figure 2H-L.

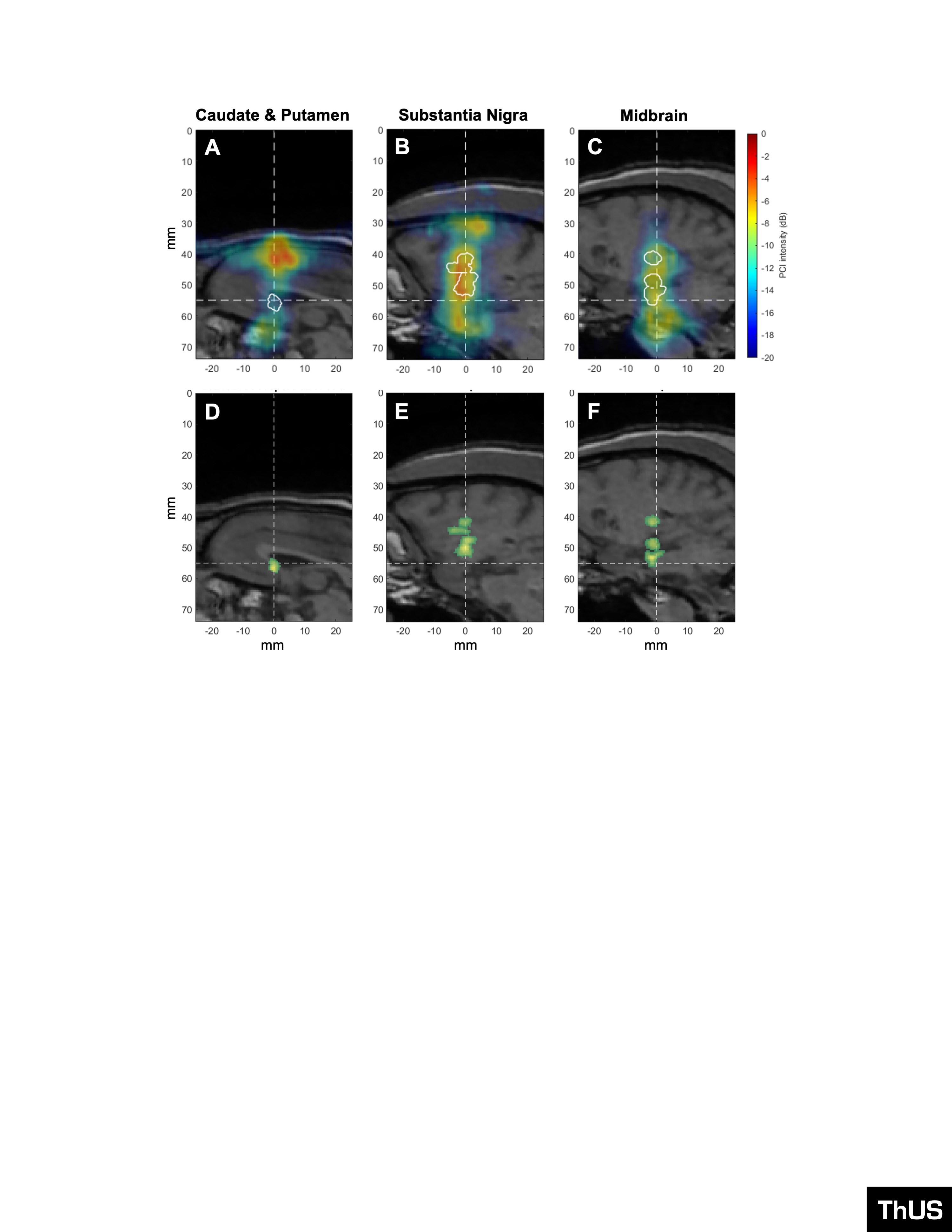

Figure 587: PCI and BBB opening volumes corresponding to targeted AAV9-CAG-GFP delivery with ThUS. PCI (colored overlays) and BBB opening contour (white outlines) superimposed onto central T1-weighted MRI slice after sonication with A) ThUS RASTA targeted to the caudate & putamen, B) a single ThUS focus targeted to the substantia nigra, and C) a single ThUS focus targeted to the midbrain. BBB opening volumes denoted by the green overlays for D) the caudate & putamen target (V_BBB_ = 19.61 mm3), E) substantia nigra target (V_BBB_ = 78.65 mm3), and F) the midbrain (V_BBB_ = 65.02 mm3). Dashed lines indicate the center of the planned ThUS focus after targeting. Note that the PCI and BBB opening shown in (A) and (D) correspond to the sonication depicted in Figure 2H-L.

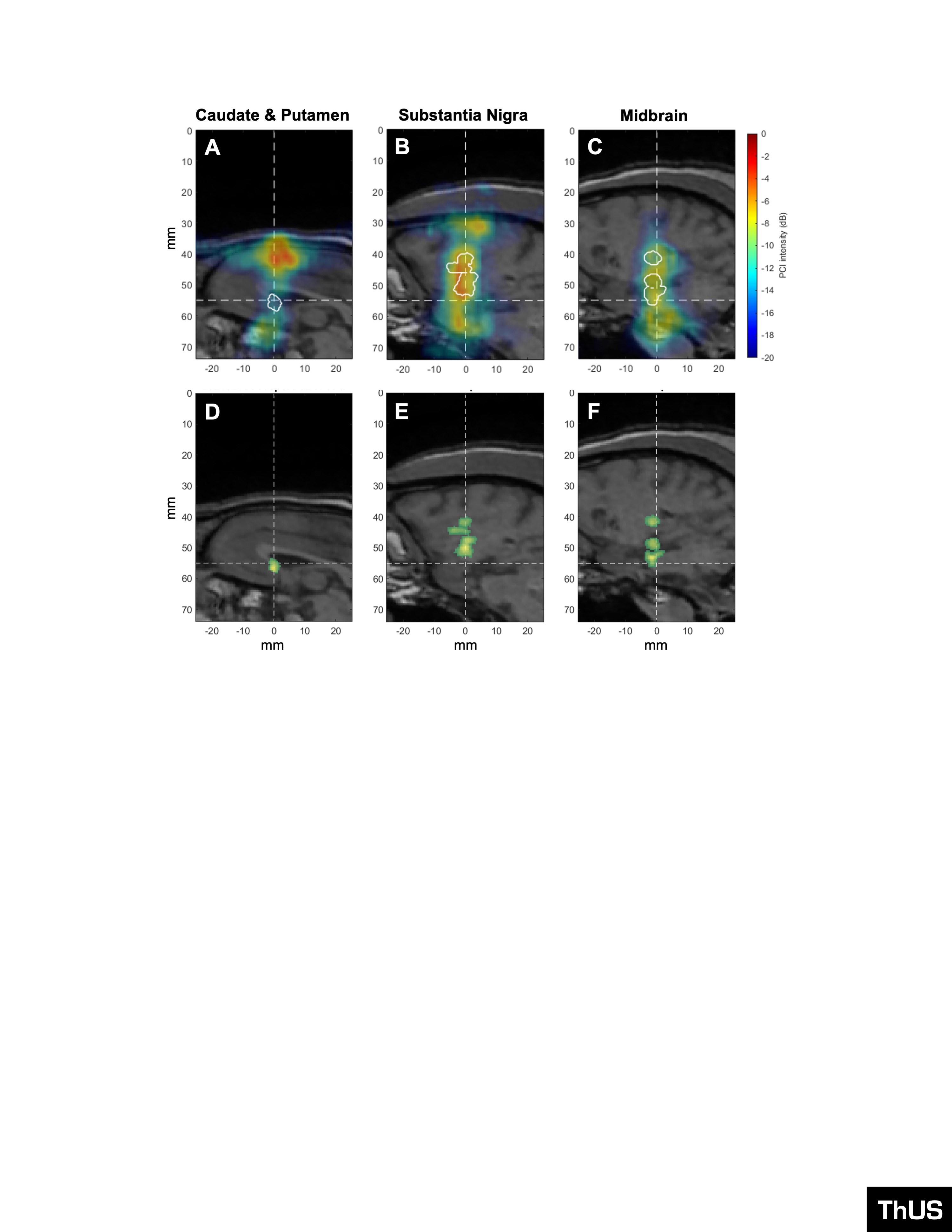

Figure 588: PCI and BBB opening volumes corresponding to targeted AAV9-CAG-GFP delivery with ThUS. PCI (colored overlays) and BBB opening contour (white outlines) superimposed onto central T1-weighted MRI slice after sonication with A) ThUS RASTA targeted to the caudate & putamen, B) a single ThUS focus targeted to the substantia nigra, and C) a single ThUS focus targeted to the midbrain. BBB opening volumes denoted by the green overlays for D) the caudate & putamen target (V_BBB_ = 19.61 mm3), E) substantia nigra target (V_BBB_ = 78.65 mm3), and F) the midbrain (V_BBB_ = 65.02 mm3). Dashed lines indicate the center of the planned ThUS focus after targeting. Note that the PCI and BBB opening shown in (A) and (D) correspond to the sonication depicted in Figure 2H-L.

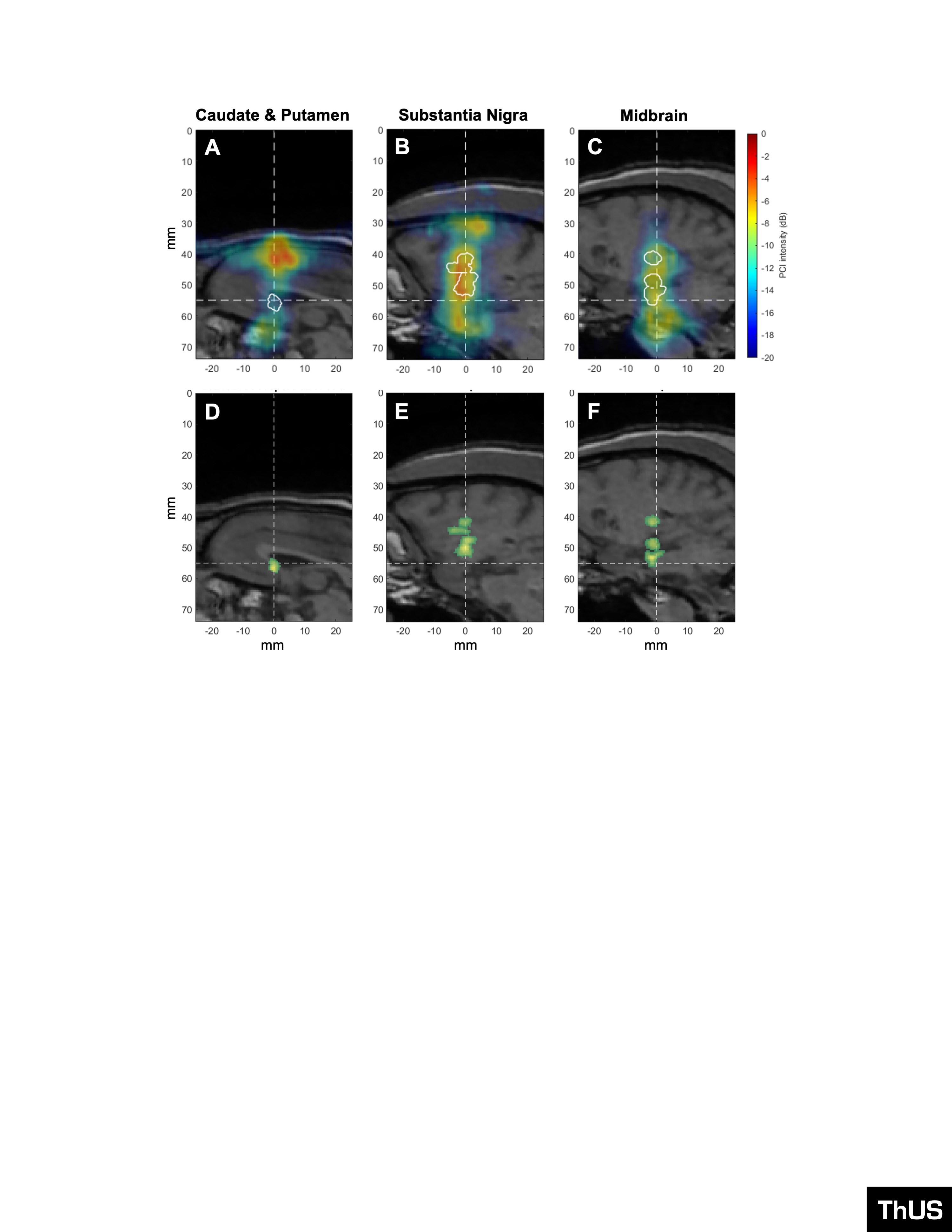

Figure 589: PCI and BBB opening volumes corresponding to targeted AAV9-CAG-GFP delivery with ThUS. PCI (colored overlays) and BBB opening contour (white outlines) superimposed onto central T1-weighted MRI slice after sonication with A) ThUS RASTA targeted to the caudate & putamen, B) a single ThUS focus targeted to the substantia nigra, and C) a single ThUS focus targeted to the midbrain. BBB opening volumes denoted by the green overlays for D) the caudate & putamen target (V_BBB_ = 19.61 mm3), E) substantia nigra target (V_BBB_ = 78.65 mm3), and F) the midbrain (V_BBB_ = 65.02 mm3). Dashed lines indicate the center of the planned ThUS focus after targeting. Note that the PCI and BBB opening shown in (A) and (D) correspond to the sonication depicted in Figure 2H-L.

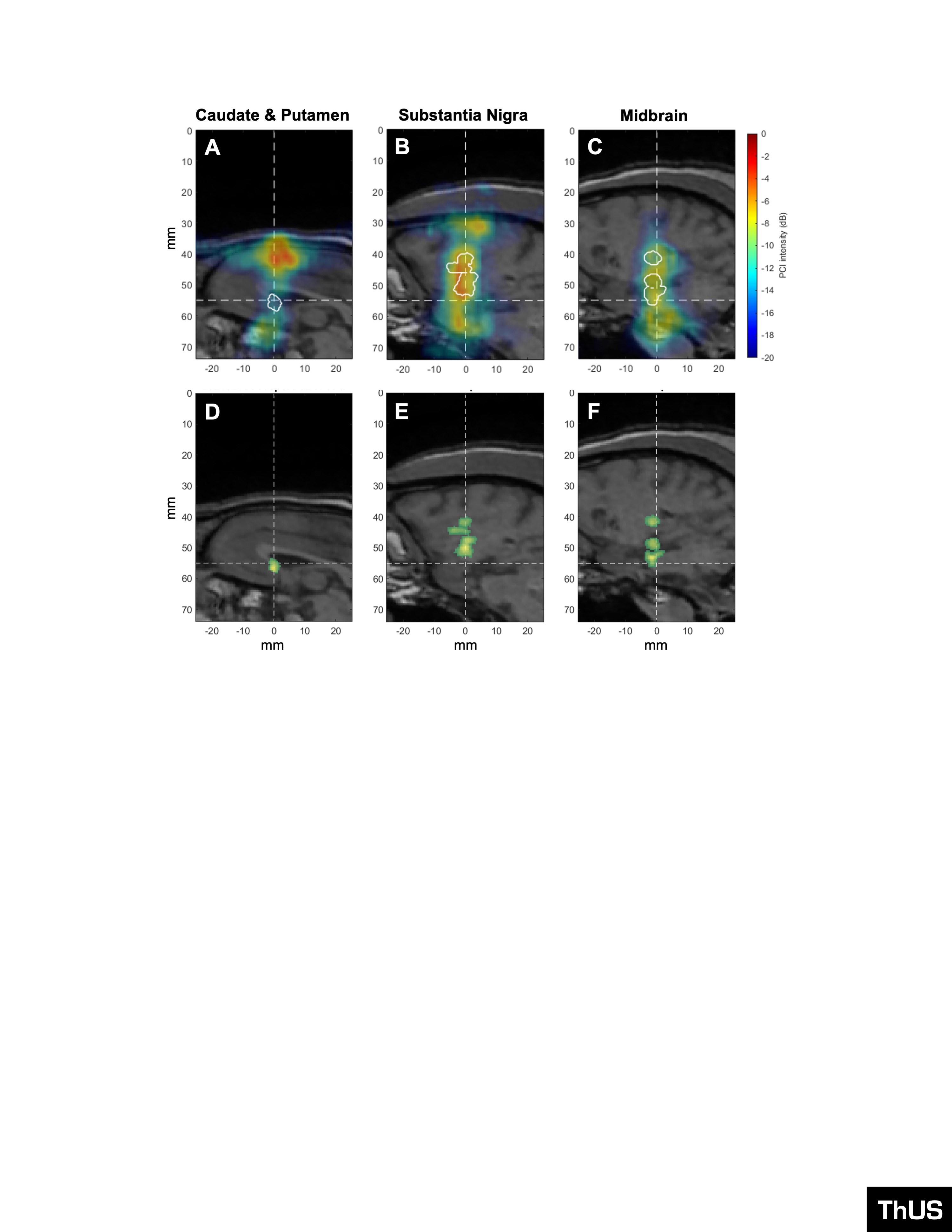

Figure 590: PCI and BBB opening volumes corresponding to targeted AAV9-CAG-GFP delivery with ThUS. PCI (colored overlays) and BBB opening contour (white outlines) superimposed onto central T1-weighted MRI slice after sonication with A) ThUS RASTA targeted to the caudate & putamen, B) a single ThUS focus targeted to the substantia nigra, and C) a single ThUS focus targeted to the midbrain. BBB opening volumes denoted by the green overlays for D) the caudate & putamen target (V_BBB_ = 19.61 mm3), E) substantia nigra target (V_BBB_ = 78.65 mm3), and F) the midbrain (V_BBB_ = 65.02 mm3). Dashed lines indicate the center of the planned ThUS focus after targeting. Note that the PCI and BBB opening shown in (A) and (D) correspond to the sonication depicted in Figure 2H-L.

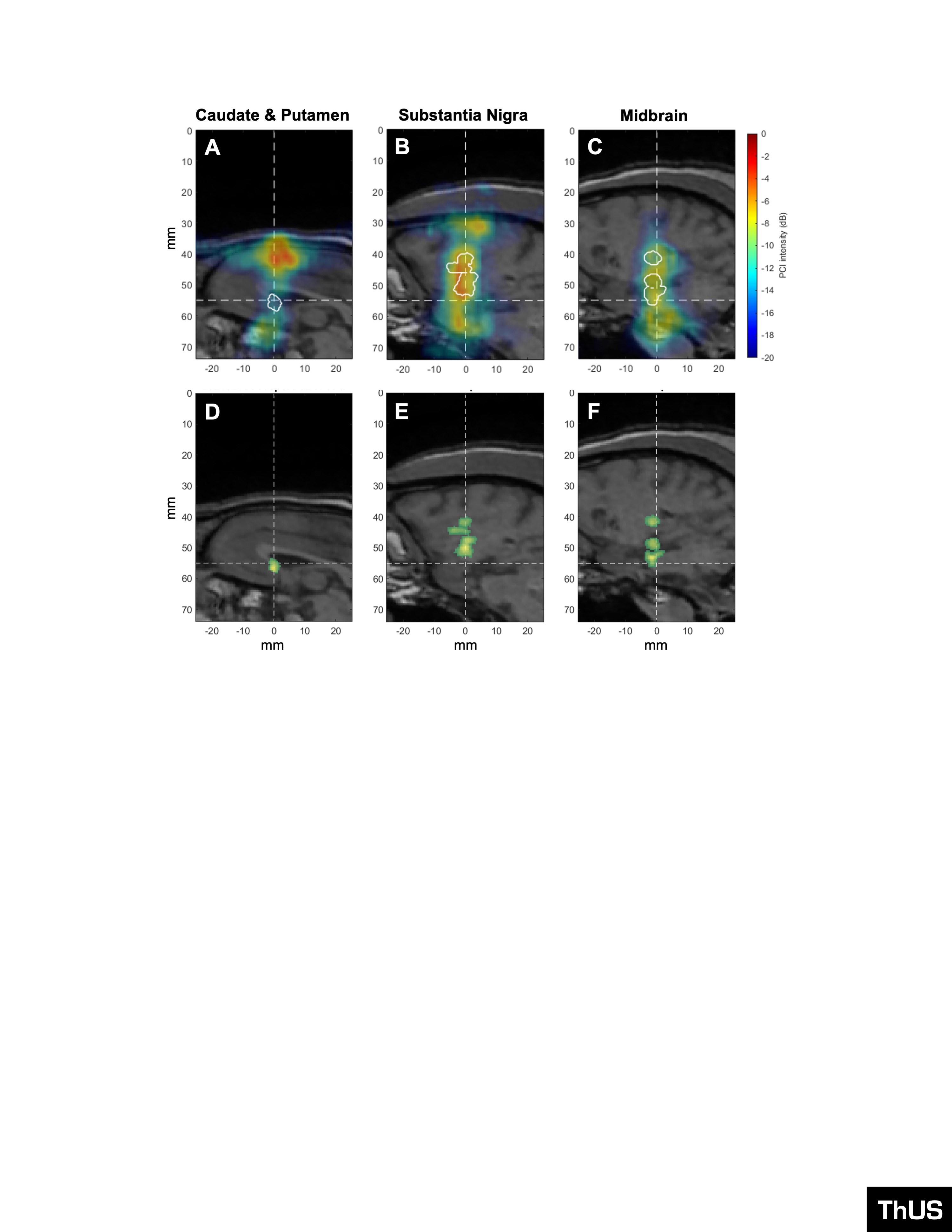

Figure 591: PCI and BBB opening volumes corresponding to targeted AAV9-CAG-GFP delivery with ThUS. PCI (colored overlays) and BBB opening contour (white outlines) superimposed onto central T1-weighted MRI slice after sonication with A) ThUS RASTA targeted to the caudate & putamen, B) a single ThUS focus targeted to the substantia nigra, and C) a single ThUS focus targeted to the midbrain. BBB opening volumes denoted by the green overlays for D) the caudate & putamen target (V_BBB_ = 19.61 mm3), E) substantia nigra target (V_BBB_ = 78.65 mm3), and F) the midbrain (V_BBB_ = 65.02 mm3). Dashed lines indicate the center of the planned ThUS focus after targeting. Note that the PCI and BBB opening shown in (A) and (D) correspond to the sonication depicted in Figure 2H-L.

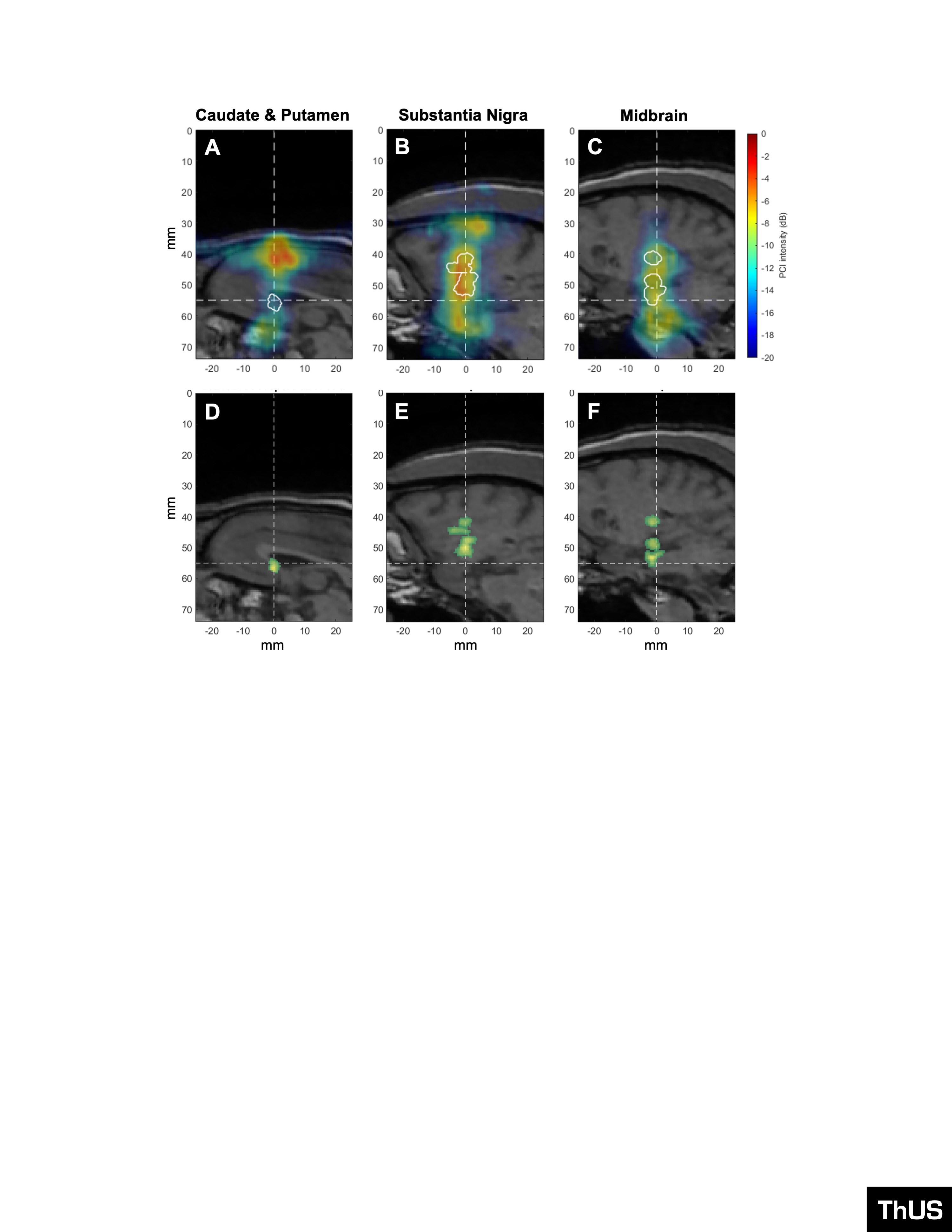

Figure 592: PCI and BBB opening volumes corresponding to targeted AAV9-CAG-GFP delivery with ThUS. PCI (colored overlays) and BBB opening contour (white outlines) superimposed onto central T1-weighted MRI slice after sonication with A) ThUS RASTA targeted to the caudate & putamen, B) a single ThUS focus targeted to the substantia nigra, and C) a single ThUS focus targeted to the midbrain. BBB opening volumes denoted by the green overlays for D) the caudate & putamen target (V_BBB_ = 19.61 mm3), E) substantia nigra target (V_BBB_ = 78.65 mm3), and F) the midbrain (V_BBB_ = 65.02 mm3). Dashed lines indicate the center of the planned ThUS focus after targeting. Note that the PCI and BBB opening shown in (A) and (D) correspond to the sonication depicted in Figure 2H-L.

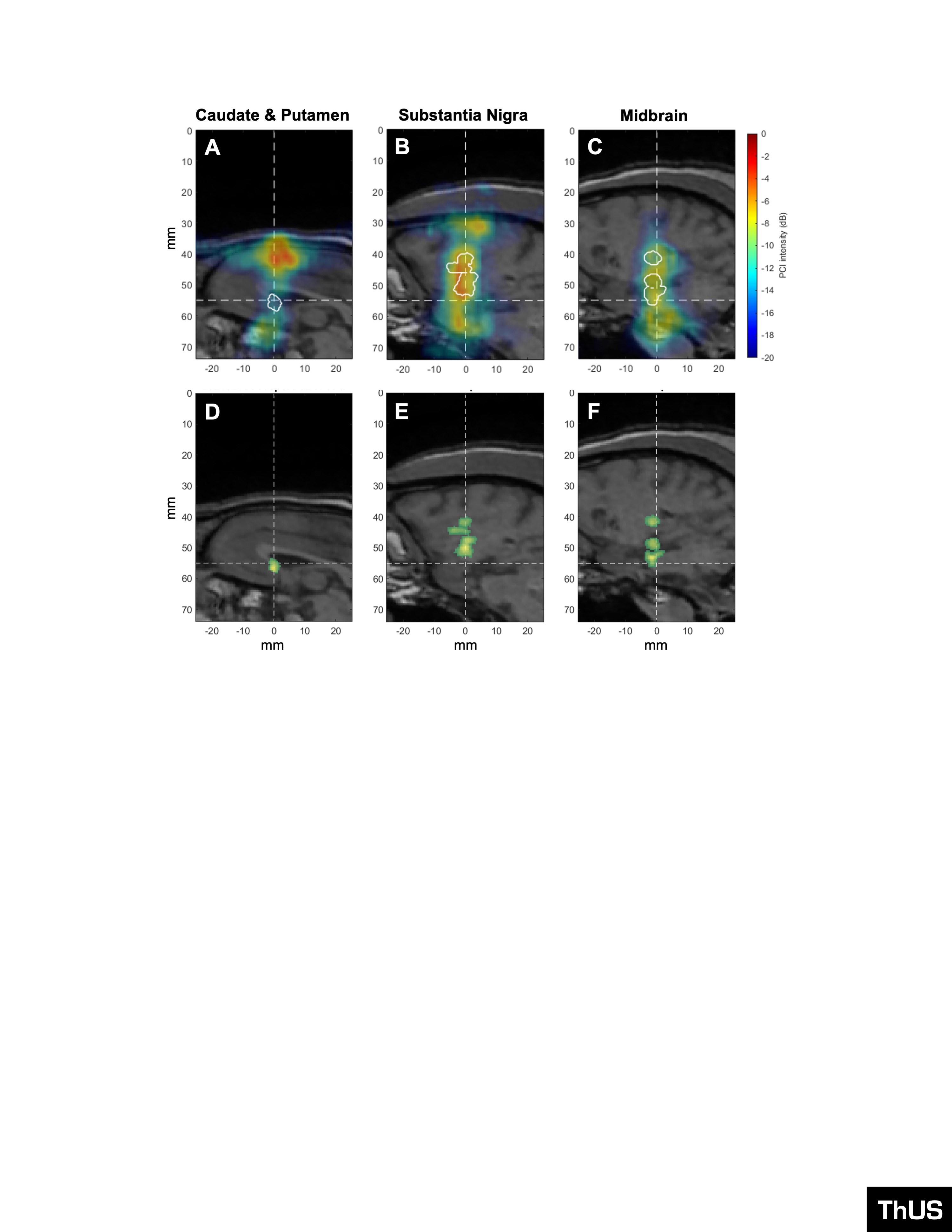

Figure 593: PCI and BBB opening volumes corresponding to targeted AAV9-CAG-GFP delivery with ThUS. PCI (colored overlays) and BBB opening contour (white outlines) superimposed onto central T1-weighted MRI slice after sonication with A) ThUS RASTA targeted to the caudate & putamen, B) a single ThUS focus targeted to the substantia nigra, and C) a single ThUS focus targeted to the midbrain. BBB opening volumes denoted by the green overlays for D) the caudate & putamen target (V_BBB_ = 19.61 mm3), E) substantia nigra target (V_BBB_ = 78.65 mm3), and F) the midbrain (V_BBB_ = 65.02 mm3). Dashed lines indicate the center of the planned ThUS focus after targeting. Note that the PCI and BBB opening shown in (A) and (D) correspond to the sonication depicted in Figure 2H-L.

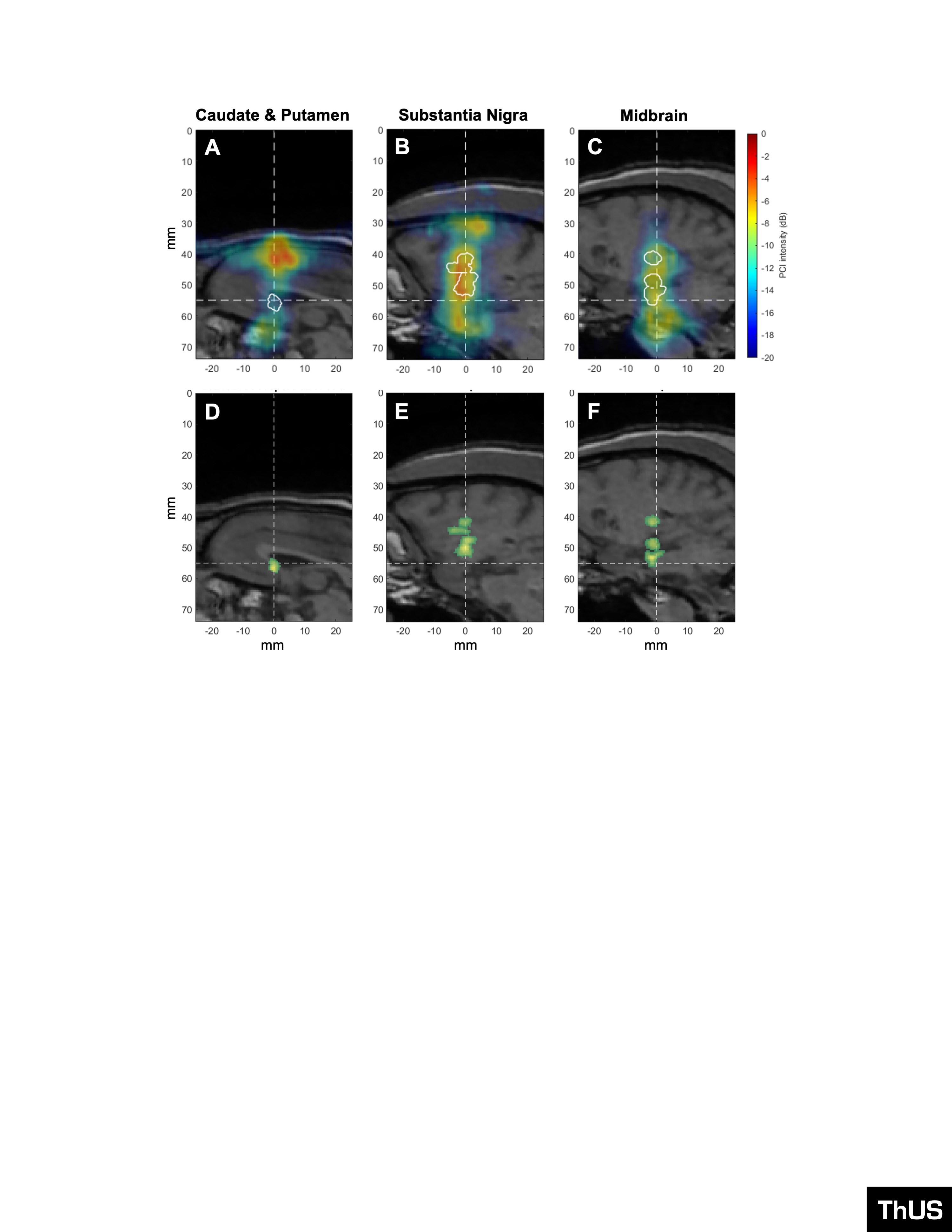

Figure 594: PCI and BBB opening volumes corresponding to targeted AAV9-CAG-GFP delivery with ThUS. PCI (colored overlays) and BBB opening contour (white outlines) superimposed onto central T1-weighted MRI slice after sonication with A) ThUS RASTA targeted to the caudate & putamen, B) a single ThUS focus targeted to the substantia nigra, and C) a single ThUS focus targeted to the midbrain. BBB opening volumes denoted by the green overlays for D) the caudate & putamen target (V_BBB_ = 19.61 mm3), E) substantia nigra target (V_BBB_ = 78.65 mm3), and F) the midbrain (V_BBB_ = 65.02 mm3). Dashed lines indicate the center of the planned ThUS focus after targeting. Note that the PCI and BBB opening shown in (A) and (D) correspond to the sonication depicted in Figure 2H-L.

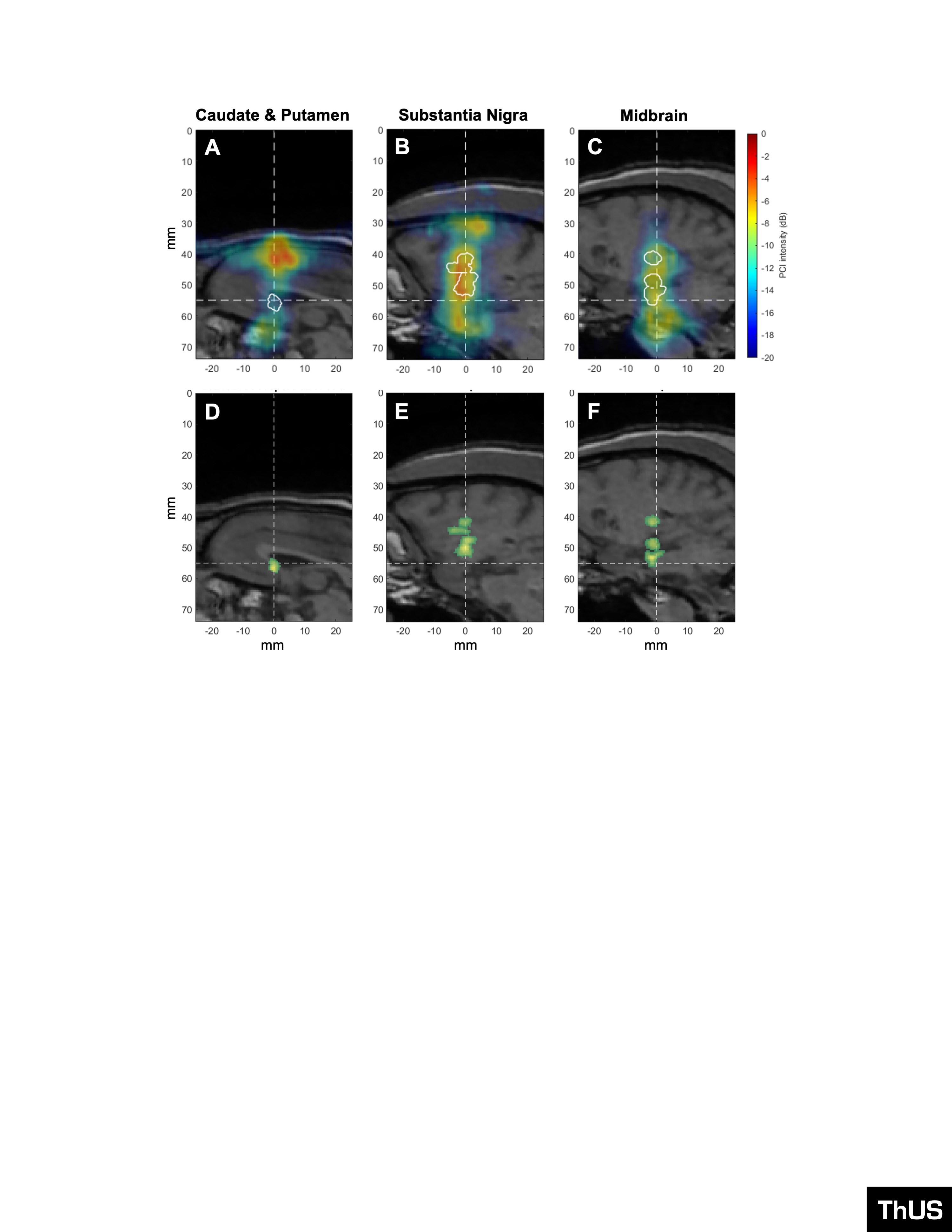

Figure 595: PCI and BBB opening volumes corresponding to targeted AAV9-CAG-GFP delivery with ThUS. PCI (colored overlays) and BBB opening contour (white outlines) superimposed onto central T1-weighted MRI slice after sonication with A) ThUS RASTA targeted to the caudate & putamen, B) a single ThUS focus targeted to the substantia nigra, and C) a single ThUS focus targeted to the midbrain. BBB opening volumes denoted by the green overlays for D) the caudate & putamen target (V_BBB_ = 19.61 mm3), E) substantia nigra target (V_BBB_ = 78.65 mm3), and F) the midbrain (V_BBB_ = 65.02 mm3). Dashed lines indicate the center of the planned ThUS focus after targeting. Note that the PCI and BBB opening shown in (A) and (D) correspond to the sonication depicted in Figure 2H-L.

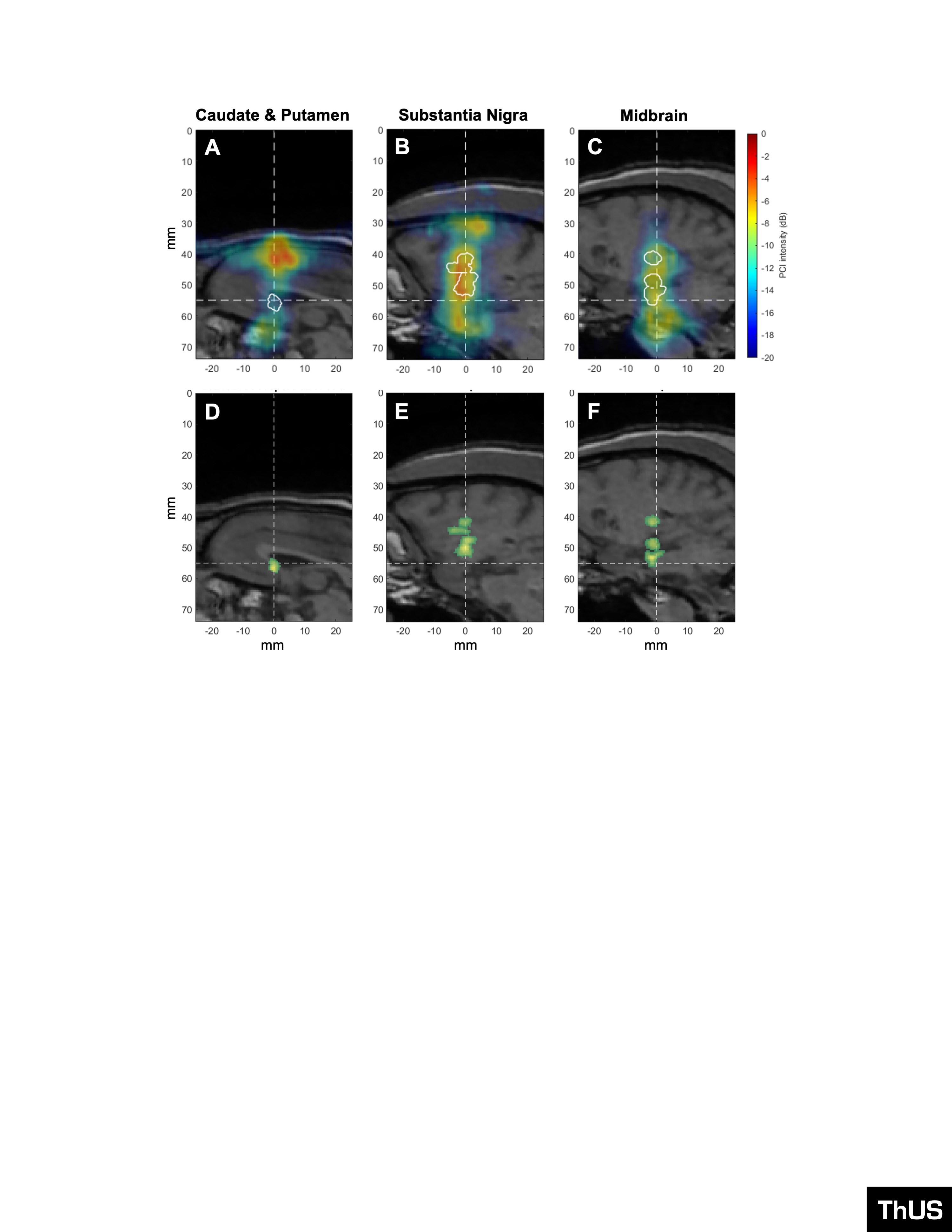

Figure 596: PCI and BBB opening volumes corresponding to targeted AAV9-CAG-GFP delivery with ThUS. PCI (colored overlays) and BBB opening contour (white outlines) superimposed onto central T1-weighted MRI slice after sonication with A) ThUS RASTA targeted to the caudate & putamen, B) a single ThUS focus targeted to the substantia nigra, and C) a single ThUS focus targeted to the midbrain. BBB opening volumes denoted by the green overlays for D) the caudate & putamen target (V_BBB_ = 19.61 mm3), E) substantia nigra target (V_BBB_ = 78.65 mm3), and F) the midbrain (V_BBB_ = 65.02 mm3). Dashed lines indicate the center of the planned ThUS focus after targeting. Note that the PCI and BBB opening shown in (A) and (D) correspond to the sonication depicted in Figure 2H-L.

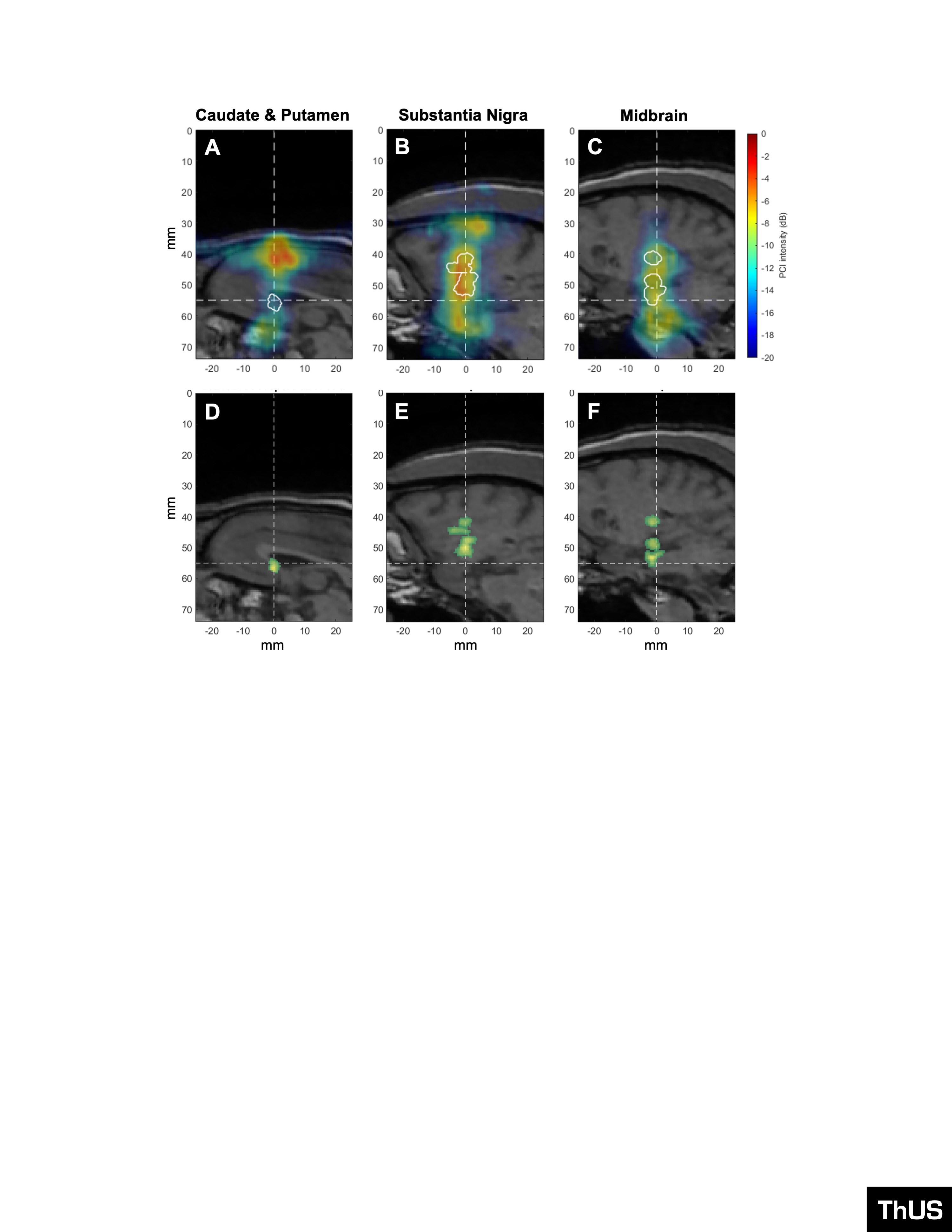

Figure 597: PCI and BBB opening volumes corresponding to targeted AAV9-CAG-GFP delivery with ThUS. PCI (colored overlays) and BBB opening contour (white outlines) superimposed onto central T1-weighted MRI slice after sonication with A) ThUS RASTA targeted to the caudate & putamen, B) a single ThUS focus targeted to the substantia nigra, and C) a single ThUS focus targeted to the midbrain. BBB opening volumes denoted by the green overlays for D) the caudate & putamen target (V_BBB_ = 19.61 mm3), E) substantia nigra target (V_BBB_ = 78.65 mm3), and F) the midbrain (V_BBB_ = 65.02 mm3). Dashed lines indicate the center of the planned ThUS focus after targeting. Note that the PCI and BBB opening shown in (A) and (D) correspond to the sonication depicted in Figure 2H-L.

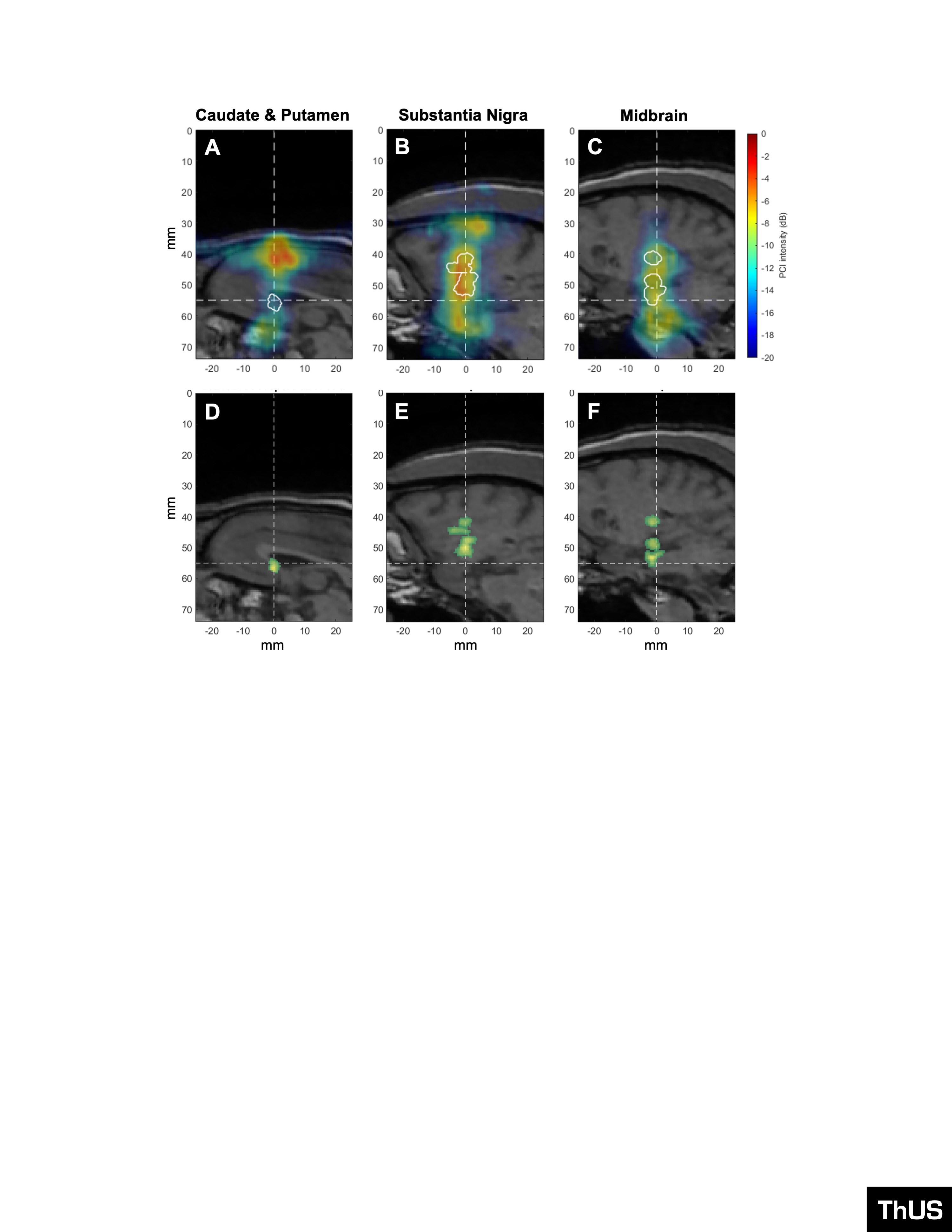

Figure 598: PCI and BBB opening volumes corresponding to targeted AAV9-CAG-GFP delivery with ThUS. PCI (colored overlays) and BBB opening contour (white outlines) superimposed onto central T1-weighted MRI slice after sonication with A) ThUS RASTA targeted to the caudate & putamen, B) a single ThUS focus targeted to the substantia nigra, and C) a single ThUS focus targeted to the midbrain. BBB opening volumes denoted by the green overlays for D) the caudate & putamen target (V_BBB_ = 19.61 mm3), E) substantia nigra target (V_BBB_ = 78.65 mm3), and F) the midbrain (V_BBB_ = 65.02 mm3). Dashed lines indicate the center of the planned ThUS focus after targeting. Note that the PCI and BBB opening shown in (A) and (D) correspond to the sonication depicted in Figure 2H-L.

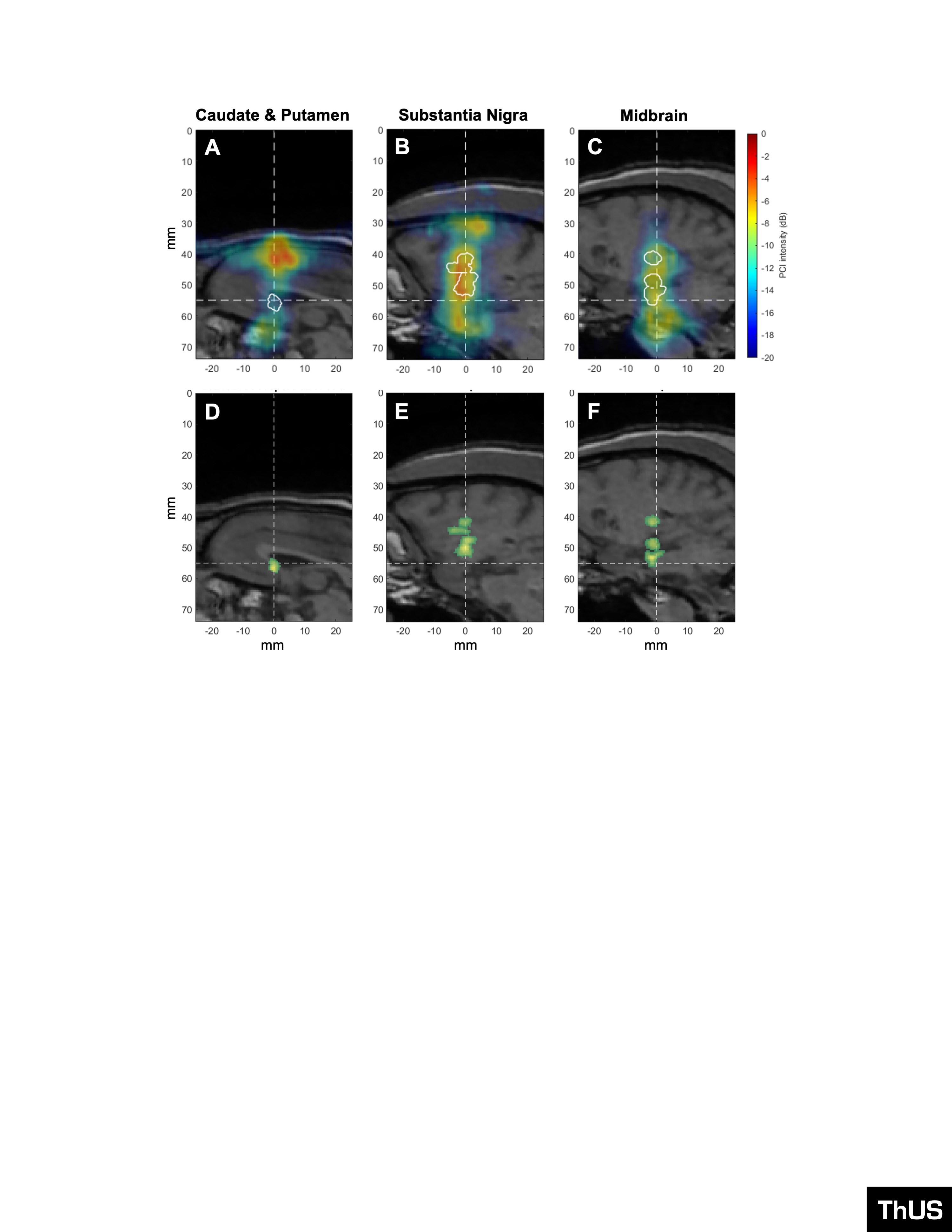

Figure 599: PCI and BBB opening volumes corresponding to targeted AAV9-CAG-GFP delivery with ThUS. PCI (colored overlays) and BBB opening contour (white outlines) superimposed onto central T1-weighted MRI slice after sonication with A) ThUS RASTA targeted to the caudate & putamen, B) a single ThUS focus targeted to the substantia nigra, and C) a single ThUS focus targeted to the midbrain. BBB opening volumes denoted by the green overlays for D) the caudate & putamen target (V_BBB_ = 19.61 mm3), E) substantia nigra target (V_BBB_ = 78.65 mm3), and F) the midbrain (V_BBB_ = 65.02 mm3). Dashed lines indicate the center of the planned ThUS focus after targeting. Note that the PCI and BBB opening shown in (A) and (D) correspond to the sonication depicted in Figure 2H-L.

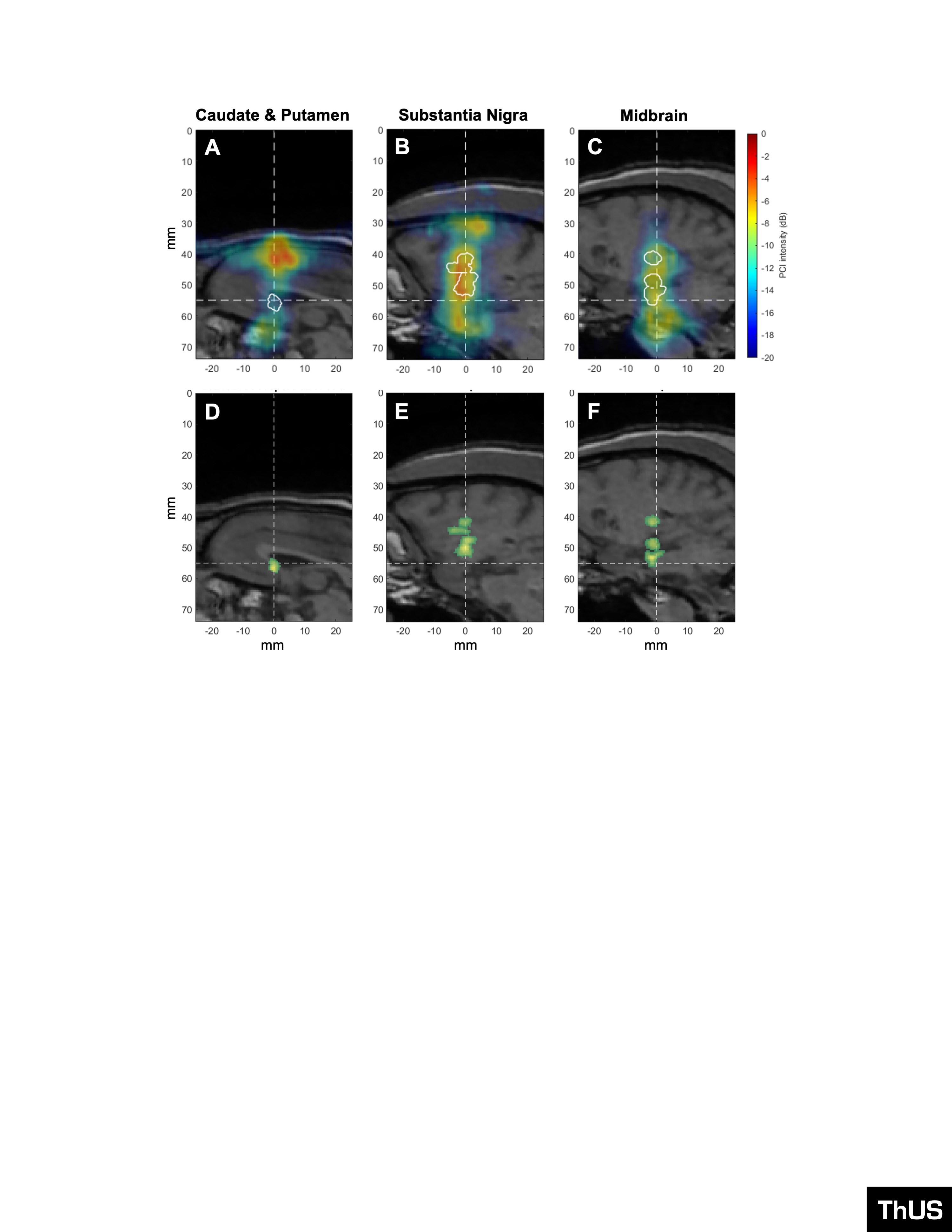

Figure 600: PCI and BBB opening volumes corresponding to targeted AAV9-CAG-GFP delivery with ThUS. PCI (colored overlays) and BBB opening contour (white outlines) superimposed onto central T1-weighted MRI slice after sonication with A) ThUS RASTA targeted to the caudate & putamen, B) a single ThUS focus targeted to the substantia nigra, and C) a single ThUS focus targeted to the midbrain. BBB opening volumes denoted by the green overlays for D) the caudate & putamen target (V_BBB_ = 19.61 mm3), E) substantia nigra target (V_BBB_ = 78.65 mm3), and F) the midbrain (V_BBB_ = 65.02 mm3). Dashed lines indicate the center of the planned ThUS focus after targeting. Note that the PCI and BBB opening shown in (A) and (D) correspond to the sonication depicted in Figure 2H-L.

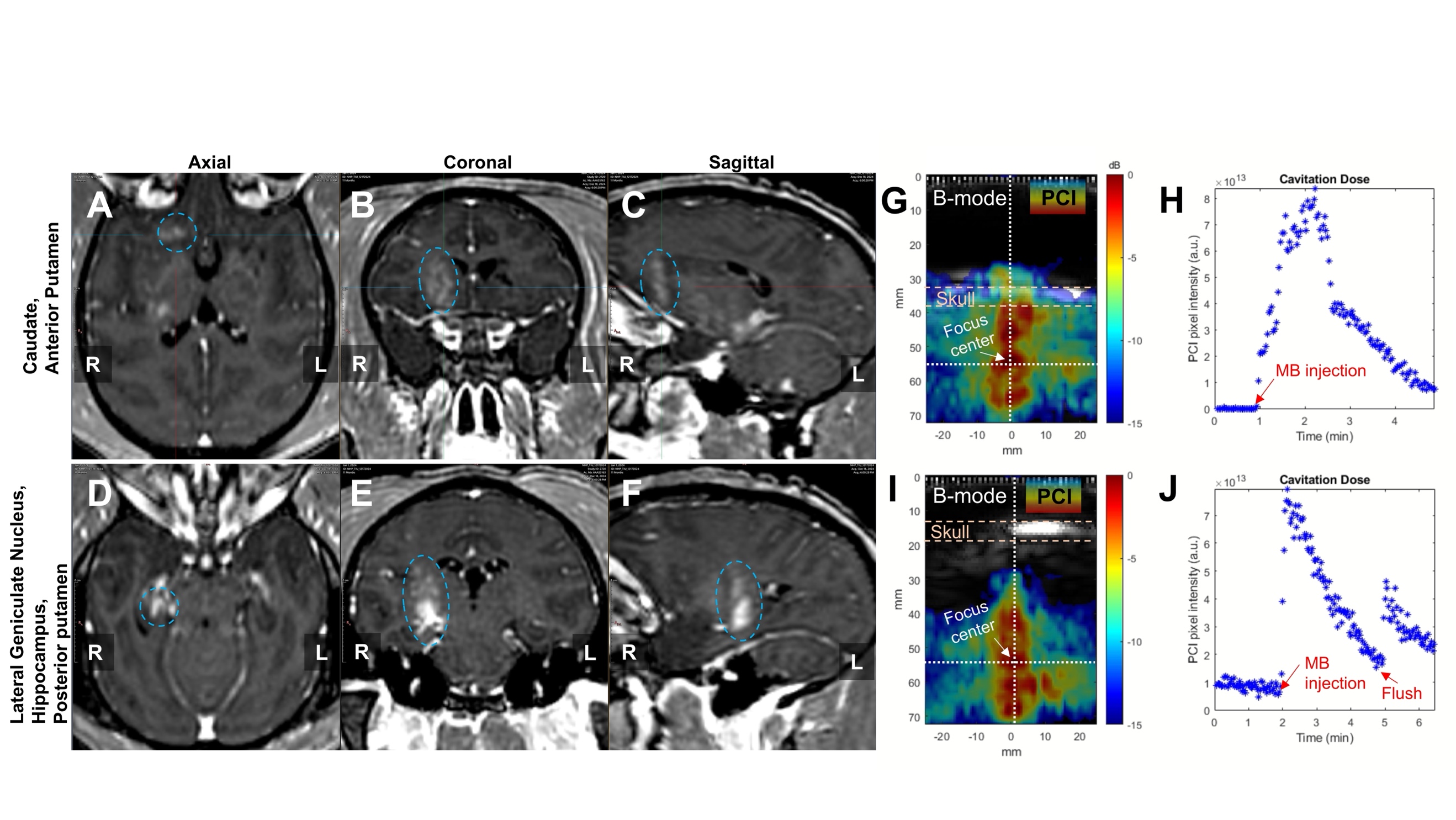

**Supplementary Figure 8: ThUS-mediated BBB opening and PCI with LUMASON microbubbles. A)** Axial, **B)** coronal, and **C)** sagittal contrast-enhanced T1-w MRI depicting BBB opening in the caudate and anterior putamen 1 hour post-sonication. **D)** Axial, **E)** coronal, and **F)** sagittal contrast-enhanced T1-w MRI depicting BBB opening in the LGN, hippocampus and posterior putamen 1 hour post-sonication. Blue ellipses denote approximate ROI depicting BBB opening. **G)** Representative 1^st^ sonication PCI overlaid onto pre-sonication B-mode image depicting localized cavitation activity. **H)** Cavitation dose over the 1^st^ sonication duration. **I)** Representative 2^nd^ sonication PCI overlaid onto pre-sonication B-mode image. **J)** Cavitation dose over the 2^nd^ sonication duration. Note the greater initial (baseline) cavitation dose from 0-2 min prior to the 2^nd^ bolus injection due to microbubbles still circulating from the 1^st^ injection.

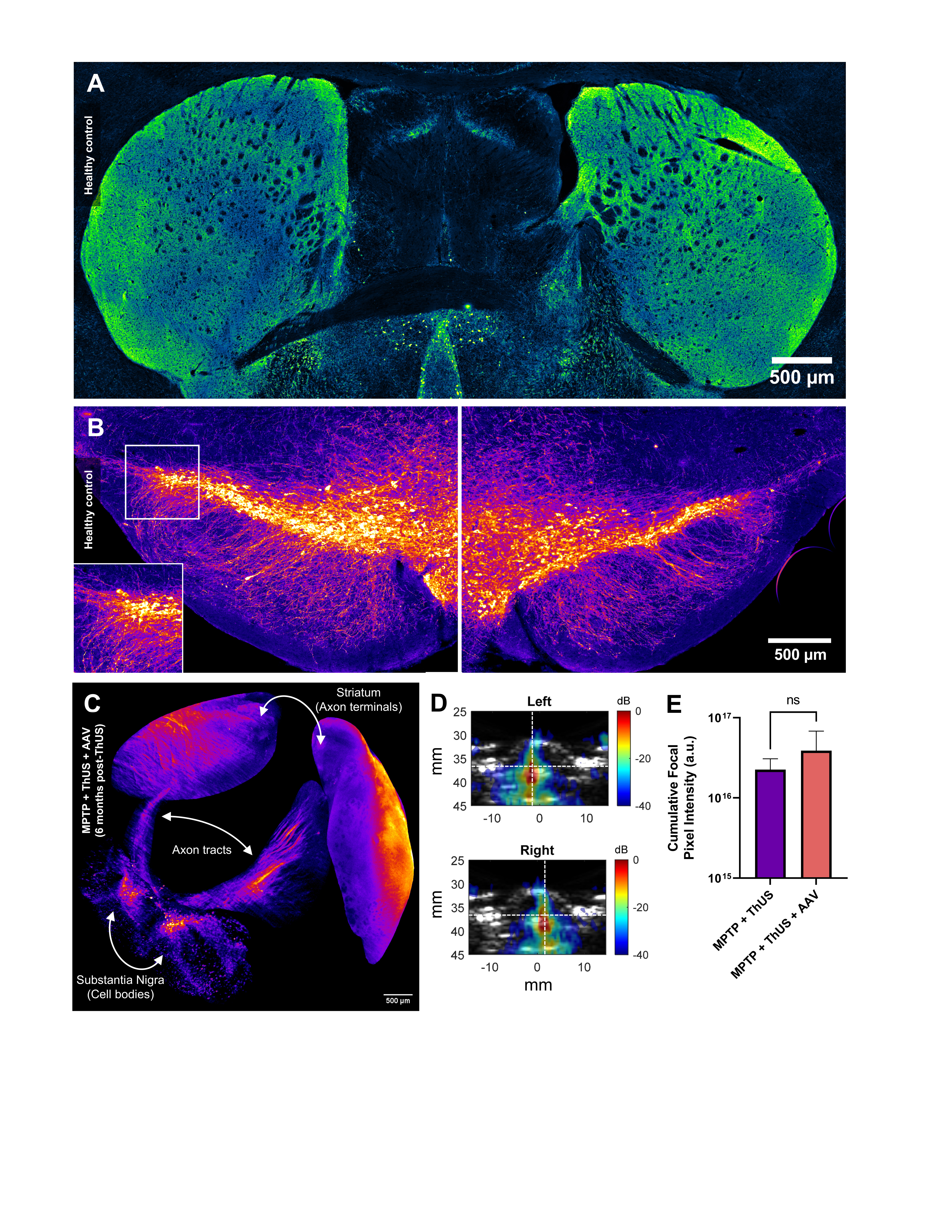

**Supplementary Figure 9: Additional MPTP histology and ThUS cavitation mapping. A)** Pseudo-colored TH immunofluorescence image of striatum from healthy control mouse not receiving MPTP. **B)** Pseudo-colored TH immunofluorescence image of substantia nigra from healthy control mouse not receiving MPTP. **C)** 3D reconstructed light-sheet microscopy image of TH immunolabeled and iDISCO tissue-cleared brain of an MPTP mouse euthanized 6 months after receiving ThUS+AAV treatment. **D)** Representative cavitation images from MPTP mice treated with ThUS+AAV. **E)** No significant differences in average cavitation doses across ThUS treatment groups determined with unpaired t-test.

**Supplementary Tables:**

**Supplementary Table 1: FUS + PAM device specifications and parameters**

| FUS transducer | H-231 (Sonic Concepts) |
| --- | --- |
| Center frequency | 0.25 MHz |
| Elements | 1 |
| Outer/inner diameter | 110/44 mm |
| Focal volume | 6 mm x 6 mm x 49 mm |
| Pulse length | 10 ms |
| Pulse repetition frequency | 2 Hz |
| Sonication duration | 2 min |
| Derated pressure | 0.4 MPa |
| Mechanical index | 0.8 |
| Imaging Array | P4-1 (ATL, Philips) |
| Center frequency | 2.5 MHz |
| Elements | 96 |
| Aperture | 19.2 mm |
| Sampling frequency | 10 MHz |
| Microbubbles | House-made polydisperse |
| Microbubble dose | 8.0e8 MBs/mL (10X clinical Definity® dose) |

**Supplementary Table 2: ThUS device specifications and parameters**

| ThUS transducer | 500 kHz linear array (Vermon) |
| --- | --- |
| Focal volume | 4 mm x 16 mm x 40 mm |
| Center frequency | 0.5 MHz |
| Elements | 32 |
| Pitch | 1.6 mm |
| Kerf | 0.1 mm |
| Elevational aperture | 25 mm |
| Acoustic lens thickness | 0.4 mm |
| Elevational plane focal distance | 55 mm |
| f-number | 1.07 |
| Pulse length | 3 cycles, or 5 cycles (NHP C) |
| Pulse repetition frequency (PRF) | 1000 Hz |
| Burst repetition frequency (BRF) | 0.5 Hz |
| Derated pressure | 1.0 MPa |
| Mechanical Index | 1.4 |
| Sonication Duration | 4 min for RASTA, 2 min/target otherwise |
| Microbubbles | House-made polydisperse |
| Microbubble dose | 8.0e8 MBs/mL (10X clinical Definity® dose) |

**Supplementary Video Captions:**

**Video 1: 3D rendering of BBB opening volumes in NHP A.** Green regions indicate contrast-enhanced volume on post-FUS T_1_-weighted MRI, while blue lines and blue centroids indicate the FUS trajectory and center of the FUS focus, respectively. **Filename: NHPA_BBBO_FUS.mp4**

**Video 2: 3D rendering of BBB opening volumes in NHP B.** Green regions indicate contrast-enhanced volume on post-FUS T_1_-weighted MRI, while blue lines and blue centroids indicate the ThUS trajectory and center of the ThUS focus, respectively. Only the 3 ThUS sonications are displayed for clarity. The single FUS sonication in NHP B is not displayed in this rendering. **Filename: NHPB_BBBO_ThUS.mp4**
